# Histologically Informed Multiscale Modeling of the Neuronal Elements Activated by TMS

**DOI:** 10.64898/2026.06.05.730288

**Authors:** Torge H. Worbs, Jesper D. Nielsen, Boshuo Wang, John W. Hansen, Warren M. Grill, Angel V. Peterchev, Axel Thielscher

## Abstract

**Background:** The primary neural site(s) at which action potentials are initiated by transcranial magnetic stimulation (TMS) remain poorly understood. Multiscale computational models provide biophysically based hypotheses, but model accuracy is constrained by limited histological knowledge of the microscopic organization of neural tissue. Recent high-resolution electron microscopy, in particular the petavoxel H01 dataset, provides novel, detailed axon morphologies and myelination patterns within the human cortex and superficial white matter.

**Objective:** To compare systematically multiple candidates for the neural elements activated by TMS using computer simulations informed by an extensive body of histological measurements, including neuron models directly reconstructed from the H01 dataset.

**Methods:** We developed a novel modeling pipeline to extract individual morphologically realistic multi-compartment models with exact myelination from serial section electron microscopic segmentations. To assess candidate excitation sites, we simulated the extracted neuron models together with parameterized models of a “ball-and-two-sticks”, bifurcation, termination, and bend under uniform electric fields. In addition, smooth and sharply bending myelinated axons were embedded in a realistic human head model to evaluate activation thresholds under anatomically realistic electric field distributions.

**Results:** Axon terminations were only excitable by TMS when they were fully myelinated, which the histology suggested is unlikely. Partial myelination, even when separated by only 10 µm from the terminal, increased activation thresholds by more than 100%. Reconstructed H01 neurons exhibited correspondingly high activation thresholds at axon terminals due to a lack of myelination. Further, most other candidate structures exhibited low thresholds only for histologically unrealistic parameter choices. In contrast, myelinated axonal bends of fibers transitioning from cortex to superficial white matter consistently showed low activation thresholds for both uniform electric fields and in realistic head model simulations. These thresholds fell within physiologically realistic ranges and, for larger diameter fibers, approached experimentally measured motor thresholds.

**Conclusion:** These results identify myelinated axons bending from cortex into superficial white matter as possible neural targets for transcranial magnetic stimulation, and demonstrate the relevance of detailed histological and biophysical information to support robust modeling results.

## Introduction

Transcranial magnetic stimulation (TMS) uses a rapidly changing magnetic field to induce an electric field (E-field) in the brain that can polarize neurons. In addition to its use in basic neuroscience, TMS is approved by major regulatory agencies as a treatment for several psychiatric disorders, including major depressive disorder and obsessive-compulsive disorder [1]. Despite its widespread clinical and experimental use, the understanding of its neural stimulation mechanisms remains limited, including the question of the primary neural elements where action potentials are initiated [2].

Computational models offer a promising approach to quantify how neuronal activation depends on parameters such as E-field waveform and orientation. Several studies modeled TMS-induced activation in cortical neurons [3–6], but model accuracy was constrained by limited histological data on realistic axonal trajectories, diameters, and myelination patterns. Recently, high-resolution serial section electron microscopy (EM) has enabled dense reconstructions of large portions of cortical neurons [7–9], providing unprecedented morphological detail across all six cortical layers and part of the superficial white matter (SWM). In this work, we reconstructed human cortical neurons with exact myelination from the H01 dataset and simulated their responses to uniform E-fields. We found vastly different activation thresholds than those reported in previous studies. This led us to conduct a comprehensive investigation of multiple different neural elements to reach unbiased conclusions about the putative activation mechanisms of TMS.

Axon bends are possible targets of TMS. They occur intracortically along local and afferent axons within gray matter (GM) as well as projection fibers traversing from the cortex into superficial and deep white matter (WM) and *vice versa*. Earlier work focused primarily on smoothly curved long-range fibers projecting into deep WM and examined only a narrow set of geometric parameters [10]. Our study additionally includes short-range SWM fibers, leveraging newly available histological data. SWM tracts are widespread across the human cortex and are both myelinated and densely organized. They are also estimated to outnumber deep WM fibers by an order of magnitude [11]. This dense, locally oriented architecture makes axons bending into SWM a potential target for TMS-induced activation.

We compare multiple types of neural elements relevant to TMS-induced excitation, including realistic human neuronal morphologies, simplified models capturing axonal discontinuities (axon initial segment (AIS), bends, terminations, branches) as well as short- and long-range connections in both superficial and deep WM. These models incorporate detailed morphological features derived from recent high-resolution EM datasets and an extensive body of histological studies and provide a biologically grounded basis for evaluating TMS activation.

## Methods

This study integrates four complementary modeling approaches to identify the neuronal structures most likely to initiate activation during TMS. First, we quantified the responses of morphologically and biophysically detailed human neuron models to a uniform E-field of varying orientations. We then studied simplified representations of axonal discontinuities and introduced histologically grounded variability in their key morphological parameters to systematically assess how structural features influence excitability. Based on these analyses, we identified axonal bends as most plausible sites of activation initiation and, in a final substudy, incorporated both smooth and sharp bend geometries into a realistic head model to compare their activation thresholds with experimentally determined motor thresholds (MTs).

The biophysical parameters used in the neuronal compartment models were adapted from models of somatosensory cortical neurons of P14 male Wistar (Han) rats, as previously described by Aberra et al. [12–14]. We implemented a revised myelin model to more accurately incorporate experimentally measured internodal specific capacitance and membrane leakage conductance. A detailed description of the biophysical models, their parameter values, and our adaptations is provided in Supplementary Material A. We validated the axon model by comparing simulated and experimentally measured conduction velocities, confirming the velocity distributions to largely overlap (Supplementary Material B). The specific biophysical models used for each compartment model are detailed in the following sections.

### Simulations of Neuronal Excitability in Uniform Electric Field

#### Simulation of Biophysically Realistic Neuronal Compartment Models in NEURON

The simulations of neuronal compartment models under a uniform E-field were performed using the simulation environment NEURON 8.2 [15]. The field was numerically integrated along each neurite and the extracellular quasipotential was used as the extracellular potential in NEURON [16,6]. If not described otherwise, a monophasic TMS pulse waveform generated by a MagPro X100 stimulator (MagVenture A/S, Denmark) [6] was used to estimate the thresholds of the neuronal morphologies. Simulations were performed at a temperature of 37°C, solved using the backward Euler technique with a time step of 5 µs, with compartments no longer than 5 µm and a binary threshold search accuracy of 0.01%. The simulation was started from a steady state where the membrane potential of all compartments was allowed to equilibrate.

#### Construction and Simulation of Morphologically and Biophysically Detailed Human Cortical Neuron Models

We selected two pyramidal neurons per layer from layer II, III, IV, V and VI, included in the H01 dataset [9]. The H01 dataset provides high-resolution serial EM images of a 1 mm³ tissue sample from the left human anterior middle temporal gyrus spanning the cortex and part of the underlying gyral WM. Neurons in each cortical layer were selected based on two criteria: (1) they represented typical morphological features of the local neuronal population, and (2) a substantial portion of their axonal and dendritic arborizations was contained within the imaged volume. Neuron proofreading was performed using the online visualization platform Neuroglancer, and morphologically detailed cable models were reconstructed through a newly developed pipeline described in Supplementary Material C. All reconstructed neurons were manually validated for morphological accuracy against the original EM images. The specific biophysical models used for the extracted neurons are listed in Supplementary Table S-C1. On average, reconstruction took 18 person-hours per neuron using the semiautomatic pipeline. No general correction was applied for shrinkage due to the chemical fixation process in the H01 dataset. However, as the dehydration process used in the tissue preparation has been shown to introduce a diameter-dependent axon shrinkage, all axonal diameters were uniformly scaled by a factor of 1.59 as an approximate correction, based on published measurements in mouse cortical tissue [17]. The correction factor was a pragmatic choice in the absence of corresponding human calibration data. We then simulated each extracted neuron model under 648 spherically sampled E-field directions and identified the most excitable target location.

#### Geometric Modeling and Simulation of Synthetic Neuronal Morphologies

We constructed synthetic neuronal morphologies to study in detail four axonal geometric discontinuities that might be potentially relevant for TMS stimulation: Diameter tapering at the AIS, bifurcations, terminations, and bends. All compartments (apical dendrite, soma, AIS, axon, node of Ranvier, myelinated axon, and axon terminal) were assigned biophysical properties consistent with the altered layer V pyramidal neuron models (“L5_TTPC2_cADpyr”, for details, see Supplementary Material A). The choices of all morphological parameter ranges were based on histological measurements from literature summarized in Supplementary Material D.

For each model, activation thresholds were computed in response to a uniform E-field aligned to minimize the threshold. The optimal field direction was verified by rotating the E-field within the plane of the model and evaluating threshold changes (Supplementary Fig. S3), confirming that the threshold increased with angular offset according to the expected cosine dependence

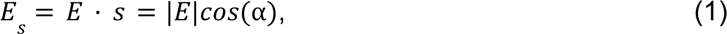

which reflects the projection of the E-field *E* onto each segment with direction indicated by unit vector *s* and angle α between *E* and *s*. Time constants and rheobase values were computed for a selected subset of parameter combinations and are discussed in Supplementary Material E.

### Ball-and-Two-Sticks

The ball-and-two-sticks model (Fig. 2A) was designed to study the putative excitability of the AIS by TMS due to diameter tapering. The morphological model consisted of a spherical soma, a straight apical dendrite emerging from the soma apex, and a straight axon extending from the base. The soma diameter *S* was varied between 10 and 100 µm, together with the axon diameter *D* between 0.2 and 10 µm, with a fixed ratio between the two. The sizes approximately spanned the reported range from small pyramidal cells up to the largest Betz cells. We tested AIS lengths *K* ranging from 10 to 80 µm and soma to AIS gap lengths *G* from 0 to 45 µm. Within the first 70 µm of the axon, the diameter tapered linearly from twice its final value *I* to its final diameter [18]. The axon maintained its constant diameter over the remaining 2 mm. The final 1 mm of the axon lacked active channels, preventing terminal activation. The apical dendrite tapered quadratically, which is theoretically optimal for current transfer in dendrites, starting at half the soma’s maximum diameter and ending with a diameter of 1 µm [18,19]. The applied uniform E-field was aligned with the somatodendritic axis and oriented toward the axon.

### Bifurcation

The second model (Fig. 2B) represented an idealized bifurcation of a myelinated axon with a node of Ranvier located at the branch point. An incoming axon split into two collaterals of equal diameter. Considering the variability in the literature concerning diameter changes at the bifurcation, we used an idealized case for this model where the incoming and the two outgoing axons had the same diameter, which maximized excitability while still capturing key characteristics of realistic bifurcations. For cases where one of the collaterals had a considerably smaller diameter than the other two, the model approached the result of a bend. Likewise, for cases where two of the branches had far smaller diameters, the results approach the terminal case. Both cases are discussed later and therefore not be included here. We varied the distance from the bifurcation to the start of the first internode *M* of both collaterals between 0 and 100 µm. The angle between the two collaterals *β* was considered in the range from 10° to 90° and the axon diameters *D* ranged from 0.2–10 µm. For myelination, we considered ratios of internode length *L* to *D* (*L*/*D*) ranging from 60 to 120 and node of Ranvier lengths from 0.5 to 4 µm. All three branches were myelinated for at least 1 mm and had at least five internodes, followed by a 1 mm unmyelinated segment where active channels were removed to prevent terminal activation. The E-field was aligned toward the bifurcation, bisecting the angle between the two collaterals.

### Termination

For the model of cortical axon terminals (Fig. 2C), the geometry was defined by a straight myelinated axon that terminated with an unmyelinated segment. We varied the axon diameter *D* between 0.1 and 1 µm and the myelin distance to the terminal *M* in the range of 0–100 µm. We modeled internode length *L* in the range of 15–100 µm and node of Ranvier lengths *N* from 0.5 to 4 µm. The terminal was modeled with a length of 1 µm. Before the termination, the axon was myelinated for at least 1 mm and at least five internodes, followed by a 1 mm unmyelinated segment where active channels were removed to prevent terminal activation. The E-field was aligned parallel with the axon and directed toward the termination.

### Bend

The final model represented a myelinated axon bend (Fig. 2D), where the myelination pattern was arranged so that a node of Ranvier lay at the midpoint of the bend. We varied the bend radius *R* in the range of 1–1000 µm and the bend angle *β* in the range of 45°–180°. The diameter *D* was varied between 0.2 and 10 µm. For myelination, we considered *L*/*D* ratios ranging from 60 to 120 and node of Ranvier lengths *N* from 0.5 to 4 µm. Each side of the bend was followed by at least 1 mm of myelinated axon and 1 mm of passive unmyelinated axon to avoid terminal activation. The applied E-field was oriented toward the bend, bisecting the bending angle. As bends in general exhibited a high sensitivity to TMS stimulation, we studied the dependence of their activation thresholds on morphological and biophysical parameters in more detail, including node of Ranvier diameter, the ratio of the inner axonal diameter to the outer diameter of the myelin sheath (g-ratio), sodium channel density, internode count, and node offset relative to the bend center.

#### Geometric Modeling and Simulation of Synthetic Axon Morphologies in the Human Motor Cortex

To further characterize the excitability of myelinated axon bends by TMS, we integrated them at various positions into GM and WM of a realistic head model to simulate activation thresholds under realistic E-field distributions. The following paragraphs describe the experimental data used to build the head model, the field simulations and the geometric modeling of the axon bends.

#### TMS Experiment

Previously acquired data of one healthy, right-handed participant was used to inform multiscale simulations of bending axons embedded in the handknob region of the human precentral gyrus (PreG-HAND). For details of the original study, please see [20]. In brief, neuronavigated TMS with monophasic pulses was applied using a MagPro X100 stimulator with a standard figure-8 coil (MC-B70). In an initial session, the motor hotspot for the first dorsal interosseous (FDI) muscle was determined by a combination of systematic sampling at a grid of positions and E-field simulations. Afterwards, input–output curves for 9 different coil orientations above the motor hotspot were acquired during controlled muscle preactivation. The active MT was then estimated from sigmoidal fits applied to the input–output curves.

#### Electric Field Modeling

An individual head model (Fig. 5A) of the participant was generated using CHARM [21] (SimNIBS 4.6 [22]) from anonymized versions of the structural T1- and T2-weighted and diffusion magnetic resonance images (MRI) acquired in the original study [20]. We defined a region of interest (ROI) covering PreG-HAND (Fig. 5E) and meshed it with a high-resolution element size of 0.4 mm (Fig. 5B). The head model contains nine different tissues. Isotropic conductivities were used for the cerebrospinal fluid (1.654 S/m), scalp (0.465 S/m), eye balls (0.5 S/m), compact bone (0.008 S/m), spongy bone (0.025 S/m), blood (0.6 S/m), and muscle (0.16 S/m). The WM anisotropic conductivity was reconstructed from the diffusion MRIs using the volume-normalized mapping approach [23] implemented in SimNIBS 4.6 (dwi2cond). The GM conductivity was set to isotropic tensors with a magnitude of 0.275 S/m. The conductivity at the GM/WM boundary was smoothed to create a graded transition zone between the gray and WM conductivities. Based on the concentration of neuron somata in the H01 dataset, we estimated that the transition zone spans from the center of layer VI to the WM surface and by the same distance further into WM [9]. In the transition zone, both GM and WM were anisotropic with equivalent isotropic conductivities that became inhomogeneous due to the smoothing. Details about the smoothing procedure and a comparison to isotropic conductivities can be found in Supplementary Material F.

The simulation was performed using a model of the MC-B70 coil [24] and for the optimal orientation estimated from the nine coil orientations tested in the original TMS experiment (Figs. 5E & 5C). For the analysis of activation thresholds, we defined three cytoarchitectonically distinct sub-areas of PreG-HAND based on Brodmann area (BA) probability maps [25]: BA6_HAND_, BA4a_HAND_, and BA4p_HAND_ [26,27] (Fig. 5D). BA4a_HAND_ was additionally manually limited to the gyral crown and lips on both sides of the precentral gyrus.

#### Geometric Modeling and Simulation of Synthetic Axon Morphologies

Axon bends were modelled at various positions in GM and WM. In general, myelinated axons run parallel to the cortical surface in five main regions: Exner’s plexus in layer I [28,9], the outer and inner bands of Baillarger in layers IV and V [29], the superficial [30–32], and the deep WM tracks. These locations are therefore potential sites where axons may undergo distinct bending patterns. Sharp turns into Exner’s plexus have been reported [33], with rising fibers entering the layer perpendicularly and then abruptly reorienting to run parallel to the surface. Fibers in the inner and outer bands of Baillarger are typically collaterals of pyramidal cells [34]. They were included here because cortico-cortical terminating projections may bend into these regions as they pass through. In the WM, both sharp bends, which typically occur when fibers turn into the SWM [35–37], and smoother, gradual bends, which lead into deeper white-matter regions [9,38,39], have been observed. Based on these anatomical patterns, we chose to model three types of sharp axonal bends and the smooth axonal bends:

- For modeling sharp bends of ascending fibers in cortical layers I and IV, we created surfaces for layer I at 6% normalized depth and layer IV at 55% normalized depth, with the depth estimates based on layer boundaries from cortical slices of primates [40]. The layer surfaces were generated using the equivolume approach [41,42] as implemented in the cortech package (https://github.com/simnibs/cortech). To model sharp bends of fibers bending from or into the SWM, an additional surface was placed 0.5 mm below the WM boundary inside the SWM. We then generated sharp bends of axons running in different directions for each surface position by first slicing the cortical surface by a plane (*pyvista.DataObjectFilters.slice* [43]) perpendicular to the surface at angles with a step of 20° (Fig. 5F). The resulting paths were cut to a length of 3 mm, offset in relation to the surface and a straight part was added normal to the surface, ending at least 100 µm from the pial surface. The resulting bend was then adjusted to have a radius of 100 µm.
- Myelinated smooth axon bends of projections to deep WM were generated by solving the Laplace equation on a domain including the handknob area and subsequently tracing streamlines from every surface position on the resulting vector field using a forward Euler integration scheme (Fig. 5G) (for details, see Supplementary Material G). The smoothness of the bends was modulated by manipulating the diffusivity ratio of the WM and GM compartments of the domain. We used a diffusivity ratio of 0.2 (GM/WM), which resulted in bends that largely agreed with experimental observations [38,39].

We simulated activation thresholds for monophasic stimulation with axon diameters of 0.5 to 4 µm for sharp bends and 1 to 6 µm for smooth bends. For sharp bends with an axon diameter of 2 µm, we also applied biphasic stimulation to compare experimental findings to the simulation results. For biphasic stimulation, the E-field direction was also reported for the first phase as for monophasic pulses. But as neural response was determined by the second longer phase, biphasic stimulation was compared with monophasic stimulation having initial E-field in the opposite direction. For the smooth bends, we estimated time constants using TMS waveforms with three different durations as described in Supplementary Material E. All axons were myelinated with an internode length ratio (*L*/*D*) of 100 and a node length of 1 µm. The myelination pattern was aligned so that a node of Ranvier was always placed in the middle of the bend. The first and the last millimeter were made passive and unmyelinated to prevent terminal activation.

## Results

### Realistic human single neuron models show the lowest thresholds at bends in myelinated axons

We evaluated the activation threshold of morphologically realistic model neurons in uniform E-fields with 648 spherically sampled E-field directions (Fig. 1). In unmyelinated models, action potentials were initiated at the AIS for the optimal field direction. Neurons with smoothly bending truncated myelinated axons exhibited action potential initiation at nodes of Ranvier along the axon due to diameter variations (Fig. 1E). In both cases, the lowest thresholds were above 950 V/m. These field intensities are an order of magnitude higher than physiologically realistic intensities occurring in TMS experiments, rendering these cells unlikely to be activated by TMS. In contrast, the remaining neuron models exhibited lower minimum activation thresholds ranging from 235 to 593 V/m (Fig. 1D). For the optimal field direction, action potentials were initiated at nodes of Ranvier close to sharp axonal bends or bifurcations where one collateral branch had a markedly smaller diameter than the other branches. These activation sites were located within layer VI and the superficial white matter.

**Figure 1:**
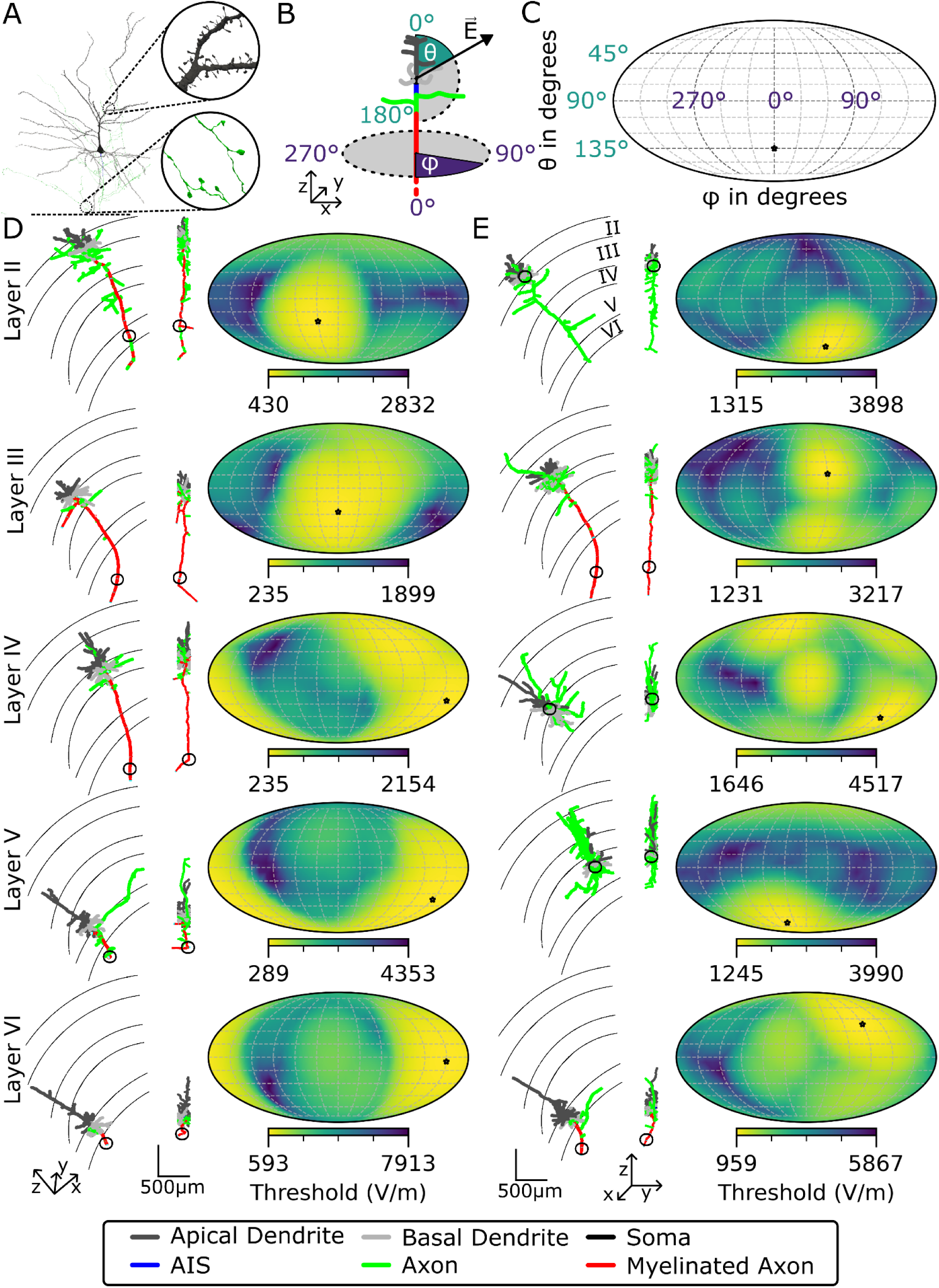
The activation threshold under a spherically sampled uniform E-field driven by a monophasic waveform for pyramidal neurons from layer II, III, IV, V, and VI. (A) The high-resolution 3D mesh of the right layer II neuron with detailed synapses and axonal boutons, which was used to create the compartment model. (B). The spherical coordinate system is illustrated on a schematic neuron with color-coded labels indicating the type of neurite. The somato-dendritic axis is aligned to the z-axis, with polar angle θ (turquoise) and azimuthal angle φ (purple). The E-field vector is indicated with *E*. (C) 3D threshold–direction map for the schematic neuron, projected into 2D using Mollweide projection. The black star, used to indicate the lowest threshold position, points to (180°, 135°). (D and E) Each row shows two neurons from each layer and their threshold–direction map using 648 E-field directions. The neuron morphologies are color-coded in correspondence and plotted on top of cortical layer boundaries. Nodes of Ranvier are not visible at this scale. A black star indicates the minimum threshold in the threshold maps and a black circle indicates the activation site at the minimum threshold.

**Figure 2:**
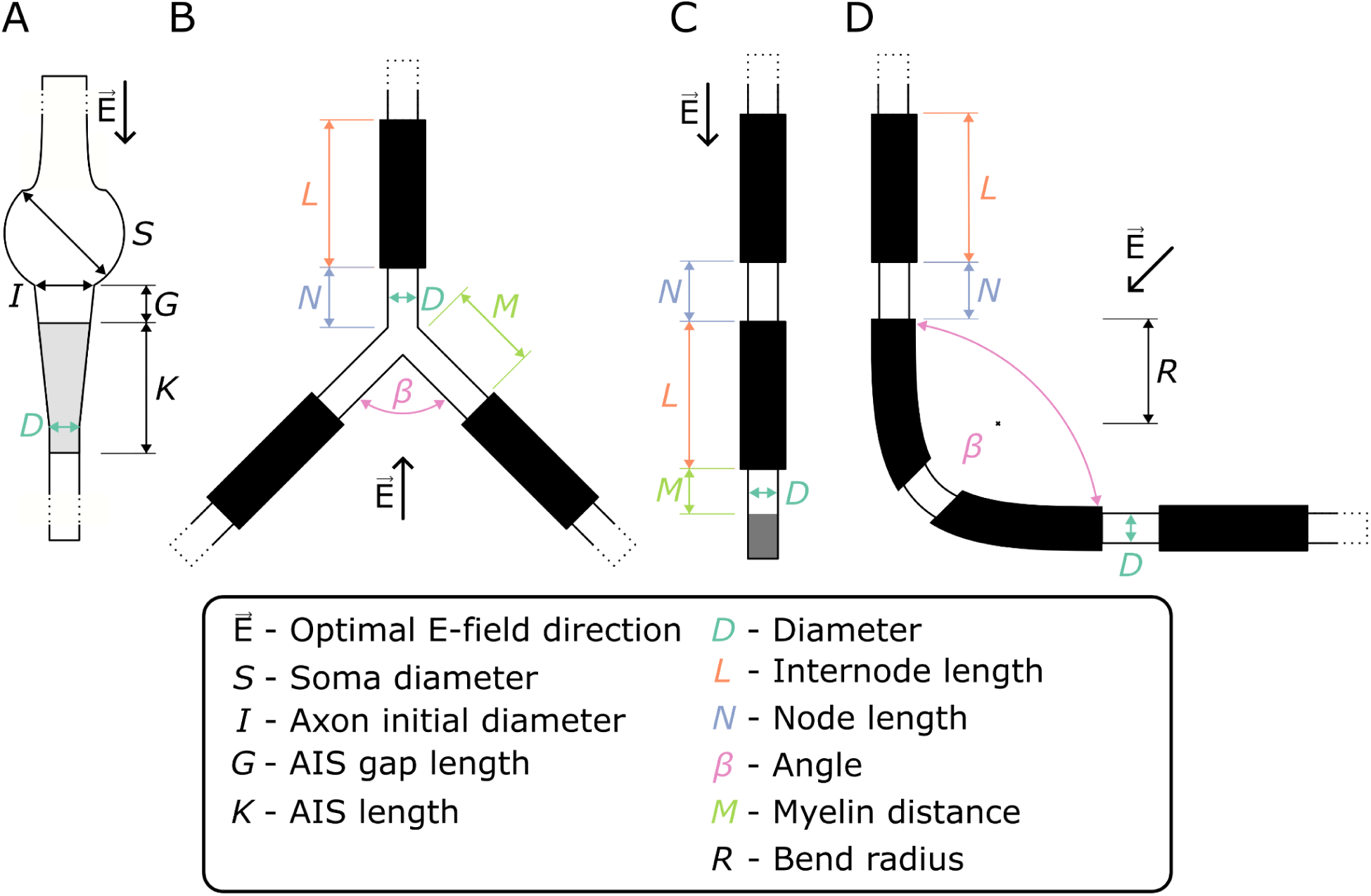
Synthetic neuronal morphologies illustrating geometric and myelination variables. The E-field vector indicates the optimal E-field direction for each model. (A) The ball-and-two-sticks model consists of a spherical soma, a straight apical dendrite emerging from the soma apex, and a straight axon extending from the soma base. The axon diameter tapers over a fixed length. The AIS, shown in gray, begins after an optional gap from the soma. (B) The idealized bifurcation model consists of a parent axon that splits into two child branches of equal diameter. The parent axon and both child branches are myelinated, shown in black, and a node of Ranvier is placed at the branch point. Each child branch may contain an additional unmyelinated gap right after the bifurcation. (C) The cortical axon termination model consists of a straight myelinated axon that terminates in an unmyelinated terminal segment of constant length, shown in gray, following an optional myelin gap. (D) The bend model consists of a myelinated bending axon, with the myelination pattern arranged so that a node of Ranvier lies at the midpoint of the bend.

### Of the known possible mechanisms of axonal activation by TMS, the activation of myelinated axonal bends remains the most likely mechanism

We simulated four models of axonal discontinuities (ball-and-two-sticks, termination, bifurcation, and bend) across a broad parameter space to assess how variations in morphology affect activation thresholds. We present only the parameters with the strongest influence on the threshold for each model. Analyses of the remaining parameters are provided in Supplementary Fig. S9–S11. We consider thresholds < 400 V/m as achievable with standard TMS stimulators and thresholds < 150 V/m as broadly consistent with the estimated electric field strengths at MT [20]. Fig. 3 illustrates approximately exponential decrease in TMS activation threshold with increasing axon diameters for all four models, similar to theoretical predictions [44].

**Figure 3:**
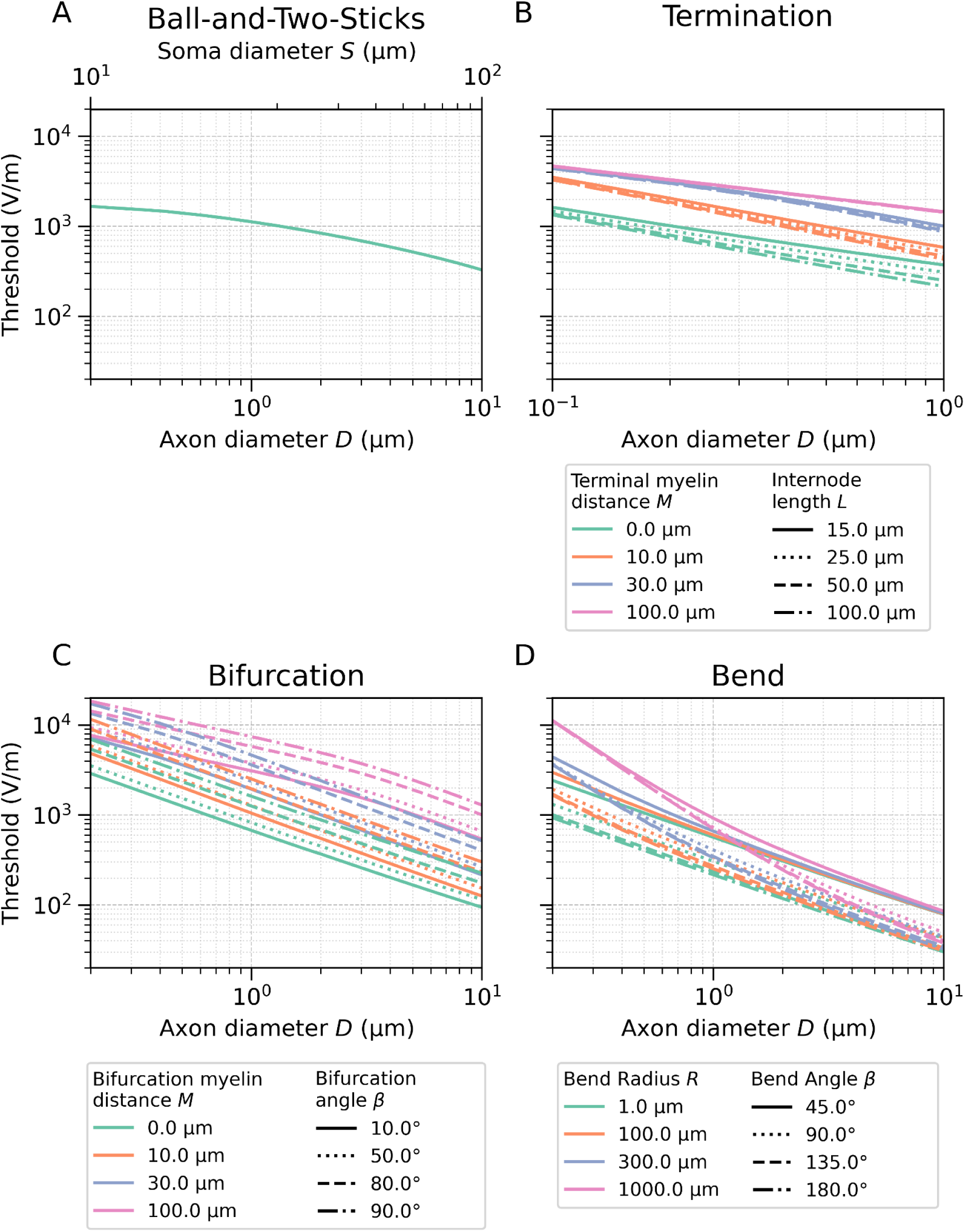
Relationship between key model parameters and the activation thresholds under a uniform E-field driven by a monophasic waveform for (A) the ball-and-two-sticks model, (B) the termination model, (C) the bifurcation model, and (D) the bend model. In (A), the AIS has a length of 30 μm and there is a 9 μm gap between the soma and the AIS. For panels (B–D), the nodes of Ranvier had a length of 1 μm. For panels (C-D), the internode length ratio (*L*/*D*) was 100.

### Ball-and-Two-Sticks

Variations in the length of the soma–AIS gap and in AIS length produced only small changes in thresholds, consistent with the similar biophysical properties assigned to the AIS and axon. Only for soma and axon sizes corresponding to Betz cells, thresholds within the range of TMS were observed, still far exceeding MT estimates.

### Termination

The termination model yielded physiologically plausible thresholds only when the axon was myelinated up to the terminal end, a configuration that has not been reported for terminal branches of mammalian cortical neurons. Achieving MT-level activation would furthermore require terminal diameters greater than 1 µm, which likewise appear inconsistent with existing anatomical reports (for details see Supplementary Material E). The distance between the terminal and the first internode strongly affects the activation threshold (Fig. 4). For all terminal axons with a diameter ranging from 0.1 to 1 µm, the threshold more than doubles when the terminal-internode distance reaches 10 µm.

**Figure 4:**
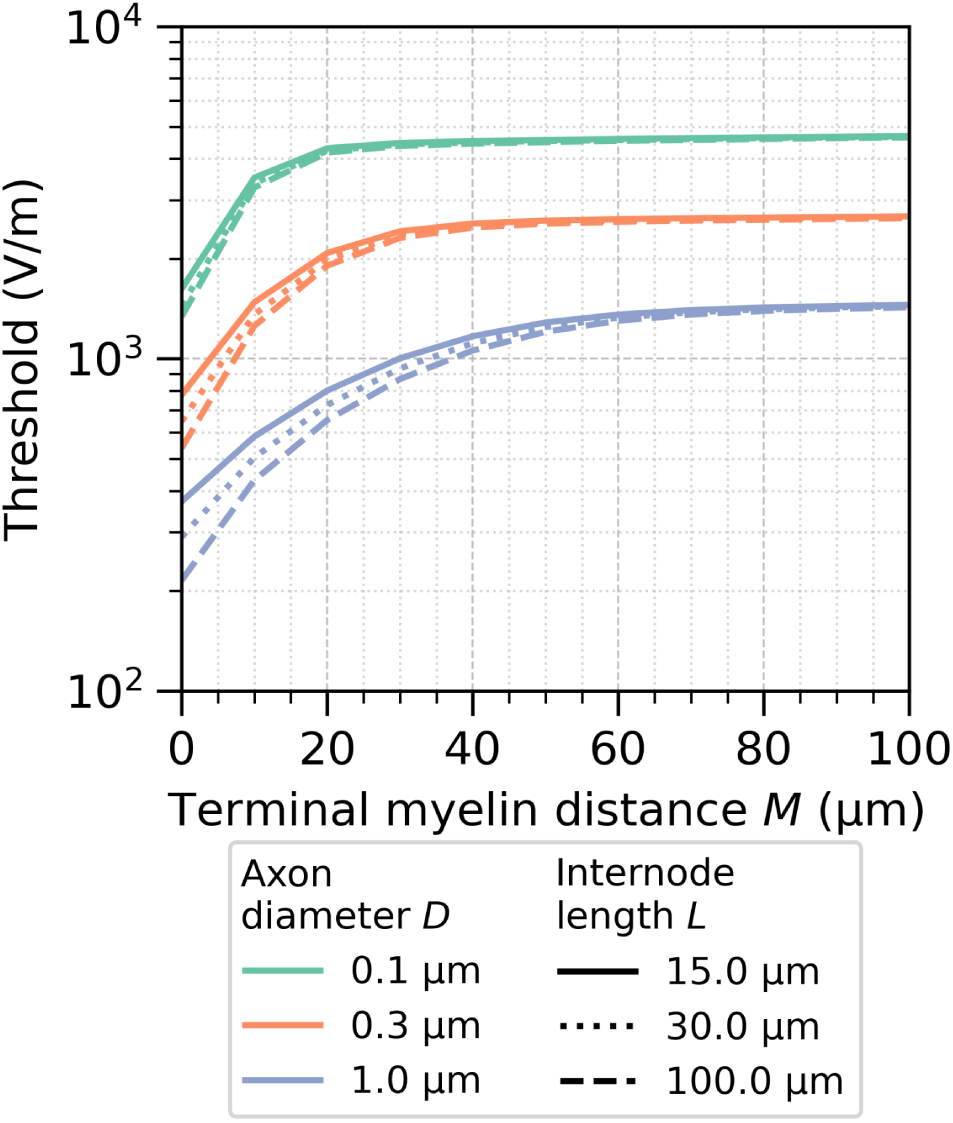
The effect of the distance between the axon terminal and the start of the first internode on the activation threshold under a uniform E-field driven by a monophasic waveform. The nodes of Ranvier had a length of 1 μm.

**Figure 5:**
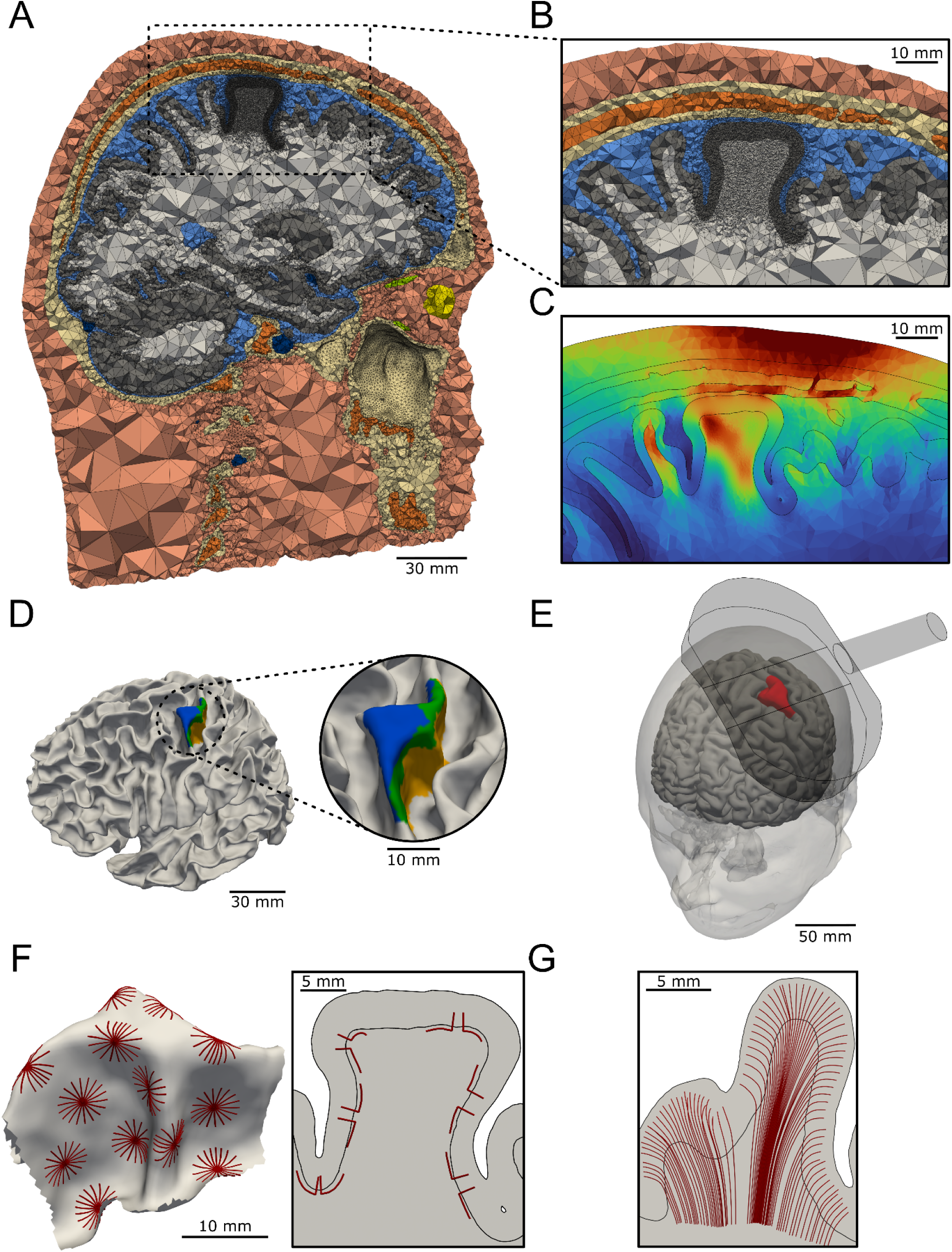
Realistic head model constructed with SimNIBS v4.6 using CHARM with a high-resolution M1 ROI. (A) Clip through the M1 ROI of the head model showing the nine compartments (scalp, compact bone, spongy bone, eye balls, muscle, blood, CSF, GM, and WM). (B) Magnification of the region around the M1 ROI. (C) E-field distribution simulated with SimNIBS from the stimulation setup in (E). (D) Areas of the PreG-HAND are shown on the WM surface. BA4p_HAND_ is yellow, BA4a_HAND_ is green, and the crown and lip part in the M1 of BA6_HAND_ is blue. (E) The setup of the TMS experiment using an MC-B70 TMS coil to stimulate the PreG-HAND in the posterior-anterior (PA) direction. The PreG-HAND ROI is colored red. (F) The 18 directions of sharp bending axons for a selection of surface locations on the left side and a small selection of sharp bending axons projected on a 2D slice of the ROI. (G) A selection of the smooth bending axons projected on a 2D slice of the ROI.

### Bifurcation

Bifurcations reached thresholds below TMS maximum stimulator output (MSO) only when they were fully myelinated and had diameters of > 6 μm. However, these diameters are typically observed only for the main axons of Betz cells, which are unlikely to branch into equally sized collaterals, suggesting that such bifurcations are anatomically uncommon.

### Bend

Axons with diameters greater than 1 µm and realistic bend geometries (e.g., 90° bends with small radii < 100 µm) exhibited thresholds below MSO. Thresholds consistent with MT estimates were obtained for diameters greater than 2 µm. The bend axon models showed an inverse-cosine dependence on E-field misalignment (Supplementary Fig. S3), such that moderate deviations (±30°) increased thresholds by only ∼15%. We also evaluated the model’s sensitivity to an expanded set of morphological and biophysical parameters (Supplementary Fig. S12). For broadly realistic bend configurations with a diameter on the larger end of the histological axon diameter distribution (Table S-I5) with 3 µm, a 90° bend and 100 µm bend radius, threshold variations were generally modest. Node positioning produced only small changes (+2.7% at 20% offset), and altering the node diameter yielded similarly limited effects (0.5 times internode diameter: +0.6%; 2 times: +6.4%). Adjusting the g-ratio had a slightly larger influence (20th percentile vs. median: −7.3%), and sodium-channel density produced a stronger modulation (0.5 times: +10.7%; 2 times: −9%). The number of internodes flanking the bend strongly affected thresholds when the count was below three: Thresholds decreased exponentially with internode count increase, with a ∼14% threshold reduction when increasing from three internodes per side to 14, approaching the asymptotic threshold minimum.

We compared simulated time constants to *in vivo* measurements (τ = 200 ± 33 μs [45,46]), both estimated using the same approach based on evaluating the strength–duration relation. Across axonal discontinuity models, our simulated time constants tended to lie on the higher end of the experimentally measured range. For axonal bends with tighter bend radii, particularly those with larger diameters, the simulated thresholds approached physiological values (Supplementary Fig. S4).

### Thresholds of model myelinated sharp axonal bends within the human PreG-HAND approach experimental motor thresholds

We simulated the response of bending myelinated axons to monophasic TMS stimulation to determine whether their activation thresholds align with *in vivo* measurements. The measured MT for the participant was 59.7 A/µs [20], corresponding to 38% of the 155.3 A/µs MSO [24]. This served as the physiological reference level when evaluating which axons became recruited. We report results for each of the three sub-areas of PreG-HAND, namely BA6_HAND_, BA4a_HAND_ and BA4p_HAND_. E-fields were oriented either posterior–anterior (PA), using the optimal coil orientation from the experiments, or inverted to anterior–posterior (AP).

### Cortical layers I and IV

Sharp bends of ascending fibers in layers I and IV (Figs. S13–S16) exhibited similar activation behavior, with slightly higher thresholds in layer IV, consistent with marginally weaker local E-fields (Figs. S13D vs. S15D). In both cases, BA4a_HAND_ and BA4p_HAND_ were predominantly activated by AP stimulation, in contrast to experimental findings showing higher MT for this direction compared to PA [47]. BA6_HAND_ was activated by both AP and PA stimulation, with a slightly stronger activation from PA stimulation.

### Superficial white matter

SWM bends in BA6_HAND_ showed slightly stronger and more anterior activation under AP versus PA stimulation (Fig. 6). They exhibited the strongest peak activation of the three PreG-HAND sub-areas. For AP stimulation, 14.8% of all axons with a diameter > 2 µm fell below MSO. This corresponded to an area of effect of 105 mm² in which at least one of the 18 axon directions was activated. For PA, the activation side moved more anteriorly and 11.3% of the axons with a diameter > 2 µm fell below MSO, while the area of effect was at 87.9 mm².

**Figure 6:**
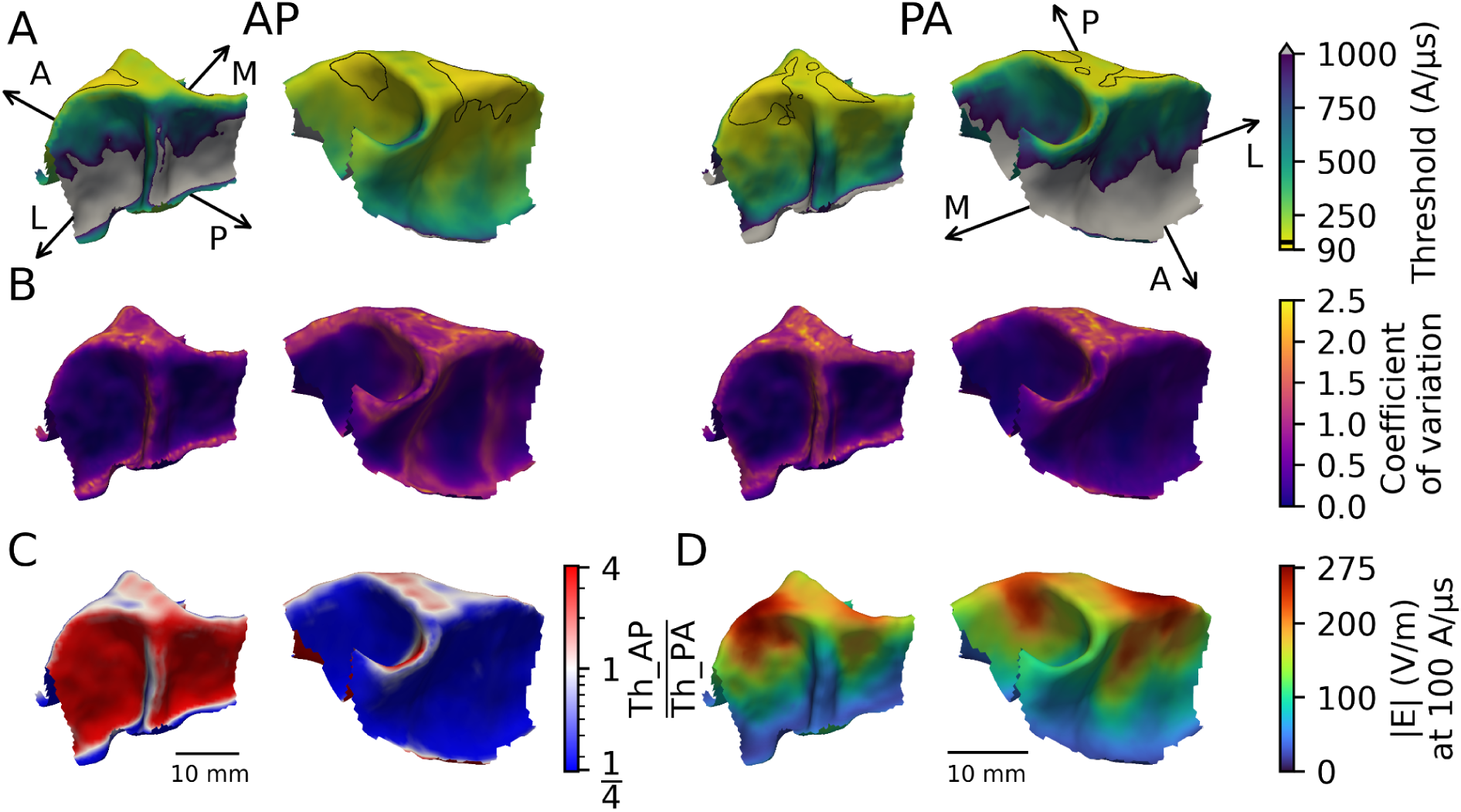
Excitability towards a monophasic pulse of bend axons 0.5 mm below the WM surface (bend radius 100 µm, diameter 2 µm, and internode length ratio of *L*/*D* 100). For each surface position, 18 axon directions are simulated. (A) The minimum activation thresholds in A/µs for each position for E-fields oriented in the anterior-posterior (AP, left) and posterior-anterior (PA, right) directions. The black border surrounds the regions with a minimum threshold at or below the 5th percentile (AP: 122 A/µs, PA: 129 A/µs). Medial-lateral and anterior-posterior anatomical axes are annotated on the surfaces. (B) The coefficient of variation 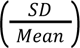 of activation thresholds across the 18 axon orientations at each surface position. (C) Log-scaled ratio between AP and PA stimulation thresholds at each surface position. (D) E-field magnitude interpolated at 0.5 mm below the WM surface for a stimulator intensity of 100 A/µs.

SWM bends in BA4a_HAND_ and BA4p_HAND_ were predominantly activated by PA stimulation (Fig. 7), in line with experimental results [47]. In BA4a_HAND_, most fibers were recruited below MSO compared to the other areas. Under PA stimulation, 18.4% of axons > 2 µm were recruited below MSO, corresponding to an area of effect of 113 mm². The smallest activation appeared in area BA4p_HAND_, where the E-field was the weakest due to the increased distance to the coil. Only 0.23% of the fibers with a diameter > 2 µm were activated under PA stimulation below MSO, yielding an area of effect of 6.9 mm².

**Figure 7:**
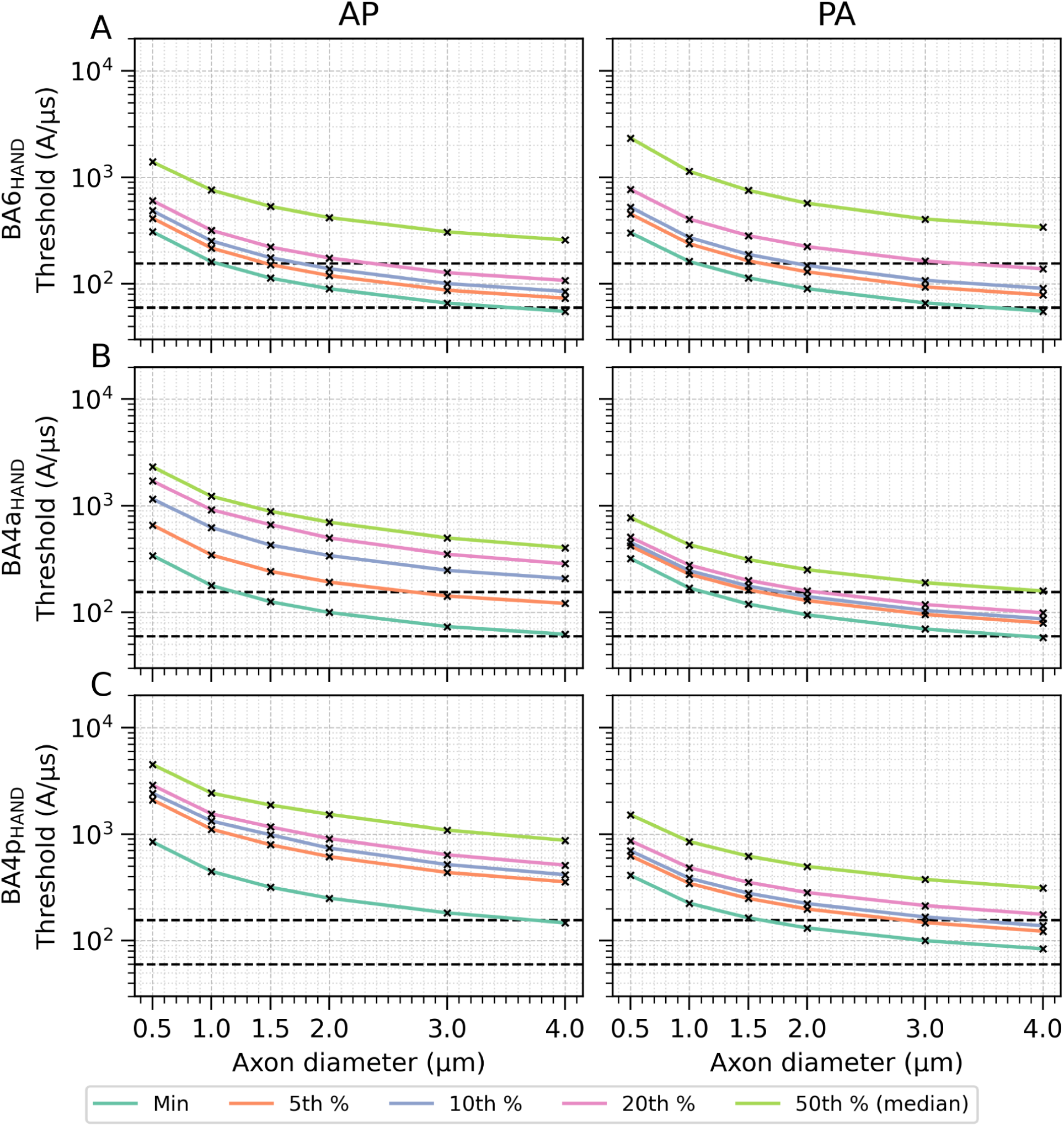
Excitability towards a monophasic pulse for populations of axon bends located 0.5 mm below the WM surface (bend radius 100 µm and internode length ratio of *L/D* 100). Simulations were run separately for six axon diameters (0.5, 1.0, 1.5, 2.0, 3.0, and 4.0 µm). For each condition, the minimum activation threshold and four percentiles (5th, 10th, 20th, and 50th) of the threshold distribution for 18 rotations per location are shown for three cortical regions: (A) BA6_HAND_ with 555 locations, (B) BA4a_HAND_ with 558 locations and (C) BA4p_HAND_ with 1537 locations. The lower dashed line marks the experimentally estimated MT of 59.7 A/µs, while the upper dashed line marks the maximum stimulator output at 155.3 A/µs.

In general, only a few axons were activated below MT. The highest amount occurred in BA6_HAND_, where 0.16% (AP) and 0.25% (PA) of the axons > 4 µm were stimulated below MT. This corresponded to an area of effect of 3.62 mm² and 2.92 mm², respectively. The variation in activation threshold between the 18 different axon directions after the bend is shown in Fig. 6B (see Figs. S13B & S15B for further visualizations). Directional dependence was greatest in BA6_HAND_, where the local field orientation relative to axon direction varied strongly across the crown. In contrast, BA4a_HAND_ and BA4p_HAND_ showed lower directional sensitivity because the field remained largely parallel to the pre-bend axon orientation.

### Monophasic versus biphasic pulses

For bending axons with a diameter of 2 µm, we additionally applied biphasic TMS stimulation (Figs. S17–S19). As in the monophasic case, bends located in layers I and IV exhibited spatial activation patterns that were inverted relative to experimental observations [47], and bends in the SWM behaved consistently with experiments [47]. To put our results for different stimulation paradigms into context, we compared them to *in vivo* measurements of MTs in the PreG-HAND [47].

At posterior locations, the threshold ratio between AP and PA biphasic pulses was ∼1.35, while the ratio between posterior AP and anterior PA biphasic stimulation was approximately ∼1. In experiments, the corresponding AP versus PA biphasic ratio was 1.17.

In contrast, monophasic stimulation produced substantially larger polarity-dependent differences in the model. For anterior locations, the ratio between PA and AP stimulation was ∼4, whereas the ratio between posterior PA and anterior AP stimulation was ∼1. Experimental measurements reported a mean ratio of 1.34 for PA versus AP stimulation.

Direct comparisons between biphasic and monophasic stimulation further highlighted the lower activation thresholds for biphasic stimulation. The ratio between AP biphasic and PA monophasic stimulation at anterior locations was approximately 1.29, as was the ratio between PA biphasic and AP monophasic stimulation at posterior locations. Experimentally, the corresponding mean ratios were 1.4 for AP biphasic versus PA monophasic stimulation and 1.61 for PA biphasic versus AP monophasic stimulation. Thus, the interpretation of this comparison depends strongly on the hypothesis where the stimulation occurs within the motor cortex.

### Models of myelinated smooth axonal bends within the human PreG-HAND are recruited only at high intensities

Thresholds under monophasic stimulation for the smoothly bending fibers are shown in Figs. 8 and 9. We find high thresholds in deep areas (including BA4p_HAND_) for both field directions. At the gyral crown (BA6_HAND_), thresholds were generally lower, and PA field directions led to lower thresholds compared to AP. The lowest thresholds occurred for PA at the superficial parts of the sulcal wall corresponding to BA4a_HAND_, whereby changing to AP direction resulted in high thresholds at those positions. Thus, the local thresholds depended on a combination of local field strength, bending curvature and field direction.

**Figure 8:**
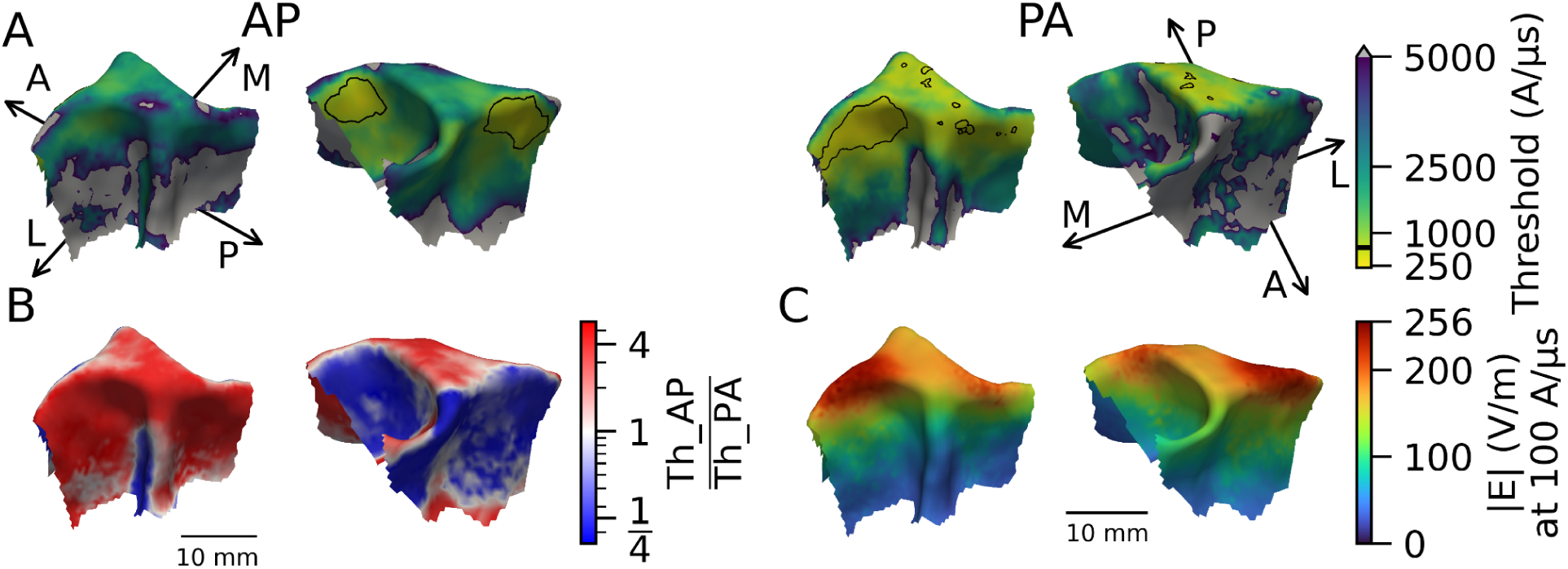
Excitability towards a monophasic TMS-pulse of smoothly bending axons projecting into deep WM. The axons start 0.5 mm below the pial surface and terminate in deep WM. They have a diameter of 2 µm and an internode length ratio (*L*/*D*) of 100. (A) The minimum activation thresholds (in A/µs) for each position for E-fields oriented in the anterior-posterior (AP, left) and posterior-anterior (PA, right) directions. The black border surrounds the regions with a minimum threshold at or below the 5th percentile (AP: 774 A/µs, PA: 558 A/µs). Medial-lateral and anterior-posterior anatomical axes are annotated on the surfaces. (B) Log-scaled ratio between AP and PA stimulation thresholds at each surface position. (C) E-field magnitude interpolated at the point of minimum bend radius of each simulated fiber.

**Figure 9:**
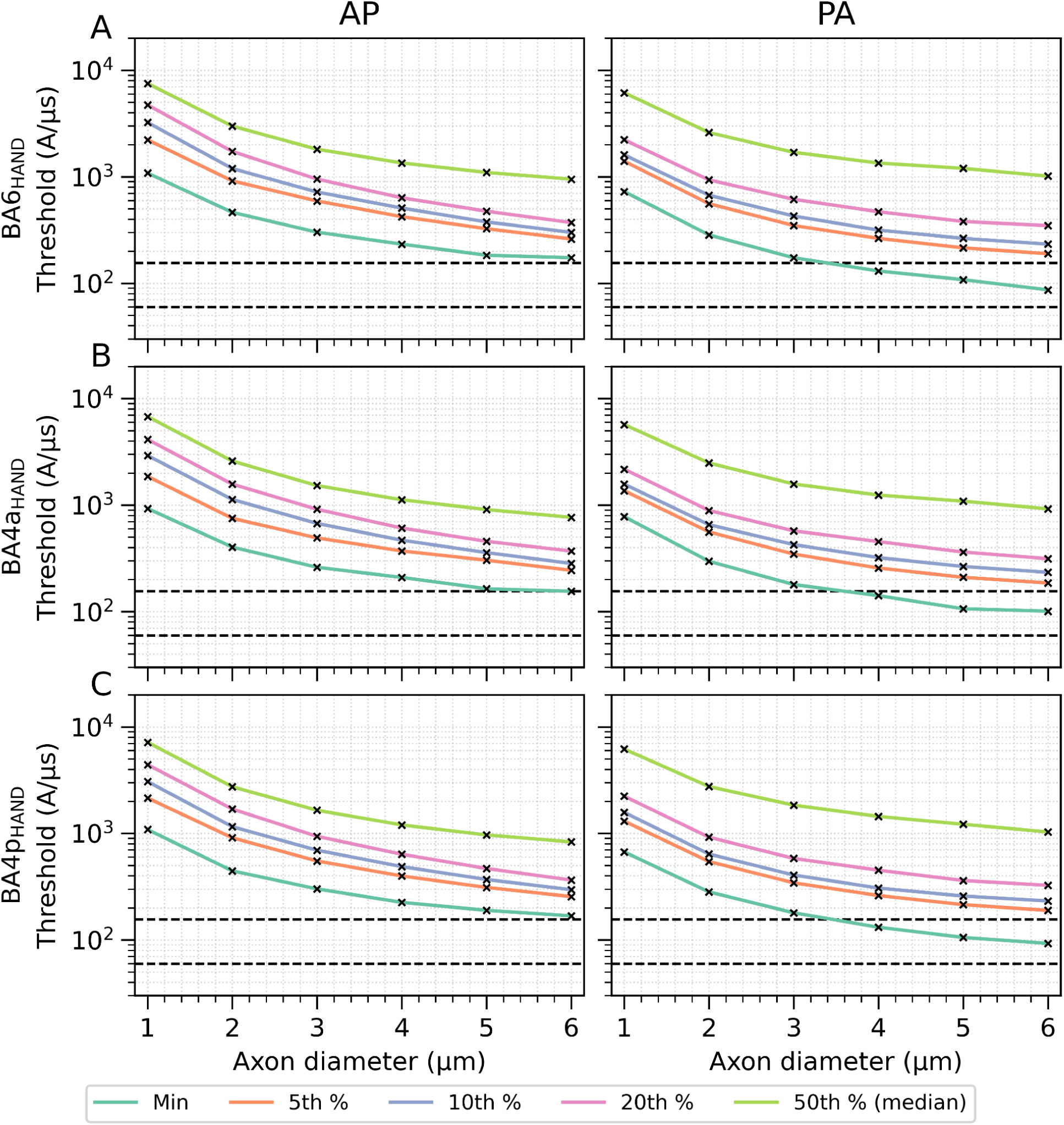
Excitability towards a monophasic pulse of smoothly bending axon projections into deep WM. The axons start 0.5 mm below the pial surface and terminate in deep WM and have an internode length factor (*L*/*D*) of 100. Simulations were run separately for six axon diameters (1.0, 2.0, 3.0, 4.0, 5.0, and 6.0 µm). For each condition, the minimum activation threshold and four percentiles (5th, 10th, 20th, and 50th) are shown for three cortical regions: (A) BA6_HAND_ with 550 locations, (B) BA4a_HAND_ with 553 locations and (C) BA4p_HAND_ with 1517 locations. The lower dashed line marks the experimentally estimated MT of 59.7 A/µs, while the upper dashed line marks the maximum stimulator output at 155.3 A/µs.

Fig. 9 shows a summary of the thresholds for different axon diameters, revealing that none of the smoothly bending axons produced thresholds low enough to explain the experimentally determined MT. Only a few of the larger axons with diameters > 3.5 µm were activated by PA stimulation below MSO. Similarly, while biphasic pulses showed the expected reduction in activation thresholds (Supplementary Fig. S20), the reduction was not significant enough to explain physiologically measured MTs.

## Discussion

The study examined the excitability of axons to TMS and tested several distinct positions along their trajectories, including the initial parts at the soma, terminations, and bifurcations in GM, and distal bends extending into superficial and deeper WM. Neuron models were reconstructed from high-resolution human electron microscopy data using a novel modeling pipeline, and were complemented by histologically informed simplified representations of candidate target structures. Overall, we showed that the AIS, symmetric bifurcations, and axon terminals are unlikely to be systematically stimulated during TMS. In contrast, sharp myelinated axonal bends within superficial white matter emerged as preferential activation sites. Simulations of these bends embedded in realistic human head models yielded activation thresholds approaching experimental MTs.

### Comparison with experimental results

The simulations of morphologically and biophysically realistic pyramidal neuron models, exposed to uniform E-fields, revealed markedly higher activation thresholds at axonal terminals than those reported in previous studies. Instead, the lowest thresholds occurred at bends of fully myelinated main axons of cortical pyramidal neurons near the GM-WM boundary. This discrepancy is largely attributable to differences in myelination patterns: Our reconstructions, directly derived from serial EM [9], exhibited realistic heterogeneous myelination, whereas earlier models assumed artificially uniform, fully myelinated axons [6,48]. A limitation of our dataset is that all reconstructed neurons originated from the left human anterior middle temporal gyrus, a region known to have lower myelin content than areas such as the motor cortex [49]. This regional difference could arise from a lower proportion of myelinated neurons or from individually more extensive myelination in motor cortex neurons. To our knowledge, however, no reports have described fully myelinated axon terminal branches in the mammalian brain. On the contrary, histological studies report thin [35,36] and highly branched [35–37] terminal arbors, both of which have been shown to reduce the likelihood of myelination [50,51]. Even if terminal branches were substantially myelinated, effective activation would require myelination extending to within < 20 μm of the terminal, as thresholds increased exponentially the further the myelin ends from the terminal (Fig. 4). Moreover, terminals would need diameters exceeding those of more than half of the axon population to yield physiologically plausible thresholds (see Tables S-I4–S-I6), which seems unlikely, given that terminal branches are typically among the thinnest axons [35,36]. Taken together, these observations suggest that axon terminals are unlikely to represent the primary targets of TMS. While occasional terminal activation cannot be excluded, a systematic targeting of axon terminals is improbable based on current histological evidence.

As expected from cable theory, bends are, in general, less excitable to TMS-induced activation than terminals. Despite this fact, our simulations of morphologically and biophysically realistic human neurons showed that activation thresholds were lowest at axonal bends within the SWM. Examples from histological studies confirm the notion of myelinated axons making sharp bends when traversing from the GM into the SWM [35–37]. Also, our further results support myelinated axon bends in SWM as possible activation sites. First, on a general level, myelinated bends consistently exhibited the lowest thresholds within realistic morphological parameter ranges among the four types of axonal discontinuities studied in uniform E-fields. Second, myelinated axon bends with tight bend radii, particularly those with larger diameters, approached experimentally measured time constants. Third, within the PreG-HAND area of a realistic head model, histologically realistic axon bends in SWM approached physiologically measured thresholds. When inverting the field direction, the change in activation patterns mirrored the established predictions: Stimulation was localized to the crown and lip regions for AP pulses and shifted anteriorly for PA pulses [10,20]. Finally, our results also align with experimental evidence suggesting that activation generally occurs at distal axonal sites, including the effects of voltage-gated sodium channel blockade and the insensitivity of the I_1_ wave to voluntary contraction, GABA agonists and inhibitory paired-pulse paradigms [2].

In contrast, axons bending into Exner’s plexus (layer I) or the bands of Baillarger (layers IV and V) exhibited activation patterns opposite to those expected for motor output, making their direct contribution to TMS-induced corticospinal descending volleys unlikely. Although our model predicts that sufficiently large axons in Exner’s plexus could exhibit relatively low thresholds, their actual morphology, diameter distribution, and functional role remain poorly understood. These fibers may, however, contribute to sub-MT inhibitory effects reported in physiological studies [52], a hypothesis that warrants targeted anatomical and functional investigation.

Smoothly bending myelinated axons projecting into the deep WM had higher thresholds than sharp bends in SWM. On average, these axons are also expected to be larger, which partially offsets their higher bend radius in terms of threshold. Very thick axons bending smoothly into deeper WM are expected from Betz cells in BA4a and most predominantly in BA4p. Thus, D-waves, which require higher simulation intensities [53], could originate from smoothly bending thick axons of motor neurons in the sulcal wall of the PreG-HAND. While we did not use a coil position optimized for D-waves in our simulations, the results still suggested the activation of very thick, smoothly bending axons at high stimulator outputs slightly below MSO. An interesting observation is that the simulated time constants of smooth bends were more variable and can far exceed those of sharp SWM bends. We suggest that the activation behavior of sharp bends is dominated by the node of Ranvier at the bend, so that fitting their time constants using a single pair of resistance and capacitance values will give reliable results within the physiologically measured range. In contrast, the behavior of smooth bends can depend on the spatial interactions between neighboring membrane parts with different properties along the bend, which cannot be accurately fitted with a single resistance-capacitance pair. However, this limitation applies to both the simulated and measured time constants to the same extent. Thus, our findings suggest that the measured time constants are in line with sharp, but may not be consistent in all regions with smooth, myelinated bends.

### Previous studies

Our study addresses limitations of previous E-field and multiscale simulation studies that restricted their analyses only to specific parts of PreG-HAND [20] or tested isolated components of the complex axonal morphology [3,6,10]. While informative, these studies provided only a partial view of axonal excitability, which can bias interpretations by neglecting other relevant sites. For example, the analyses in Bungert et al. [20] were based on the assumption that the neural targets of TMS were in GM. Aberra et al. [6] focused on axons in proximity to the soma in the GM and relied on likely unrealistic assumptions about terminal myelination to achieve low thresholds. Laakso et al. [10] suggested deeper targets using probabilistic activation target modelling, but relied on multiscale simulation for validation that used unrealistic thickness estimates for the myelinated axons (10 μm) and estimated bending geometries from diffusion MRI, which is unreliable close to the GM. The simulated MTs remained a factor 2 above experimental values. While still to be confirmed, we suggest that the experimental results of Bungert et al. [20] and Laakso et al. [10] are likely in line with activation in SWM.

### Limitations

Despite new insights enabled by large-scale high-resolution histology datasets, limitations remain in both E-field and biophysical modeling:

#### E-field simulations

The simulations rely on tissue conductivity values that are challenging to measure accurately. The GM is approximately isotropic at the macroscale, although mesoscale conductivity variations across cortical layers have been reported [54]. These mesoscale variations may be related to cortical features such as blood vessels, Exner’s plexus, the bands of Baillarger, and local variations in cell and soma density. The putative impact of mesoscale conductivity variations in GM on the activation thresholds was not evaluated in our study. WM exhibits strongly anisotropic conductivity that is governed by fiber orientation. The microstructural transition between GM and WM remains poorly characterized in terms of its spatial extent, geometry, and conductivity gradients, even though these gradients can substantially affect the computed E-field. In this study, we used a conservative approach to modeling the conductivity transition between the GM and the WM (Supplementary Material F). Using our conductivity model, the sharp bends showed slightly higher thresholds compared to standard isotropic conductivities, while the activation thresholds for smooth bending axons were reduced but remained high compared to the sharp bending axons.

#### Cell biophysics

The biophysics of existing neuronal models rely heavily on parameters derived from juvenile rodent tissue and provide limited information on distal axonal compartments, including nodes of Ranvier, axon terminals, and unmyelinated segments. Studies investigating axonal ion channel dynamics suggest complex dynamics even along the unmyelinated axon [55]. Similarly, the electrophysiological properties of myelinated internodes remain incompletely understood. Most parameter estimates are derived from small samples, potentially affecting the reliability of current biophysical simulations.

#### Diameters of myelinated axons

An important limitation of our analysis is the uncertainty in histologically derived estimates of axon diameters, as they strongly influenced the simulated activation thresholds. Yet, histological measurements are prone to preparation-induced artifacts. Recent work comparing chemically fixed and cryo-preserved tissue suggests diameter-dependent shrinkage of up to ∼53%, with potentially even larger effects for the largest fibers [17]. Because no true *in vivo* ground-truth morphology exists, such distortions introduce substantial uncertainty when interpreting axon diameter measurements. Diameter estimation is further complicated by pronounced local fluctuations along individual fibers, so that single-section EM measurements can substantially over- or underestimate the true fiber-averaged diameter. This bias tends to be most pronounced at the tails of the diameter distribution, where capturing a single atypical cross-section can disproportionately distort the inferred extremes. Moreover, axon diameters in cortical tissue follow a highly skewed distribution. As a consequence, small or spatially limited samples might fail to include the relatively rare large-diameter axons that might be relevant for TMS because of their lower activation thresholds. Taken together, these factors indicate that histological measurements of axon diameter should be interpreted cautiously. In particular, incorporating shrinkage-corrected diameter distributions could bring simulated activation behavior into closer alignment with *in vivo* measurements.

#### Neuron and axon morphologies

Existing neuronal reconstructions usually rely on biocytin-filled cortical neurons imaged with light microscopy in cleared brain slices. However, standard light imaging approaches often lack the resolution needed to capture fine axonal collaterals with diameters below ∼0.25 µm [56]. Diameter measurements are also limited by the low spatial resolution of the imaging process, preventing accurate assessment of local diameter fluctuations. In addition, the positions of slice boundaries are rarely documented, which can lead to the erroneous classification of transected axons as terminal branches. Information about myelination is also largely absent from these datasets because it requires additional immunofluorescence labeling and imaging that are not part of standard reconstruction workflows. Together, these limitations introduce morphological inaccuracies that restrict the direct reuse of existing neuronal reconstructions for TMS simulations and can lead to systematic errors in both predicted activation thresholds and inferred sites of action potential initiation.

We resolved the above limitations by reconstructing realistic human neuronal morphologies from a high-resolution EM dataset. However, our semiautomatic pipeline is computationally and time demanding, resulting in a relatively small sample size that may not fully capture the diversity of human neurons. Although we selected representative morphologies, larger-scale reconstruction initiatives will be necessary to obtain a more comprehensive characterization of cellular morphologies. Consequently, the myelin distribution across entire cells remains poorly understood in humans, especially within motor cortical areas where access to surgically resected, living tissue is very limited.

Despite recent histological work on the SWM [31], quantitative descriptions of axonal geometry remain sparse across large portions of the brain. The axonal bending parameters and bending locations used in this study were inferred from a limited set of experimentally extracted data points and therefore do not capture the full variability of axonal morphology. Our results demonstrated a clear directional dependence of excitation in the gyral crown for axons bending in different orientations within the SWM. Axons bending toward the posterior crown of the precentral gyrus were preferentially activated under PA stimulation, whereas anterior-directed fibers projecting into BA6 were more responsive under AP stimulation. This sensitivity to local microstructural organization indicates that accurate evaluation of stimulation effects requires more comprehensive and region-specific measurements of SWM microarchitecture.

### Conclusion

Our analysis indicates that sharp myelinated axonal bends in SWM are a probable site of TMS induced activation, based on their particularly low activation thresholds and their high abundance compared to other candidate structures. Activation at axon terminals is improbable because histological evidence shows little or no myelin in the vicinity of terminals, which is required for physiologically realistic activation thresholds. Smooth myelinated bends are markedly less excitable than SWM bends. Their thresholds are below maximum stimulator output for only a small fraction of axons with very large diameters. Axon initial segments and bifurcations have even higher stimulation thresholds that reach realistic ranges only for diameters corresponding to the largest Betz cells. Overall, among all modeled candidate structures, SWM bends exhibited activation patterns most consistent with experimental results. Our study makes clear-cut hypotheses about the morphological features of axons that are critical for low TMS excitation thresholds, and demonstrates the relevance of detailed histological and biophysical information to support robust modeling results.

## Acknowledgments

This work was supported by the National Institute of Mental Health and the National Institute of Neurological Disorders and Stroke of the US National Institutes of Health under grant Nos. R01MH128422, R01 NS117405, and U01NS126052. A. Thielscher was supported by the Lundbeck Foundation grant R313-2019-622. The content is solely the responsibility of the authors and does not necessarily represent the official views of the funding agency.

## Declaration of Generative AI and AI-Assisted Technologies

During the preparation of this work, the authors used Generative AI in order to improve readability. After using these tools, the authors reviewed and edited the content as needed and take full responsibility for the content of the publication.

## CRediT Authorship Contribution Statement

H. Worbs: Conceptualization, Methodology, Software, Validation, Formal analysis, Investigation, Data Curation, Writing - Original Draft, Writing - Review & Editing, Visualization. D. Nielsen: Methodology, Software, Investigation, Writing - Original Draft, Visualization. B. Wang: Conceptualization, Formal analysis, Investigation, Writing - Review & Editing. W. Hansen: Software, Validation, Data Curation. W. M. Grill: Conceptualization, Writing - Review & Editing, Project administration, Funding acquisition. A. V. Peterchev: Conceptualization, Writing - Review & Editing, Project administration, Funding acquisition. Thielscher: Conceptualization, Methodology, Investigation, Writing - Original Draft, Writing - Review & Editing, Project administration, Funding acquisition

## Conflict of Interest

H. Worbs, J. D. Nielsen, B. Wang, J. W. Hansen, W. M. Grill and A. Thielscher declare no relevant conflict of interest. A. V. Peterchev has received patent royalties and consulting fees from Rogue Research; equity options, scientific advisory board membership, and consulting fees from Ampa Health; equity options, consulting fees, and travel support from Magnetic Tides; consulting fees from Soterix Medical; equipment loans from MagVenture; hardware donations from Magstim; and research funding from Motif.

