## Supplemental Material for "Histologically Informed Multiscale Modeling of the Neuronal Elements Activated by TMS"

### Contents

Supplementary Material A: Biophysics and myelin models
Supplementary Material B: Simulation of conduction velocity
Supplementary Material C: Reconstruction of neuron models for compartment simulations
Supplementary Material D: Morphology of biological neuronal components
Supplementary Material E: Simulation of model time constants
Supplementary Material F: Creation of anisotropic conductivity distribution
Supplementary Material G: Smooth bending white matter axon projections
Supplementary Material H: Supplementary Figures S1–S20
Supplementary Material I: Supplementary Tables S-I1–S-I9

### Supplementary Material A

#### Biophysics Model

The ion channel parameters in the biophysical models were originally estimated from brain slices of the somatosensory cortex of P14 male Wistar (Han) rats [1]. The initial release included compartment-specific models for the soma, apical and basal dendrites, and the AIS. Aberra et al. [2] extended this set by introducing additional models for myelinated axonal segments, nodes of Ranvier, unmyelinated axon regions, and axon terminals. These axonal compartment models were derived from the AIS model by removing calcium channels. For the nodes of Ranvier and axon terminals, the transient sodium conductance was doubled to reflect their higher concentration of sodium channels. The fast sodium channel ($m^3h$) used in these models was further adapted by scaling the time constant by 0.4 for both the activation ($m$) and inactivation ($h$) gates [3].

#### Myelin Model

We modeled a myelinated axon with inner and outer radii of $R_i$ and $R_o$, respectively, and a g-ratio of $g = R_i/R_o$. Given the thickness of a single myelin membrane $d$, which was approximately 5 nm [4,5], the number of myelin membranes, $n$, was approximately given as

$$n = 2\lfloor\frac{R_o - R_i}{2d}\rfloor \approx \frac{1-g}{g\cdot x}. \quad \text{(A1)}$$

Here, with two membranes per lamella, $n$ was twice the number of myelin lamella and hence an even number, and $x = d/R_i$ was the ratio between the membrane thickness and the inner radius and had an order-of-magnitude $10^{-3}$ to $10^{-2}$ for axons with radius between 0.5 and 5 μm. The radius of the $k$-th myelin membrane was

$$R^{(k)} = R_i + (k - 0.5)\cdot d,\ k = 1, 2, ..., n. \quad \text{(A2)}$$

The specific capacitance and leakage conductance of a single myelin membrane were given as $\hat{c}_m$ and $\hat{g}_m$, respectively. The $n$ myelin membranes had a total impedance per unit length of

$$Z_m(\omega) = \sum_{k=1}^{n} \frac{1}{\left(j\omega\hat{c}_m + \hat{g}_m\right)\cdot 2\pi R^{(k)}} = \frac{n}{\left(j\omega\hat{c}_m + \hat{g}_m\right)\cdot 2\pi\cdot HM_N\left(R^{(k)}\right)} \quad \text{(A3)}$$

where $HM_n$ was the harmonic mean of $n$ terms, and ω was the frequency. Using effective specific capacitance and conductance $c_m$ and $g_m$ for an equivalent single membrane of radius $R_i$, the total impedance can also be described as

$$Z_m(\omega) = \frac{1}{\left(j\omega c_m + g_m\right)\cdot 2\pi R_i}, \quad \text{(A4)}$$

which yielded

$$\frac{c_m}{\hat{c}_m} = \frac{g_m}{\hat{g}_m} = \frac{HM_N\left(R^{(k)}\right)}{N\cdot R_i} \approx \frac{g\cdot x}{1-g}\cdot HM_n(1 + k\cdot x). \quad \text{(A5)}$$

Hence, the effective specific capacitance and conductance of the entire myelin sheath was approximately reduced to $x$ times that of a single membrane, given that the remaining terms $g\cdot(1 - g)^{-1} \cdot HM_n$ had an order-of-magnitude of one.

The total effective membrane properties of the axon were a combination of myelin and internodal axon membrane capacitance and leakage conductance. To consider the equivalent single-cable axon of radius $R_i$, the total impedance was a series combination of the impedance of the axon and myelin, with the periaxonal space ignored for simplicity. With $c_i$ and $g_i$ respectively being the specific membrane capacitance and conductance of the internodal axon, the total impedance per unit length was given by

$$Z = Z_i + Z_m = Z_m \cdot \left(1 + \frac{Z_i}{Z_m}\right) = \frac{1+\eta(\omega)}{\left(j\omega c_m + g_m\right)\cdot 2\pi R_i} \tag{A6}$$

with $\eta(\omega) = \frac{Z_i}{Z_m} = \frac{j\omega c_m + g_m}{j\omega c_i + g_i}$ as the impedance ratio between the internodal axon and the total myelin. Because $c_m \approx x\cdot\hat{c}_m \ll c_i$ and $g_m \approx x\cdot\hat{g}_m \ll g_i$, and hence $|\eta(\omega)| \ll 1$ for all $\omega$. Therefore, the myelin dominated the membrane properties, with the internode's membrane properties resulting in a small frequency-dependent correction term. For simplicity, we chose to obtain frequency-independent parameters matching the parameters at dc and the high-frequency asymptote with

$$g_{asym} = \frac{g_m g_i}{g_m + g_i},\ c_{asym} = \frac{c_m c_i}{c_m + c_i}. \tag{A7}$$

For typical myelin and axon parameters, this resulted in maximum relative magnitude errors of less than 2% and maximum phase errors of less than 2° in the frequency range between 100 Hz and 10 kHz, which covered neural activity (action potentials) and stimulation pulse frequencies (corresponding to 100 μs to 10 ms pulse durations).

Specific membrane capacitance $\hat{c}_m$, $c_i$, and $c_n$ were consistently around 1 μF/cm$^2$ for myelin and axon internodes and nodes and other biological membranes (range 0.5 to 1.5 μF/cm$^2$) [4,6–10]. Specific leakage conductance $\hat{g}_m$ per myelin membrane was between 0.1 and 2 mS/cm$^2$ [6,7,9,10], and $g_i$ for the internode was between 0.04 and 0.1 mS/cm$^2$ [10,11].

To estimate the g-ratio for myelinated axon models, we used histological measurements published by Ruthig et al. [12] showing the relation between the inner and outer diameter of SWM fibers. The measurements were made below and in between M1 and S1, as well as within V2. We fitted logarithmic functions to the binned median of the distribution between the 50th (median) and the 1st percentile to capture a range of g-ratios. Fig. S1 shows the distribution as well as the logarithmic fit.

### Supplementary Material B

#### Simulation of Conduction Velocity

To evaluate the biophysical realism of our axon model, we quantified conduction velocity across a physiologically relevant parameter range, varying axon diameters ($D$ = 0.2–10 μm), internode length ratios ($L/D$ = 60, 100, 120), node of Ranvier lengths (1 or 3 μm), and g-ratio fits (1st and 50th percentile). We used a higher temporal resolution (time steps of 1 μs) to accurately capture the arrival time of the action potential. Each simulation used a straight axon flanked by 1 mm of passive axon at both ends, followed by a 200 μm unmyelinated transition region to minimize boundary artifacts. The central myelinated section consisted of 20 node-internode pairs, ensuring sufficient internodes for stable velocity estimation. We initiated action potentials by injecting a brief current pulse at the first node of Ranvier and calculated conduction velocity from the propagation time between the third and third-to-last nodes. The resulting conduction-velocity profiles as a function of axon diameter are shown in Fig. S2.

Firmin et al. [13] measured conduction velocities between the medullary pyramid and the primary motor cortex (M1) in macaque monkeys and reported values ranging from 5 to 94 m/s. This measured distribution largely overlaps our simulated distribution, ranging from 0.9 to 45 m/s. Their study predominantly stimulated larger-diameter fibers and consistently failed to capture small myelinated axons, which likely explains the absence of the lower end of our velocity distribution in their results. Nevertheless, the conduction velocities obtained in our simulation did not reach the highest values reported by Firmin et al.. This discrepancy may reflect limitations in the biophysical realism of the ion channel properties used in our models, or the absence of sufficiently large-diameter axons required to achieve the fastest conduction velocities. More generally, the interpretation of conduction velocity measurements depends critically on histological estimates of axon diameter, which are known to be susceptible to preparation-induced artifacts that can systematically shift the observed diameter distribution [14].

### Supplementary Material C

#### Reconstruction of Neuron Models for Compartment Simulations

Reconstructions of cortical neurons, including the myelin distribution at the axon, were extracted from the H01 dataset. The pipeline presented creates a skeleton model of a neuron from a volume segmentation, first turning the segmentation into a surface mesh, then creating a skeleton representation of the mesh, including the diameter distribution over the neurites and finally turning the skeleton into unbranched cable representations. Various error sources originating from imaging and segmentation artifacts are accounted for within the pipeline.

First, the Neuroglancer tool was used to combine partial segmentation of a single neuron and annotate different information for use in the pipeline. A soma was selected manually in a specific cortical layer based on its centrality within the volume. Starting from there, every connected branch was investigated manually in a depth-first search-like (DFS) approach, where its end was investigated. If a connected component to the branch was found, it was added to the segmentation.

We used 5 annotations in the Neuroglancer tool:

1. The “volume ending” point annotation marked where a branch ended when the end of the volume was reached.
2. The “false ending” point annotation marked where a branch ended and cannot be further followed due to image artifacts.
3. The “merge” line annotation marked two branch segments that belonged together but were separate due to missing segmentation.
4. The “false merge” line annotation marked two branches that were falsely connected in the segmentation.
5. The “label” point annotation marked transitions between compartment types. Starting at the soma, it labeled each branch with apical dendrite, basal dendrite, or AIS. In the axon, the label changed when the AIS ended, the axon became myelinated or unmyelinated and when a node of Ranvier was encountered.

#### Creation of the False-Merge Mask

To solve the manually annotated false merges where the segmentation combined neurites that did not belong to the same cell, we used a max-flow min-cut algorithm implemented in the *preview_split* function in PyChunkedGraph[1] . This function split the segmentation into two and we used it to create a mask for the segmentation that removed the falsely merged segmentation.

#### Error Correction of the Volume Segmentation

To process the volume segmentation of the neuron, subvolumes with a size of 256 × 256 × 256 voxels and a voxel resolution of 32 × 32 × 33 $nm^3$ that contained voxels of given segmentation IDs were identified. A padding of 128 voxels was added to each subvolume in each dimension as an overlap between neighboring subvolumes, which resulted in a total subvolume size of 512 × 512 × 512 voxels. Five common errors in the segmentation were corrected:

1. Self-connections
2. Holes that are fully contained within the segmentation
3. Holes that have a connection to the background
4. Missing or corrupted slices
5. Unconnected, fully contained elements

The first common error, self-connections, needed to be addressed manually. When the segmentation connected parts that were not physically connected, the segmentation was manually edited in a voxel segmentation editor and corrected. In the first step of the automated error correction, *scipy.ndimage.binary_fill_holes* was used to fill all fully-contained 3D holes in the segmentation. It was then also used on every 2D slice in every 3D axis direction to fill holes

[1] https://github.com/CAVEconnectome/PyChunkedGraph

that were connected to the outside of the segmentation. In the second step, missing slices were detected and interpolated from valid neighboring slices. For every 2D slice in the z-direction, the inner and outer distances were calculated using *scipy.ndimage.distance_transform_edt*. The maximum inner distance was calculated for every slice, as well as the maximum inner distance of neighbors of each slice. The normalized maximum distance gradient was calculated based on the gradient of the maximum inner distance between neighbors. Slices with a gradient that was below −0.65 were identified as starting slices after which one or multiple slices were missing or were corrupted. Slices with a gradient above 0.65 were identified as ending slices, where a section with one or multiple missing or corrupted slices was ending. The gradient threshold at 0.65 was experimentally identified as a good trade-off between false identification and missed identification. The distance transform for slices between the starting and ending slices were then linearly interpolated between the distance transform of the starting and ending slice and thresholded at 0.5 to generate a binary segmentation for the missing or corrupted slices. In the third step, the first step was repeated to fill holes that might have been generated by the interpolation of missing or corrupted slices. Finally, unconnected, fully contained elements were connected to the rest of the segmentation. They were detected by labeling all unconnected structures using *scipy.ndimage.label* and identifying the structures that did not have any voxels on the border of the subvolume. To reconnect the element, the voxel coordinates that were closest between the element and the rest of the segmentation were identified. To connect the two coordinates with a straight line, *skimage.draw.line_nd* was used. The connecting line was dilated for two iterations using *scipy.ndimage.binary_dilation.* Using the same method, the annotated “merge” line annotations were integrated into the segmentation.

#### Volume Segmentation of the Soma

The soma as a special compartment of the neuron was best represented not by a cylinder like other neural compartments, but by a sphere. It also acted as a good root node for the neuron graph. To segment the soma, the center of the soma was estimated using the skeleton provided through the H01 dataset. The node with the largest diameter was a first estimate of the center of the soma. The volume segmentation centered around the soma center with a size of 156 × 156 × 151 voxels and a voxel resolution of 192 × 192 × 198 $nm^3$ was then processed. First, the same voxel segmentation error correction steps as for the other segmentation subvolumes were applied. Then, the integral inner distance was computed using *scipy.ndimage.distance_transform_edt.* A better estimation of the center was computed as the position with the maximum inner distance. To estimate where the soma ended and the neurites began, the thickness was calculated at each voxel using the fast local thickness method described by Dahl and Dahl [15]. The gradient of the thickness was smaller within the neurites and the strongest at the transition between the soma and the neurites. The gradient of the local thickness was calculated using *scipy.ndimage.gaussian_gradient_magnitude* with a Gaussian filter standard deviation of 2 to reduce noise in the gradient. To preserve the original edges, the gradient was masked with the original segmentation. Voxels with a gradient less than or equal to 1.2 were extracted. This separated the soma from the neurites, and finally, the soma was extracted using *scipy.ndimage.label* to identify the separated segment that contained the soma center coordinate.

#### Neuronal Surface Mesh Generation

For each error corrected subvolume of the segmentation with size 256 × 256 × 256 voxels and a voxel resolution of 32 × 32 × 33 $nm^3$, discarding the padding, an open surface mesh was generated. The marching cubes algorithm, which was implemented in *skimage.measure.marching_cubes,* was used to reconstruct the isosurface at an isovalue of 0.5. The resulting surface mesh was then downsampled to a target edge length of 64 nm using *CGAL::Polygon_mesh_processing::isotropic_remeshing* while preserving all border edges. All surface meshes resulting from all subvolumes were then stitched together using *CGAL::Polygon_mesh_processing::stitch_borders,* followed by another downsample step to a target edge length of 96 nm. All unconnected surface elements that consisted of fewer than 20 triangles were discarded. This threshold was experimentally identified as a good trade-off between removing invasive objects and preserving smaller, unconnected neural elements that can be reconnected in a later step.

#### Skeletonization of Neuronal Surface Meshes

Compartment simulations of neurons required modelling of neurons as a tree structure with meta information about the diameter of each tree node. The tree structure can be generated from a surface mesh using a TEASAR like algorithm [16] developed by extending the implementation in MeshParty[2]. The mesh was smoothed using Taubin smoothing [17] implemented in *pyvista.PolyDataFilters.smooth_taubin* with 10 iterations and a pass band of 0.1. The connected mesh components were represented as a mesh graph where the mesh vertices were the graph nodes and the edges between vertices had weights calculated as the Euclidean distance. Shortest-path graph searches were performed using *sparse.csgraph.dijkstra*. All connected mesh components were separately skeletonized and connected afterwards. The component containing the soma had its root set to the surface node that was closest to the center of the soma, and the nodes that were part of the soma were invalidated using the soma segmentation. For all other components, the root was selected as the node furthest away from the node with index 0. This set the root to one of the "ends" of the component, and only the root node was invalidated.

The skeletonization finished when all graph nodes were invalidated. For each step, the node furthest away from the root node on the graph was identified. To connect this node to the rest of the skeleton, the shortest path from each skeleton node was calculated, and the shortest path was then added to the skeleton. This newly added path was referred to as the skeleton segment in the following.

Each graph node that was part of the same neurite as the skeleton segment needed to be invalidated, and at the same time, nodes on bifurcations coming off this neurite needed to be avoided. Each skeleton segment node was assigned the direction vector to the next skeleton segment node. The direction vectors were smoothed along the path using *scipy.ndimage.gaussian_filter1d* with a standard deviation for the Gaussian kernel of 1. Each node of the graph was assigned the direction of the closest skeleton segment. Two new directed graphs were constructed: the orthogonal directed graph and the parallel directed graph. The orthogonal directed graph was constructed by calculating for each directed edge the cross product of the surface normal of the edge and the closest path direction to receive the

[2] https://github.com/CAVEconnectome/MeshParty

orthogonal direction. The orthogonal direction was the direction that pointed in the direction around the neurite. Then, only directed edges in the graph that were less than 90° to the surrounding direction were kept, determined by the dot product between the surrounding direction and the edge direction. All edges that were included in the skeleton segment were removed. The Euclidean edge weights were weighted by the dot product. The parallel directed graph was constructed by calculating for each directed edge of the graph the dot product between the closest path direction and the edge direction, and only allowing edges that were less than 90° to the closest path direction.

For the first and last node of the skeleton segment, a surrounding, non-trivial cycle was computed by finding the shortest path for each node on the graph to the skeleton segment on the orthogonal graph and the inverse orthogonal graph. If the closest skeleton node was the same on the orthogonal graph and the inverse orthogonal graph, a non-trivial cycle around the neurite was found. The shortest cycle of this kind was chosen for the first and the last skeleton segment node. Nodes of the graph that were part of the neurite that the skeleton segment was on could now be identified as nodes that had a connecting path on the parallel graph to the ending cycle and on the inverse parallel graph to the initial cycle. A binary closing operation with a closing range of 500 nm was then performed on these nodes to avoid uninvalidated islands that did not represent a bifurcating neurite.

For each invalidated graph node, the skeleton segment ID that invalidated it, and for each skeleton segment node, whether the node was already invalidated at the start of this iteration, were stored. This information was used in the diameter estimation and the creation of the neuron model.

The next step of the skeletonization was to estimate the diameter around the neurite as well as the center position of the neurite for each skeleton node. The same method used for the invalidation procedure was applied to compute non-trivial surrounding cycles around the neurite at each skeleton node. The diameter was calculated from the circumference of the cycle, and the center was estimated as the mean coordinate of all cycle nodes. In the case that no such cycle could be found for a skeleton node, the diameter and center coordinate were linearly interpolated from the neighboring nodes.

In the last step, all skeletons resulting from the connected components of the neuronal surface mesh were connected. For each skeleton that was not connected to the soma, the pair of points that were the closest between the points of this skeleton and the points of all other skeletons was found and connected. This was done until all skeletons were connected to one skeleton.

#### Skeletonization of the Soma

To estimate the soma volume as a spheroid and generate a realistic skeleton, the volume segmentation of the soma was used. From the volume segmentation, the center of the soma was estimated as the mean of all soma voxel coordinates. The principal direction was calculated using the principal component analysis implemented in *numpy.linalg.svd.* The center and the principal direction were used to create the center axis of the soma by tracing along the principal direction and the inverse principal direction in fixed steps starting at the center and ending when the axis left the soma volume. For each point on the center axis, evenly sampled rays were shot perpendicular to the axis to find the border points surrounding that point. The radius of the soma surrounding each point was then calculated as the mean distance between the center and the intersection points.

#### NEURON Model Generation from Skeleton

The generation of neuron models that can be used in the simulation environment NEURON started with manually resolving errors in the skeleton. Diameter and center coordinates that appeared to be outliers or where the diameter was unknown (e.g., where the skeleton was created based on a “merge” line annotation) were removed and linearly interpolated from neighboring nodes of the skeleton. Misplaced edges were removed or replaced by manually determined edges.

The pre-computed center axis of the soma was integrated into the skeleton by removing all skeleton nodes that were within the soma volume segmentation and reconnecting the dangling skeleton nodes to the center of the soma axis. The nodes connected to the soma axis were marked as virtual edges. Virtual edges were used only for connectivity purposes, but did not represent geometrical features of the neuron. They were used in connecting neurites to the soma and bifurcation branches to the main branch. The diameters of the soma axis were then smoothed along the axis using graph-based Laplacian regularization with an alpha value of 50.

In the next step, the skeleton was centered in the neuron. To reduce over- and underestimation of the diameter and the center coordinates, they were median filtered along the skeleton with a neighborhood of five nodes. The skeleton was then centered by replacing the skeleton node coordinates with the center coordinates. Within the skeletonization, skeleton nodes were marked based on the invalidation state of the node when it was added to the skeleton. The nodes that were invalidated before they were added to the skeleton were now removed from the skeleton, and the disconnected branches were reconnected using virtual edges. The traversal of the skeleton starting at the soma was performed using the depth-first ordering implemented in *scipy.sparse.csgraph.depth_first_order.* To reduce noise in the skeleton node coordinates, all nodes except the soma axis and nodes that were part of virtual edges were smoothed using graph-based Laplacian regularization with an alpha value of 100.

The biophysics of different neurites were highly distinct. To correctly apply them, the nodes in the skeleton needed to be labeled. For neurons, six groups of compartment types were applicable: axon, dendrite, soma, AIS, myelinated axon and node of Ranvier. The soma label was applied based on the volume segmentation of the soma. The “label” point annotations were projected to the closest skeleton points, excluding the already labeled soma points. Each neurite starting at the soma was labeled by the closest point annotation. A DFS traversal through the skeleton, starting at the soma root node, marked the skeleton points based on the last label that was encountered.

To prevent stimulation effects from where neurites were cut due to the volume ending or artifacts, the “volume ending” and “false ending” point annotations were projected to their closest skeleton leaf nodes. If a myelinated axon was cut, a straight segment was added pointing in the same direction as the pre-cut axon. The straight part consisted of a continuation of the internode by a node of Ranvier and finally followed by a 1 mm long passive axon. The length of the internode and the node of Ranvier were based on the internodes previous to the cut internode. The same approach was applied when an axon was cut at a node of Ranvier or when it was cut closely after myelination ended on an axon.

To properly represent each section separated by virtual edges or defined by connected nodes of the same label, they needed to consist of at least two nodes to represent their direction. To resolve this, new nodes were added. A list of sections was generated by traversing the skeleton starting at the soma, performed using depth-first ordering implemented in *scipy.sparse.csgraph.depth_first_order.* A new section began either when the label changed or after a virtual edge. For each section consisting of a single node, a node splitting the previous edge leading to the node was added. If the previous edge was a virtual edge, the previous edge was split by adding a new node close to the section node. Otherwise, the previous edge was split by adding a node in the middle of the edge.

At terminations, the skeleton often wrapped around the termination site. To preserve the realistic direction of the termination, all termination sections consisting of at most 15 points before the termination were processed. The direction vector of each node pointing to the previous node was computed, and they were then used to calculate the angle between different subsections starting from the termination of the section. If the maximum angle on the termination section exceeded 38°, the section was identified as a bend termination. These sections were then corrected by replacing the part after the bend with a straight prolongation of the unbend part. The diameter was corrected by tapering the diameter of the last 3 nodes linearly down to 200 nm.

The skeleton was now rotated so that its manually determined somatodendritic axis was aligned with the *z*-axis, which will allow the neuron model to be placed inside cortical layers accordingly.

The last step was to turn the skeleton with its labels into a section-based format. Each section was defined by its points and their diameter, the label, and the fraction along its parent where it was connected. The skeleton node coordinates and node diameter were used to define the sections. The soma was a single section without a parent and served as the root. All sections connected to the soma were connected to the center of the soma, so the fraction along their parent is 0.5. All other sections were connected to the end of their parent sections at 1.0. The parent relationship was defined by traversal of the skeleton starting at the soma using the depth-first ordering implemented in *scipy.sparse.csgraph.depth_first_order.* A section ended either when the label changed, before a virtual edge, or when the path ended. The biophysics were applied according to the section labels.

**Table S-C1:** The biophysics models used for each neuron extracted from the H01 dataset. The biophysics models were named as originally given by Markram et al. [1]. The cell ID corresponds to the segment ID of the segment containing the soma of the neuron in the H01 dataset [18].

| Layer | Cell ID | Biophysics Model |
|---|---|---|
| II | 3970566758 | L23_PC_cADpyr |
| II | 2571514217 | L23_PC_cADpyr |
| III | 2962791579 | L23_PC_cADpyr |

| | | |
|---|---|---|
| III | 3269032756 | L23_PC_cADpyr |
| IV | 4696659954 | L4_PC_cADpyr |
| IV | 3325304783 | L4_PC_cADpyr |
| V | 3338693880 | L5_TTPC2_cADpyr |
| V | 33067397116 | L5_TTPC2_cADpyr |
| VI | 4443042188 | L6_TPC_L4 |
| VI | 31156125745 | L6_TPC_L4 |

### Supplementary Material D

#### Morphology of Biological Neuronal Components

The following sections analyze an extensive body of histological studies about the complex cortical axon morphology. An in-depth summary of methods and reported values is shown in Supplementary Material I.

##### Morphology of Myelinated Axon

The axon diameter *D* of cortical primate neurons is highly variable. Studies across many brain regions have described its distribution as strongly skewed towards small diameters (0.1–1.0 µm), with a long tail representing a small number of larger axons [13,19–24]. The diameter depends on both the origin and the target of the projection [21]. Across cortical regions, mean diameters increase from prefrontal (0.43–0.55 µm) [19,24] to premotor (0.57–0.66 µm) [19] and motor areas (0.72–1.02 µm) [19,21]. This pattern primarily reflects a higher proportion of larger axons (> 1.2 µm), while the modal diameter remains similar across regions [19]. Large axons (> 3 µm) have been reported in varying quantities originating from or terminating in the prefrontal cortex [24], motor cortex [13,19,21,22], somatosensory cortex [21] and in SWM tracts (BA 3, 4, and 18) [12]. The largest axons in the brain were measured in the medullary pyramid (~10 µm) [13,22], likely originating from the largest Betz cells of the motor cortex [13]. Exceptionally large and rare fibers are also found in the deep white matter (WM) of the prefrontal area (7 µm) [24], in the corpus callosum (7.48 µm) [12], in the SWM (7.073 µm) [12] and in the superior longitudinal fasciculus (9 µm) [23]. Because small-diameter axons vastly outnumber larger ones in most areas, sampling bias likely underestimates the true prevalence of large fibers. Based on the literature, we therefore considered axon diameters within the range of 0.2–10 µm.

The myelinated axon is organized into internodes, separated by nodes of Ranvier. The internode length *L* correlates with the axon diameter [25–28], but shows a wide variation, especially for larger *D* in the rats' anterior medullary velum (AMV) [25,26]. The relationship *L*/*D* also varies significantly between different parts of the mammalian CNS [8–10]. It is influenced by distance to terminals and branching points [27] and is age-dependent [29]. Individual fibers have been shown in rat AMV to have less variation in *L* [26]. On average, *L*/*D* is between ~30 and ~120 in the rodent auditory brainstem [27] and the AMV [25,26], with different region-specific intercepts. Assuming that the synthetic models resemble axons far from branching points and terminals and given that individual fibers show less variation [26], we approximated synthetic axons with a stable *L*/*D* for individual fibers between 60 and 120.

In contrast to *L*, the node of Ranvier length *N* does not appear to correlate with *D* in the rat motor cortex, and its variation is greater between axons than within individual axons [30]. In rodents, *N* has been reported to range from 0.1 to 4.4 µm in the corpus callosum [31], with similar ranges in the prefrontal cortex WM [32] and motor cortex layer V [30]. The mean was stable across the different areas, with a value of ~1.5 µm. In humans, however, mean *N* is higher, at approximately ~2.5 µm in prefrontal cortex WM [32]. Considering both interspecies differences and the broad range of reported values, we assumed a stable node of Ranvier length for individual synthetic fibers between 0.5 and 5 µm.

#### Morphology of the Soma and Axon Initial Segment

Reported soma diameters *S* in pyramidal neurons range from 9–33 µm in humans and macaques [19,33,34], with giant Betz cells reaching 27–98 µm in the human motor cortex [34–36]. Axon diameters span 0.25–1.3 µm in human and macaque pyramidal cells [19,33], but can reach ~10 µm in the medullary pyramid [13,22], likely originating from giant Betz cells. A linear relationship between the soma and axon diameter *D* has been reported in humans and macaques [19,33]. The AIS length *K* ranges from 8–85 µm in human cerebral and neocortical pyramidal neurons [37,38], 31–54 µm in macaque motor and somatosensory cortex [39] and 11–73 µm in pyramidal neurons of the mouse visual cortex in organotypic tissue cultures [40]. Its length appears independent of cell size in macaques [39]. A gap between the axon hillock and AIS *G* has also been reported: 0–7.5 µm in humans [38] and 1.5–53 µm in mouse visual cortex [40].

#### Morphology of Axonal Bifurcation

Bifurcations of myelinated axons have been reported in both cortical and subcortical regions. Cortical interneurons exhibit extensive branching, but their axons are typically thin and only partially myelinated, as observed in both humans and mice [41]. In contrast, pyramidal cells can elicit thick horizontal, likely myelinated major collaterals from their main axon. These major collaterals have been shown to exist in layer III and IV pyramidal cells in monkeys (BA 4, 3a, 3b, 1, and 2) [42,43]. Consistent myelination can not always be expected, as patchy myelination has been found in interneurons and pyramidal cells, especially in the initial area of the axon in monkeys, mice and humans [41,44–47]. In the SWM of monkeys in various areas, likely myelinated axons have been shown to bifurcate, where the main branch stays inside the SWM while the collateral branches into the cortex [42,43,48,49]. Further branching of likely myelinated projecting axons within the cortex has been observed in monkeys [43,48–50]. There is also branching in the deep WM of monkeys below the motor cortex [51], albeit less relevant due to the weak TMS-induced E-field in this area. According to Chklovskii's power law, axonal diameters at bifurcations scale predictably, although the empirical data show substantial variability [52]. A general shrinkage of collaterals at bifurcations has also been shown in mouse interneurons [41].

#### Morphology of Axon Terminals

The terminals of cortical axons are predominantly located in the gray matter (GM) of the brain, where dendrites are present. In this region, axon diameters *D* have been measured in the range of 0.275–1.3 μm in humans [33] (BA 17, 20, and 21) and macaques [19] (BA 46 and 4). Studies in basket cells in mouse visual cortex [44], macaque somatosensory cortex [53], and cat visual cortex [54] show that axons become unmyelinated prior to their terminal connections. In pyramidal cells, tracing studies in monkeys show that terminal arbors are thin [42,43] and highly branched [42,43,49]. Because myelination probability in human and rat pyramidal cells and interneurons depends on axon diameter and interbranch distance [41,45], such terminal arbors are therefore less likely to be fully myelinated. Reported myelin internode lengths *L* range from 16.9–137.8 μm in pyramidal cells and interneurons in humans [41,45] and mice [44,46]. In addition, a general, diameter-unrelated decrease in internode length close to terminals has been shown in the auditory brainstem of rodents [27].

#### Morphology of Axonal Bend

Sharp bends of myelinated axons were observed in monkeys across multiple brain regions. In deep WM below the motor cortex, myelinated axons make abrupt turns with bend radii *R* of 13–17 μm [51], even though they are likely less relevant for TMS due to the weak E-fields induced at those positions. Within the cortex, myelinated axons that project perpendicular to the pial surface can form “elbow-shaped” bends as they enter Exner’s plexus (layer I), with reported radii of 40.6, 25.75, and 23.35 μm [50]. Additional sharp bends occur where association fibers enter or leave the SWM, changing orientation from perpendicular to parallel relative to the GM–WM boundary. These have been reported between areas V1 and V2 [49] and BA 2, 3a, and 4 [42,43], with radii of 3.05–98.96 μm. In contrast to these sharp turns, smooth bending profiles have also been documented in monkeys. Long-range intracortical collaterals that follow the folding pattern of the cortex exhibit bend radii of 255.3 μm between BA 4a and 3a [42] and 461.18 μm between BA 1 and 2 [43]. Similarly, association fibers (U-fibers) that also follow the cortical folding structure but in the SWM have been found with bend radii of 1596.9 μm between BA 4a and 3b [42] and 592.12 μm between BA 3a and 3b [43]. Lastly, projection fibers can bend smoothly as they descend into deeper WM. The main axons of layer III and layer V pyramidal neurons from M1 were measured in the range of 788.7 and 1560.8 μm [55].

### Supplementary Material E

#### Simulation of Model Time Constants

The membrane time constant is a well-established measure of the speed of passive voltage change in a neuron, reflecting how quickly the membrane potential both rises and decays in response to a change in current input [56]. In the equivalent RC circuit, the membrane time constant ($\tau_m$) is described by the product of the membrane resistance $\left(R_m\right)$ and the membrane capacitance $\left(C_m\right)$

$$\tau_m = R_m \cdot C_m. \tag{E1}$$

Experimentally, the time constant can be estimated using the strength-duration relationship. By applying pulses of different durations to elicit action potentials, an exponential strength–duration relationship between pulse duration and stimulation intensity can be derived, from which the strength-duration time constant (chronaxie) and the rheobase are obtained. The rheobase is the stimulation intensity needed to evoke an action potential with an infinite pulse duration, whereas the strength-duration time constant is the minimal pulse duration at twice the rheobase.

For TMS, in vivo strength-duration relationships have been estimated using cosine pulses with different rise times ($\tau$ = 152 ± 26 μs) [57] and using cTMS pulses with square-wave-like waveforms ($\tau$ = 200 ± 33 μs) [58,59]. In our simulations, we estimated an effective membrane time constant and rheobase using the same strength-duration fitting approach as in previous studies [59–61]. Specifically, we simulated activation thresholds for three cTMS waveforms (30, 60, and 120 μs pulse duration) and fitted a theoretical strength-duration model to these simulated data. A single low-pass filter with a time constant $\tau_m$ was used to approximate the neural membrane. For a given pulse duration $t_p$, rheobase $\Theta_{rh}$ and depolarisation factor $r(\tau_m, t_p)$, defined as the peak of the low-pass filtered cTMS waveform, the predicted threshold is:

$$\widehat{\Theta}(t_p) = \frac{\Theta_{rh}}{r(\tau_m, t_p)}. \quad \text{(E2)}$$

The time constant and rheobase were estimated by least-squares fitting, minimizing the sum of squared relative errors $\left(\frac{\widehat{\Theta}}{\Theta} - 1\right)^2$ between the predicted thresholds and the simulated thresholds across the three pulse durations.

Fig. S4 shows the time constants and rheobases estimated for the four synthetic axon models. We excluded cases of mixed membrane properties, which arise when part of the axon was unmyelinated, because a single first-order low-pass filter cannot accurately represent multiple distinct membrane time constants. We also excluded fits yielding time constants greater than 1000 μs, as these values were outside the physiologically plausible range and typically reflected poor fits. For the bend model, we further excluded simulations with thresholds above 500 V/m, where activation occurred only at unrealistically high field amplitudes. Across all four models, the fitted time constants were slightly higher than the range reported in experimental TMS studies. In the bend axon model, thicker axons produced time constants that were closer to the experimentally estimated range than those of thinner axons. However, the rheobase cannot be directly compared to experimental estimates without field modeling using subject-specific head models to convert the percentage of maximum stimulator output reported in experiments into cortical field amplitudes.

Fig. S5 shows time constant and rheobase estimates for smoothly bending axon projections. We observed that the time constants were in a broadly realistic range (100 to 300 μs) in areas of low threshold (compare with PA results from Fig. 8 in the main text), which agreed with the results from Fig. S4D, specifically, the axons with bend radii of 1000 μm and bend angles of 45°-90°. In areas of high thresholds, the values were unrealistically high and the residuals were high as well, suggesting that the fit failed in these regions.

### Supplementary Material F

#### Creation of the Anisotropic Conductivity Distribution

To represent the conductivity distribution in gray and WM, we combined diffusion weighted image-based reconstruction of WM anisotropic conductivity with a histologically informed spatial smoothing of the GM-WM transition zone.

WM conductivity tensors were reconstructed from diffusion weighted images using the volume-normalized mapping approach implemented in SimNIBS 4.6 (dwi2cond). GM conductivity was modeled as isotropic with a magnitude of 0.275 S/m. All conductivities were defined on the tetrahedral volume mesh within a high-resolution region-of-interest (ROI) around the M1 region. Histological data from the H01 dataset indicated that the transition zone spanned approximately from the center of cortical layer VI to a comparable depth into WM [18]. To approximate this boundary, we constructed a surface representation of cortical layer VI at a normalized cortical depth of 85% [62] using the equivolume approach [63,64] as implemented in the cortech package[3].

A face adjacency graph was constructed on the tetrahedral mesh restricted to the ROI. Two tetrahedra were connected if they shared a face and both belonged to the ROI. Let $i$ and $j$ denote two connected tetrahedra with barycenter locations $b_i$ and $b_j$. Edge weights were defined as a Gaussian affinity

$$w_{ij} = exp(-\frac{\left\|b_i - b_j\right\|^2}{0.1^2}), \tag{F1}$$

where ||·||denotes the Euclidean norm and distances were measured in millimeters. To reflect the transition zone between gray and WM, we defined a spatially varying smoothing strength $\zeta_i$ based on the distance to the WM surface. A per-vertex proxy $t_{surf}$ was computed as the Euclidean distance between corresponding vertices of the layer VI central surface and the WM surface. Each ROI tetrahedron barycenter was mapped to its nearest white surface vertex. Let $e$ denote the barycenter to WM surface distance. The local smoothing parameter was then defined as

$$\zeta_i = max(t_{surf} - e,\ 0) \cdot 200. \tag{F2}$$

The Gaussian width parameter of 0.1 mm and the smoothing strength scaling factor of 200 were chosen based on empirical validation to enforce locality of the smoothing.

Conductivities were smoothed by solving, independently for each tensor component $\sigma_i$, the screened Laplacian system

$$(P\ +\ diag(\zeta)A)\sigma_{is} = \sigma_i, \tag{F3}$$

where $P$ was the identity matrix, $A = F - W$ was the graph Laplacian, $W$ was the symmetric affinity matrix with entries $w_{ij}$ and $F$ was the diagonal degree matrix. This formulation resulted in

[3] https://github.com/simnibs/cortech

stronger diffusion of conductivity values near the cortical boundary and weaker diffusion further away, stabilizing the transition zone while preserving anisotropy in deeper WM.

We compared stimulation thresholds for sharp and smooth bends embedded in the M1 region using standard isotropic conductivities (GM 0.275 S/m, WM 0.126 S/m) and the proposed histologically informed anisotropic model. For regions of low activation threshold, the threshold differences between the two models for sharp bends were generally small, with slightly lower thresholds with isotropic conductivities. In contrast, for smooth bends that run deeper in the WM, smoothed anisotropic conductivities yielded lower activation thresholds (Fig. S6).

### Supplementary Material G

#### Smooth Bending White Matter Axon Projections

Deep WM axon projections have previously been modeled using different approaches, most of which involve defining a vector field (in 2D or 3D) on which streamlines can be traced. For example, Nummenmaa et al. [65] assume that fiber orientation is perpendicular to the cortex in GM and use the first eigenvector from diffusion tensors in WM, whereas Gomez-Tames et al. [66] solve the Laplace equation in the GM compartment and use control points in WM to achieve different bend profiles. Here, we obtained the vector field by solving the Laplace equation on the entire 3D domain (WM and GM compartments).

##### Generating the domain

To solve the Laplace equation on a realistic domain, we used the finite element method (FEM). We cut out parts of the white and GM cortical surfaces corresponding to the M1 ROI and proceed as follows to generate an enclosed domain. A “deep WM boundary” was created by shrinking the WM surface towards its center. In particular, we applied mean curvature flow exclusively to points of positive curvature. For concave areas, we moved points slightly inwards in the direction of the anti-normal of the surface. This ensured that the surface was strictly shrunk. To close the domain, we triangulated the sides between deep WM and WM and between WM and GM and applied fairing to obtain a smooth outline. Next, we tetrahedralized the interior such that the WM–GM boundary was preserved. Finally, we remeshed the domain using MMG[4]. The entire pipeline is shown in Fig. S7.

##### Generating the vector field

To estimate the potential field on the domain, we solved the Laplace equation using FEM. We set Dirichlet boundary conditions on the deep WM (0 V) and GM (1000 V) surfaces. Finally, we differentiated the estimated field to obtain a vector field (the E-field). However, setting a single diffusivity value on the entire domain would result in streamlines that bend too smoothly, as all streamlines would converge on the centerline of a gyrus. On the other hand, if we assigned different diffusivities to the inner and outer compartments (e.g., WM and GM), we could manipulate the smoothness of the streamlines around the transition from one compartment to another, effectively achieving different bend radii. One complication was that we did not know exactly what bend radii were physiologically realistic. However, we were able to gather and estimate the following values from the literature:

- We made manual measurements using ImageJ on Fig. 4 of Shapson-Coe et al., showing the myelinated axons of the H01 dataset [18]. First, the scale was set based on the scale bar of the figure. Then, a circle with a constant radius was manually fitted to the bend, and the diameter was measured using the measuring tool. We obtained bend radii between 733 and 812 μm.

[4] https://www.mmgtools.org

- Similarly, we estimated the bend radius of axon trajectories published by Li et al. [55] for neurons in the M1 of a macaque monkey. We found a mean bend radius of 1217.3 (± 210.4) µm with a range of 788.7–1560.8 µm.
- Cottaar et al. [67] estimate a width parameter for a sigmoid function that describes fiber orientation transition in diffusion weighted imaging. From this, we calculated bend radii ranging from 710 to 2120 µm with an estimated bend radius of 1410 µm for approximately 90° of axonal bending. In the following, we describe how these bend radii were obtained in more detail.

Given these values, we aimed to generate axon projections with bend radii of approximately 1200–1410 µm at sulcal walls (~90° bends), which we found that a diffusivity ratio of 0.2 (GM/WM) achieved (Fig. S8).

#### Estimation of bend radii

The estimations from Cottaar et al. [67] were obtained in the following way. In the paper, they report a width parameter (in mm) of a sigmoid function that describes the transition of fiber orientation (radial and tangential) across the WM-GM boundary. A sigmoid function was fitted per voxel using high-resolution diffusion weighted imaging data, thus they provided a brain map of such width parameter values. Here, we transformed this parameter to a bend radius by integration. In particular, we assumed that the logistic function describes the proportion (between 0 and 1) of the radial vector component

$$\varsigma(x) = \frac{1}{1-exp(-x)}. \tag{G1}$$

Then the parametric representation

$$y' = (\varsigma(t),\ 1 - \varsigma(t)) \tag{G2}$$

described the mixing of radial ($\varsigma(t)$) and tangential ($1 - \varsigma(t)$) vectors in the fiber and therefore the direction of a fiber at a given point, with the parameter *t* denoting where on the curve the point is located. Consequently, it defined the unit tangent vector of another function,

$$y = \left(\int \varsigma(t)dt,\ \int 1 - \varsigma(t)dt\right), \tag{G3}$$

which described the actual path of the fiber. The curvature $u(t)$ of this fiber path was given by the norm of the second derivative of $y$ (i.e., the derivative of the logistic function)

$$u(t) = |y''(t)|, \tag{G4}$$

where

$$y'' = \left(\frac{d}{dt}\varsigma(t),\ \frac{d}{dt}(1 - \varsigma(t))\right), \tag{G5}$$

and the corresponding bend radius is

$$R(t) = \frac{1}{u(t)}, \tag{G6}$$

with the center of the osculating circle at

$$C(t) = y(t) + \frac{y''(t)}{u(t)^2}. \tag{G7}$$

By estimating $y''$, we obtained bend radii listed in Table S-G1 from realistic width parameter values found in Fig. 8 in Cottaar et al. [67].

**Table S-G1:** Estimation of bend radii based on the width parameter reported by Cottaar et al. [67]

| Width parameter (mm) | Bend radius *R* (μm) |
|---|---|
| 0.25 | 710 |
| 0.50 (~90°) | 1410 |
| 0.75 | 2120 |

#### Generating axon projections

The axon projections were generated by tracing streamlines on the vector field. Seed points for the streamlines were obtained by moving the vertices of the pial surface in the anti-normal direction by 0.5 mm. We followed the streamlines by integrating the vector field using a forward Euler scheme with a step size of approximately 10 V. The streamlines terminated on the deep WM surface.

To evaluate whether the bending characteristics of generated streamlines agreed with the values reported above, we needed to estimate the bend radius (or curvature) for each point on a fiber projection. This estimation was performed by fitting a 2nd degree polynomial to this point and its immediate neighbors. Thus, we fitted three parameters to three data points. Given these parametrically defined curves, we calculated the curvature as

$$u(t) = \frac{|y'(t) \times y''(t)|}{|y'(t)|^2}, \tag{G8}$$

and the bend radius *R* was given by (G6).

### Supplementary Material H

#### Biophysics Model

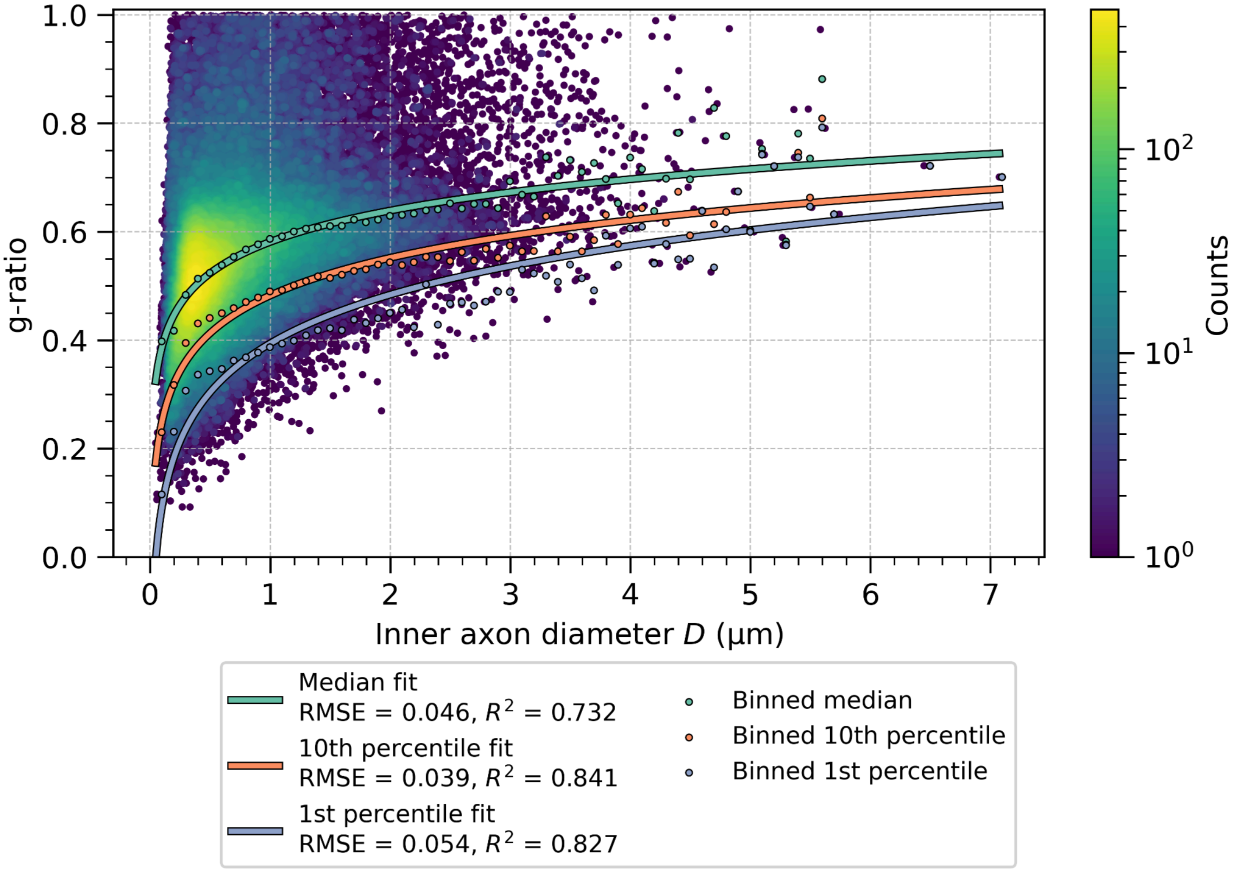


**Figure S1:** Relationship between the inner axon diameter and the g-ratio measured by Ruthig et al. [12] below and in between M1 and S1, as well as within V2. Three logarithmic fits are shown alongside the measured g-ratios: the median fit $(0.581 + 0.0834 * ln(x))$, the 10th percentile $(0.482 + 0.1002 * ln(x))$ and the 1st percentile $(0.393 + 0.1299 * ln(x))$. The root mean square error (RMSE) and the coefficient of determination ($R^2$) are listed in the legend.

### Simulation of Conduction Velocity

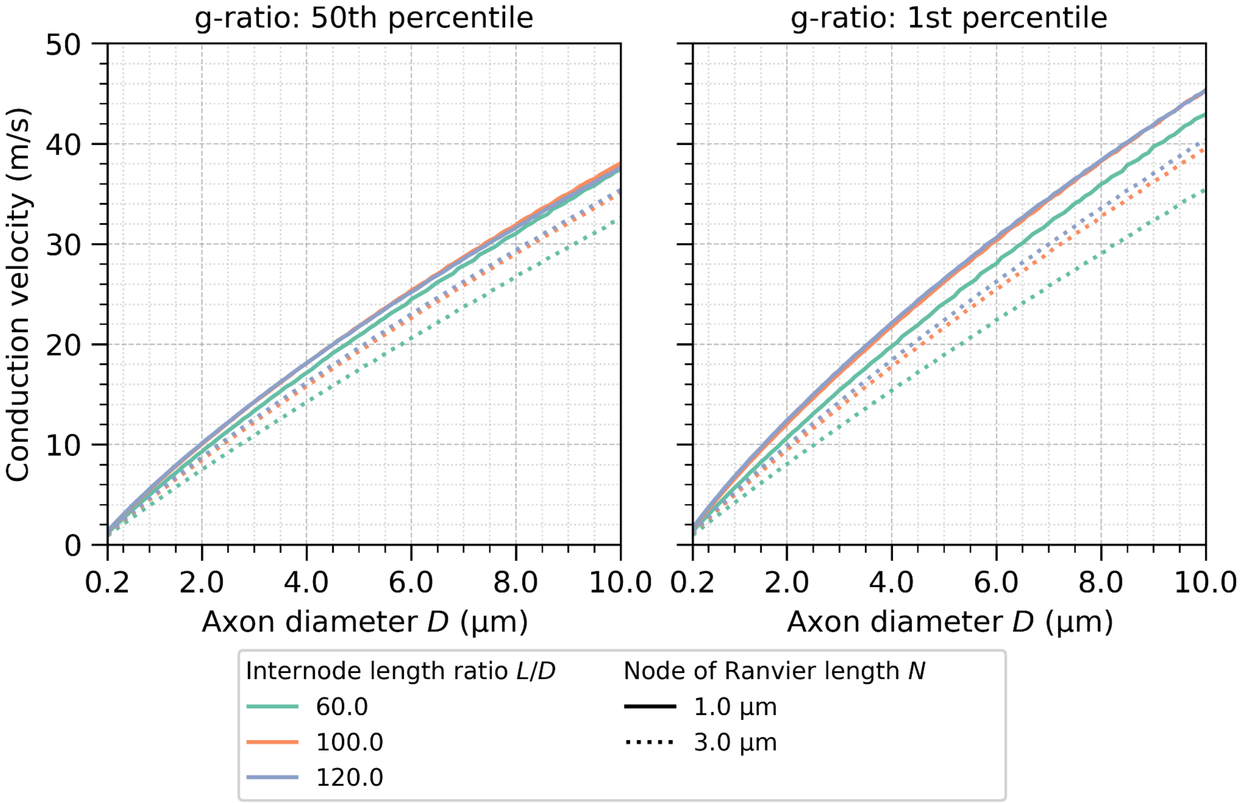


**Figure S2:** Conduction velocity of myelinated axons modeled using "L5_TTPC2_cADpyr" biophysical properties, evaluated across varying internode length ratios and node of Ranvier lengths. Results are shown for axons with g-ratios corresponding to the median fit (left) and the 1st-percentile fit (right) (see Supplementary Material A and Fig. S1).

Effects of E-field Alignment on Model Thresholds

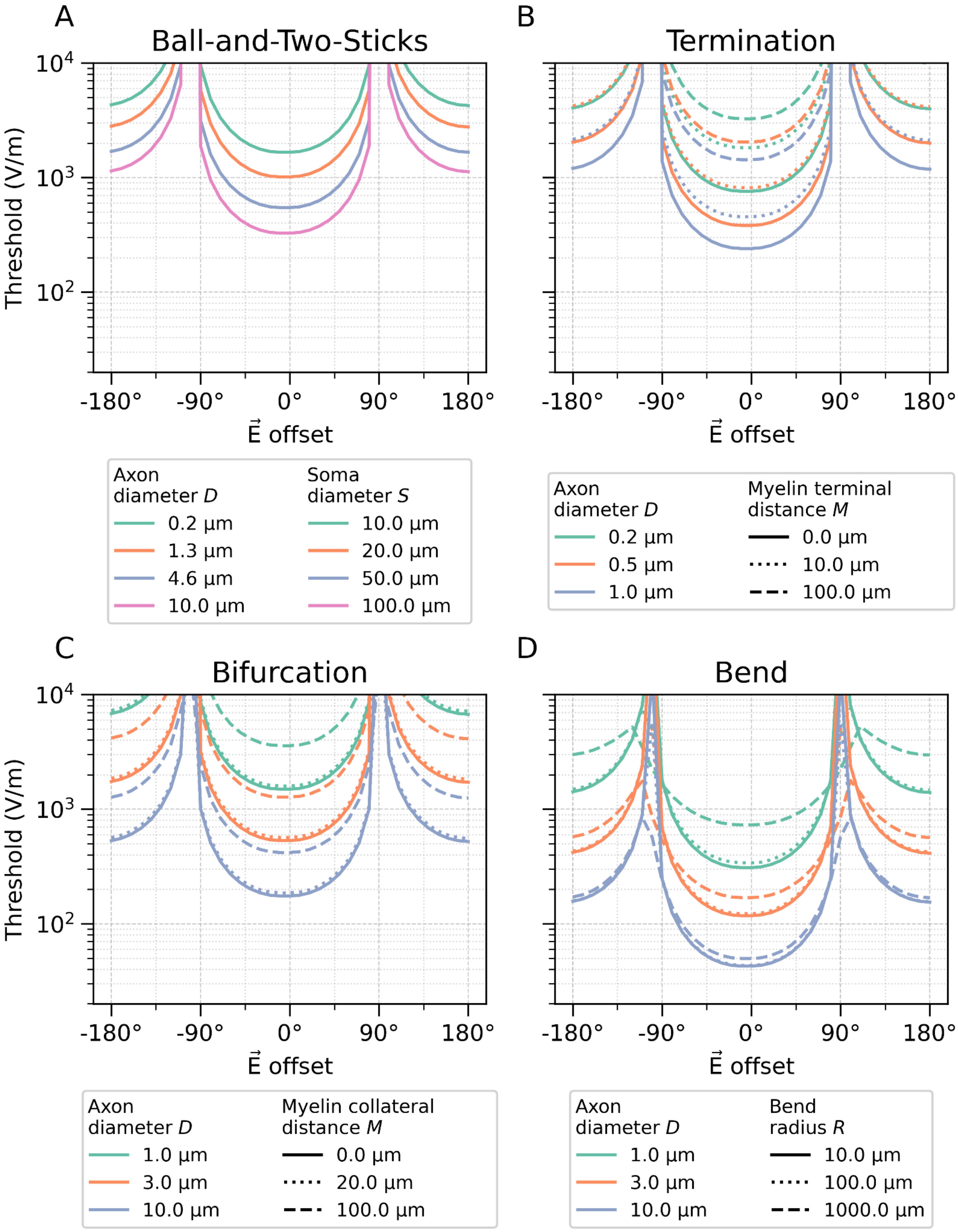

**Figure S3:** Relationship between the offset ($\alpha$) of the E-field from the optimal field direction (0°) and the activation threshold for (A) the "ball-and-two-sticks" model, (B) the terminal model, (internode length is 100 μm) (C) the bifurcation model (bifurcation angle is 30°), and (D) the bend model (bend angle is 90°). For all panels, the nodes of Ranvier has a length of 1 μm. In (A), the AIS has a length of 30 μm and there is a 9 μm gap between the soma and the AIS. For panels C and D, the internode length ratio (*L*/*D*) was 100. In all models (A–D), the threshold increased with angular offset according to the expected cosine dependence $E_s = E \cdot s = |E|\, cos(\alpha)$, which reflects the projection of the E-field $E$ onto each segment direction represented by unit vector $s$. In the interval [−180°,−90°) ∪ (90°,180°], the activation is driven by the negative second phase of the monophasic waveform, which acts as a long but weak pulse due to the coil's inductive rebound after the main positive phase.

Simulation of Model Time Constants

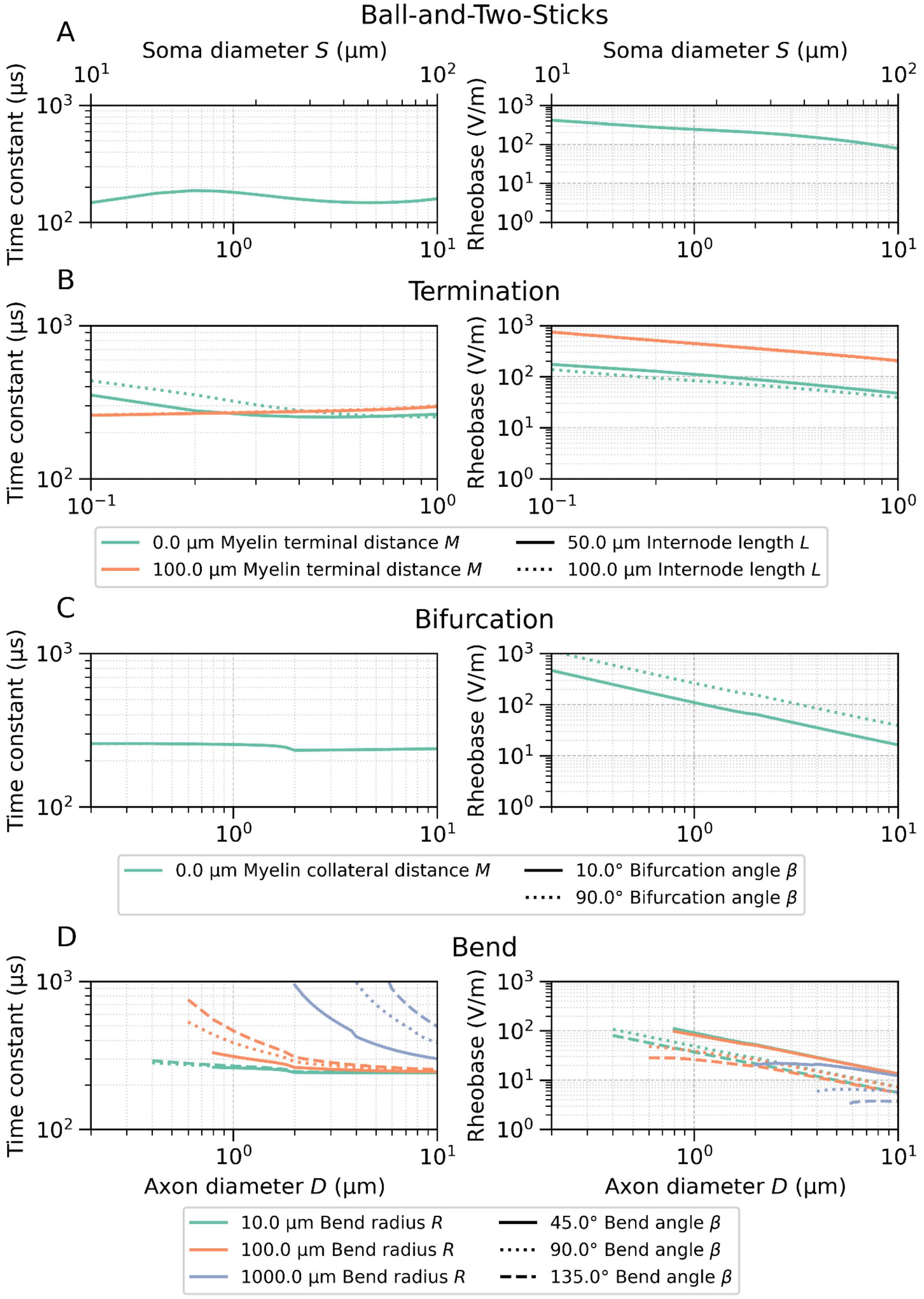

**Figure S4:** Relationship between model parameters and the time constant and rheobase for (A) the "ball-and-two-sticks" model, (B) the terminal model, (C) the bifurcation model, and (D) the bend model. We are only showing parameter combinations with consistent effective membrane properties. For panel D, we are only showing physiologically plausible time constant and rheobase estimates for simulations with a maximum activation threshold of 500 V/m and time constants below 1000 μs. In panel A, the AIS has a length of 30 μm and there is a 9 μm gap between the soma and the AIS. For panels B–D, the nodes of Ranvier had a length of 1 μm. For panels C and D, the internode length ratio (*L*/*D*) was 100.

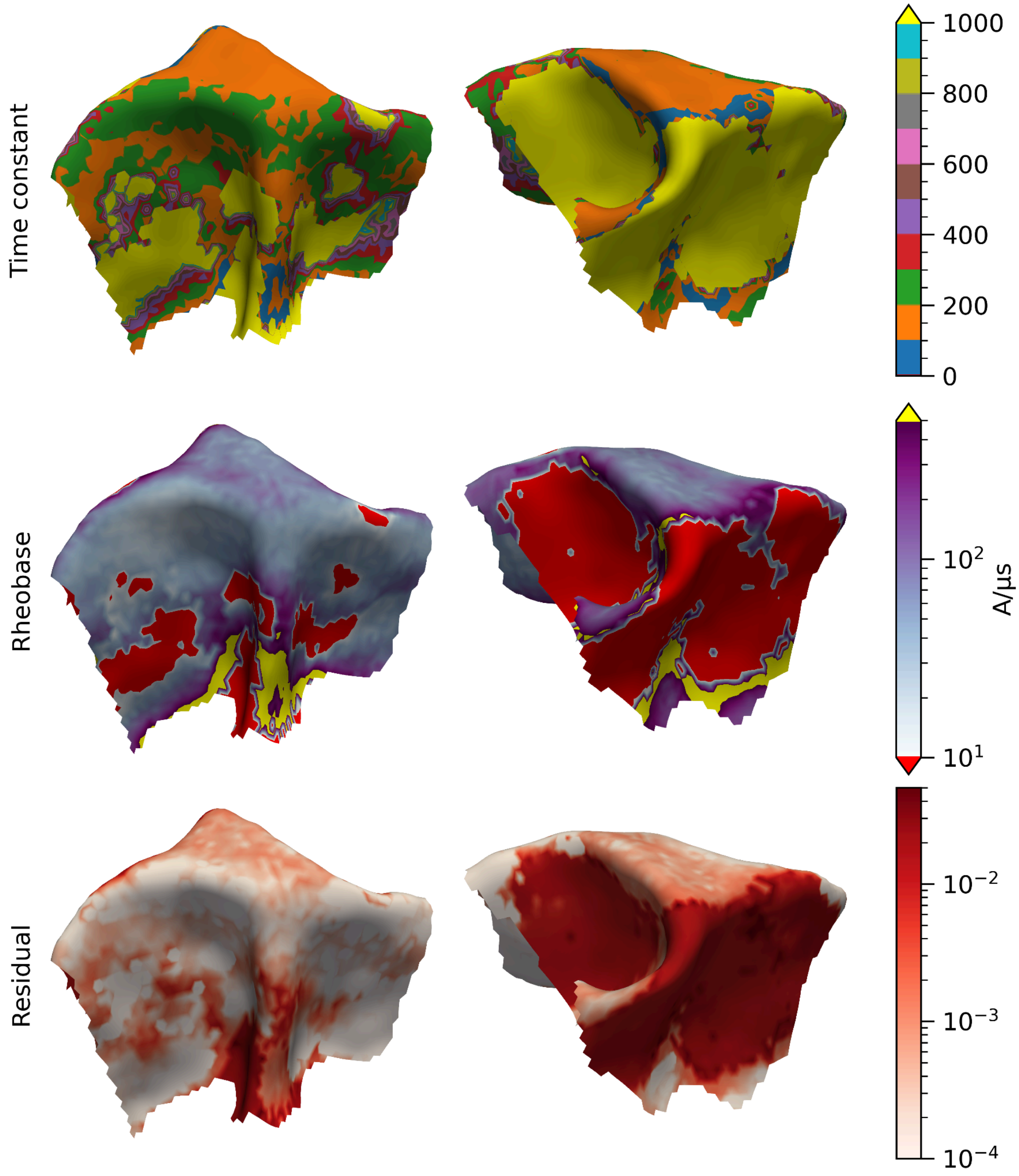


**Figure S5:** Simulation-based time constant estimation for smoothly bending axon projections with a diameter of 6 µm and cTMS pulses with PA direction. Time constants are binned in 100-µs intervals and capped at 1000 µs, and rheobase are capped at 500 A/µs.

#### Effects of the Anisotropic Conductivity Distribution

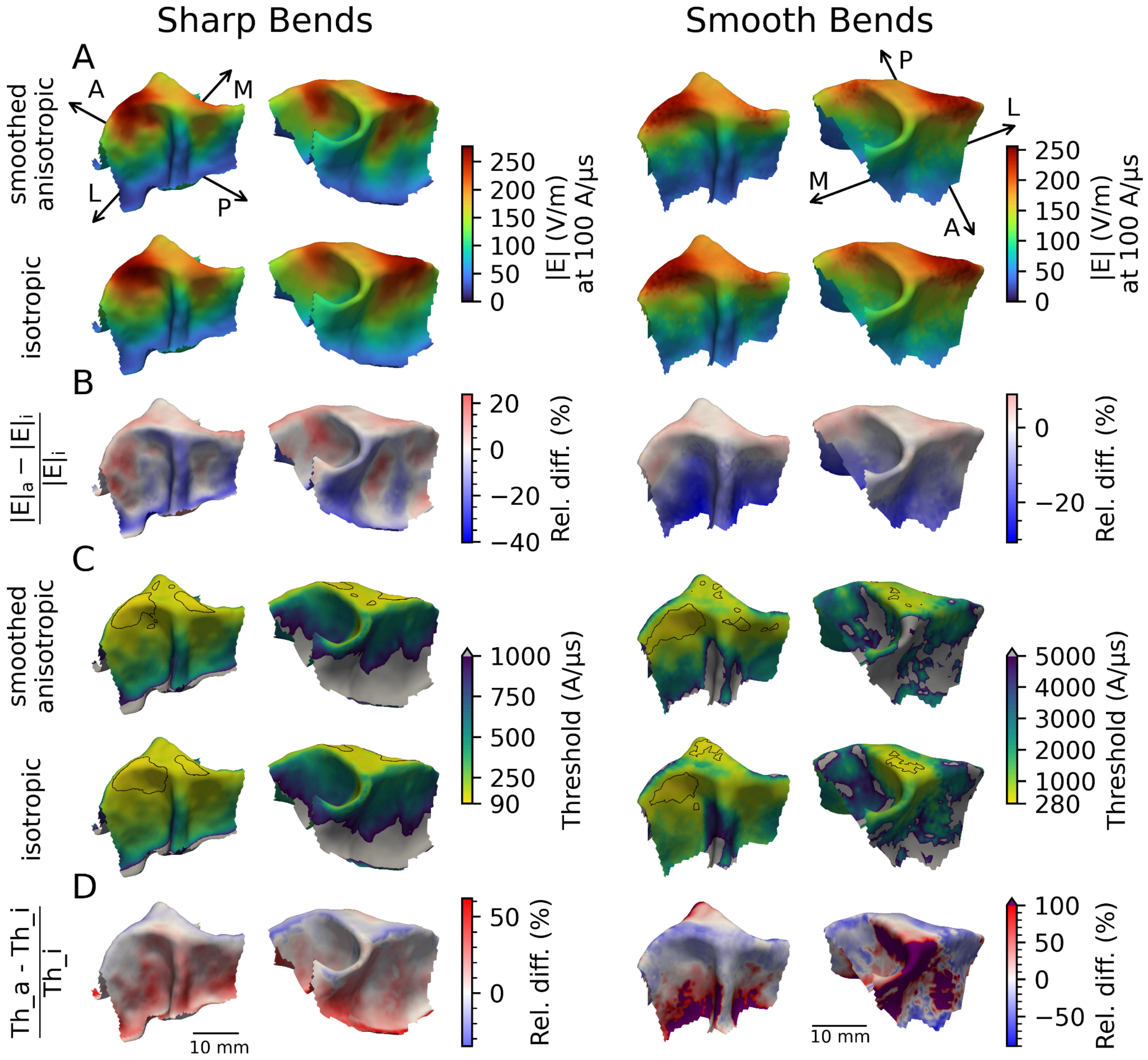


**Figure S6:** The effects of the isotropic and the smoothed anisotropic conductivity models on activation thresholds towards a monophasic PA stimulation pulse. The sharp bending axons were 0.5 mm below the WM surface with a bend radius of 100 μm, a diameter of 2 μm and an internode length ratio (*L/D*) of 100. For each surface position, 18 axon directions are simulated. The smooth bending axons started 0.5 mm below the pial surface and terminate in deep WM. They had a diameter of 2 μm and an internode length ratio (*L/D*) of 100. (A) The sampled E-field magnitude at the bend, 0.5 mm below the WM surface for sharp bends and inside the WM at the center of the bend for the smooth bends. (B) The relative difference in E-field magnitude between the isotropic and the smoothed anisotropic conductivity model. (C) The activation thresholds for the sharp and smooth bends with the isotropic and the smoothed anisotropic conductivity model. The black border surrounds the regions with a minimum threshold of

127 A/µs for sharp bends and 580 A/µs for smooth bends. (D) The relative difference in activation threshold between the isotropic and the smoothed anisotropic conductivity model.

#### Smooth Bending White Matter Axon Projections

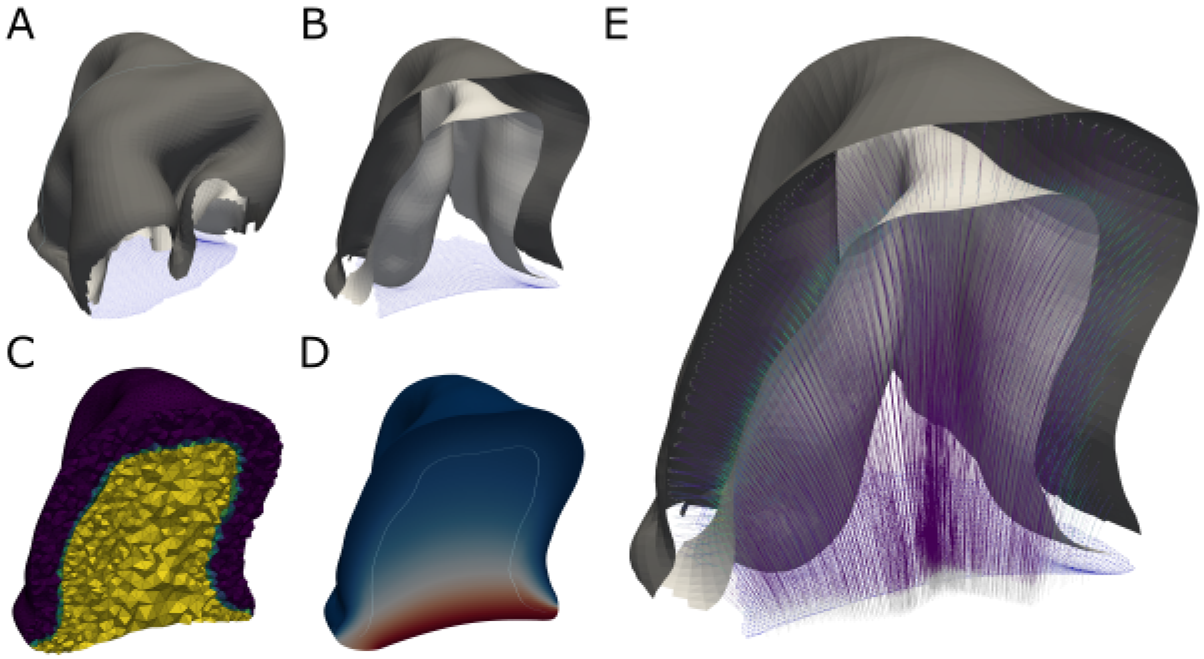


**Figure S7**. Overview of the deep WM axon generation. (A,B) The region of interest (ROI) is defined, the WM and GM surfaces are extracted and the deep WM (blue) is estimated. (C) Diffusivities are assigned to the different compartments. (D) The Laplace equation is solved, giving the potential on the domain. We set boundary conditions on the deep WM (0 V) and GM (1000 V) surfaces. (E) The E-field is calculated from the potential field by differentiation and streamlines are traced using the forward Euler method, starting just below the GM surface and terminating at the deep WM boundary. The color denotes the curvature at each position.

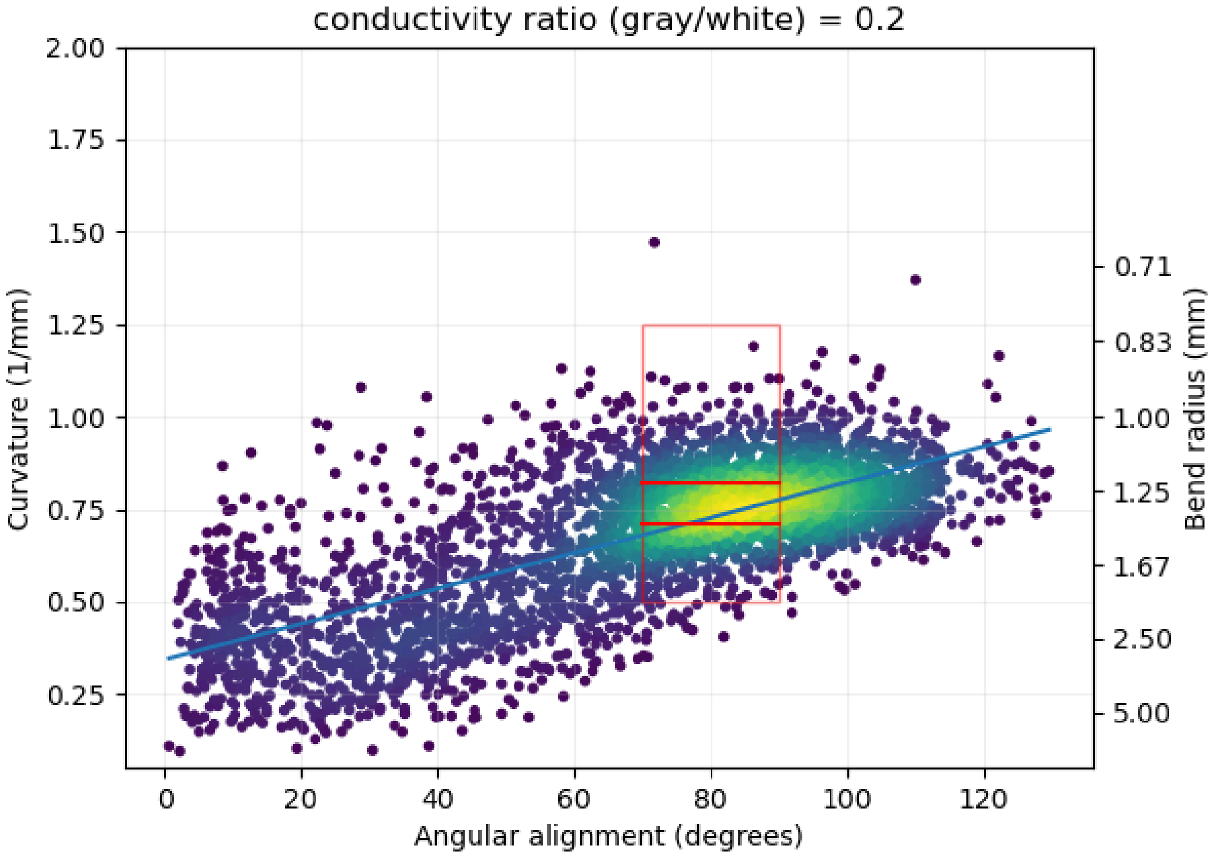


**Figure S8**. Relationship between "angular alignment" of fibers (i.e., the alignment of the surface normal at the seed point and the primary sulcal direction of the handknob area) and their maximum curvature and minimum bend radius (left and right, respectively) using a diffusivity ratio (GM/WM) of 0.2. The red box denotes the approximate range of expected bend radii (800–2000 μm) between 70°–90° alignment. The two red lines inside the box correspond to 1217 μm (from [55]) and 1410 μm (derived from [67]). Each dot, colored by a Gaussian kernel density estimate, corresponds to a fiber.

### Relationship Between Model Parameters and Activation Threshold

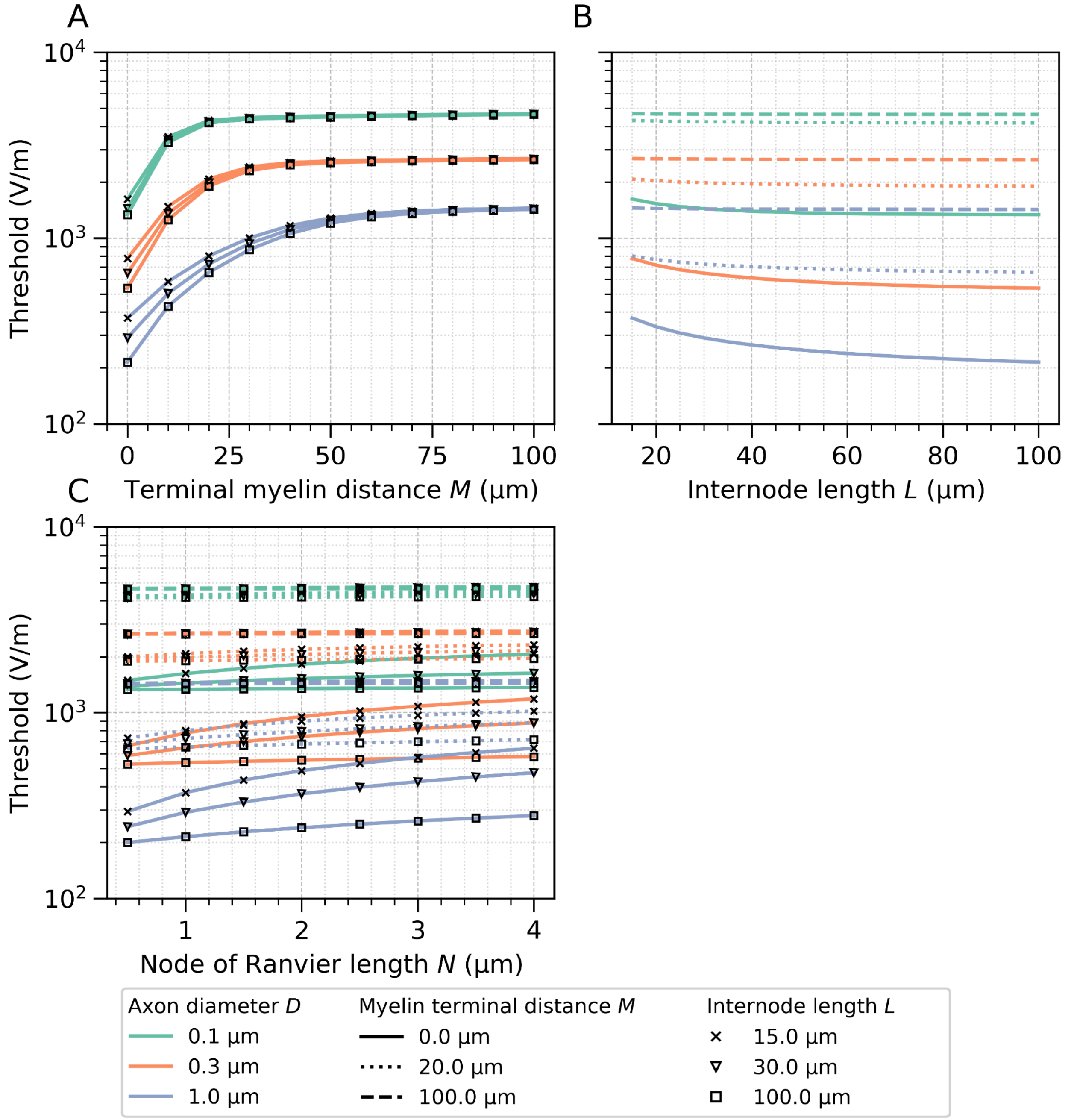


**Figure S9:** Relationship between terminal model parameters and the activation threshold for (A) the distance between the first internode and the terminal, (B) the internode length and (C) the node of Ranvier length. For panels A and B, the nodes of Ranvier had a length of 1 µm.

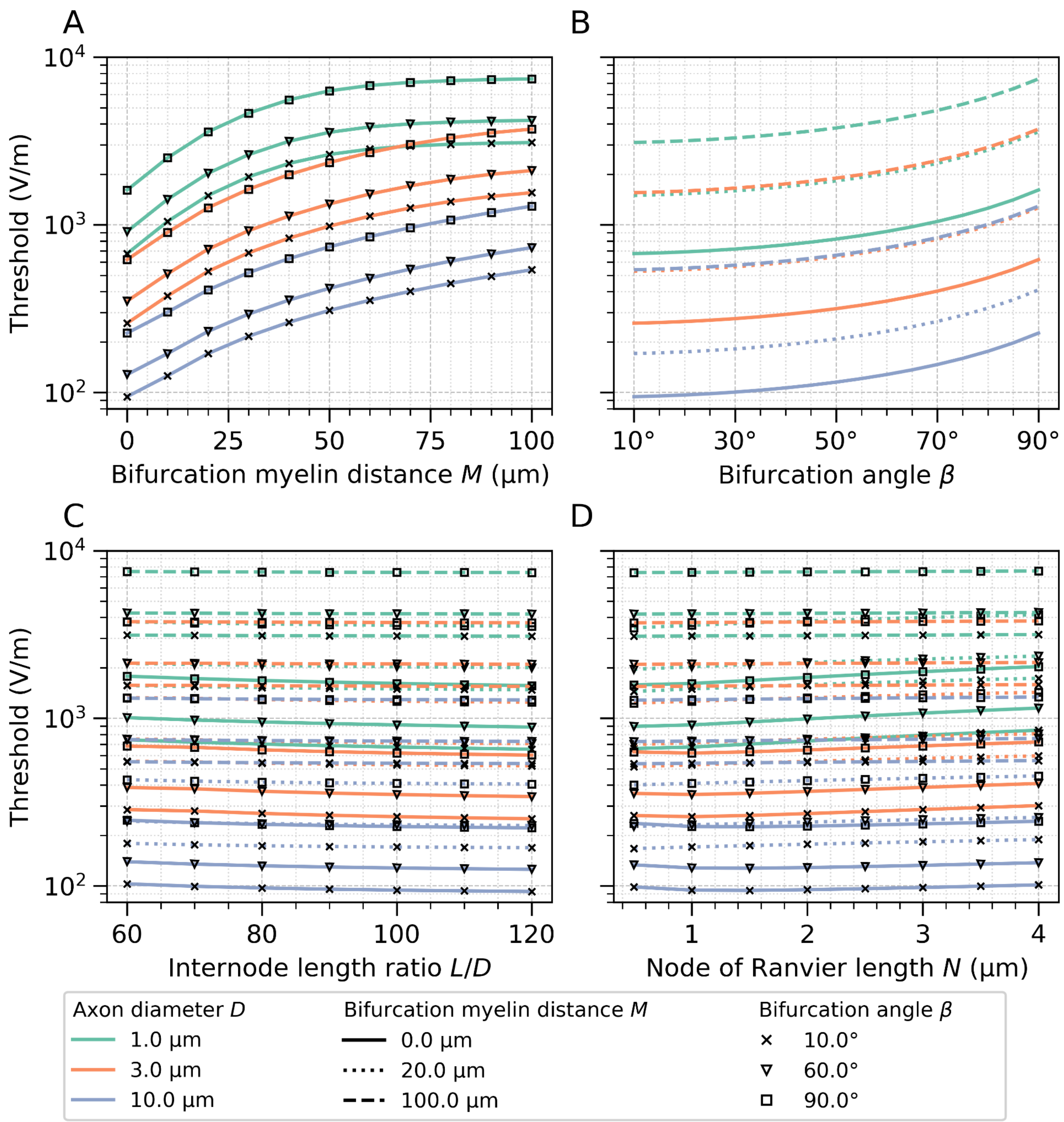


**Figure S10:** Relationship between bend model parameters and the activation threshold for (A) the bend radius, (B) the bend angle, (C) the internode length ratio ($L/D$) and (D) the node of Ranvier length. For panels A–C, the nodes of Ranvier had a length of 1 µm and for panels A, B, and D, the internode length ratio ($L/D$) was 100.

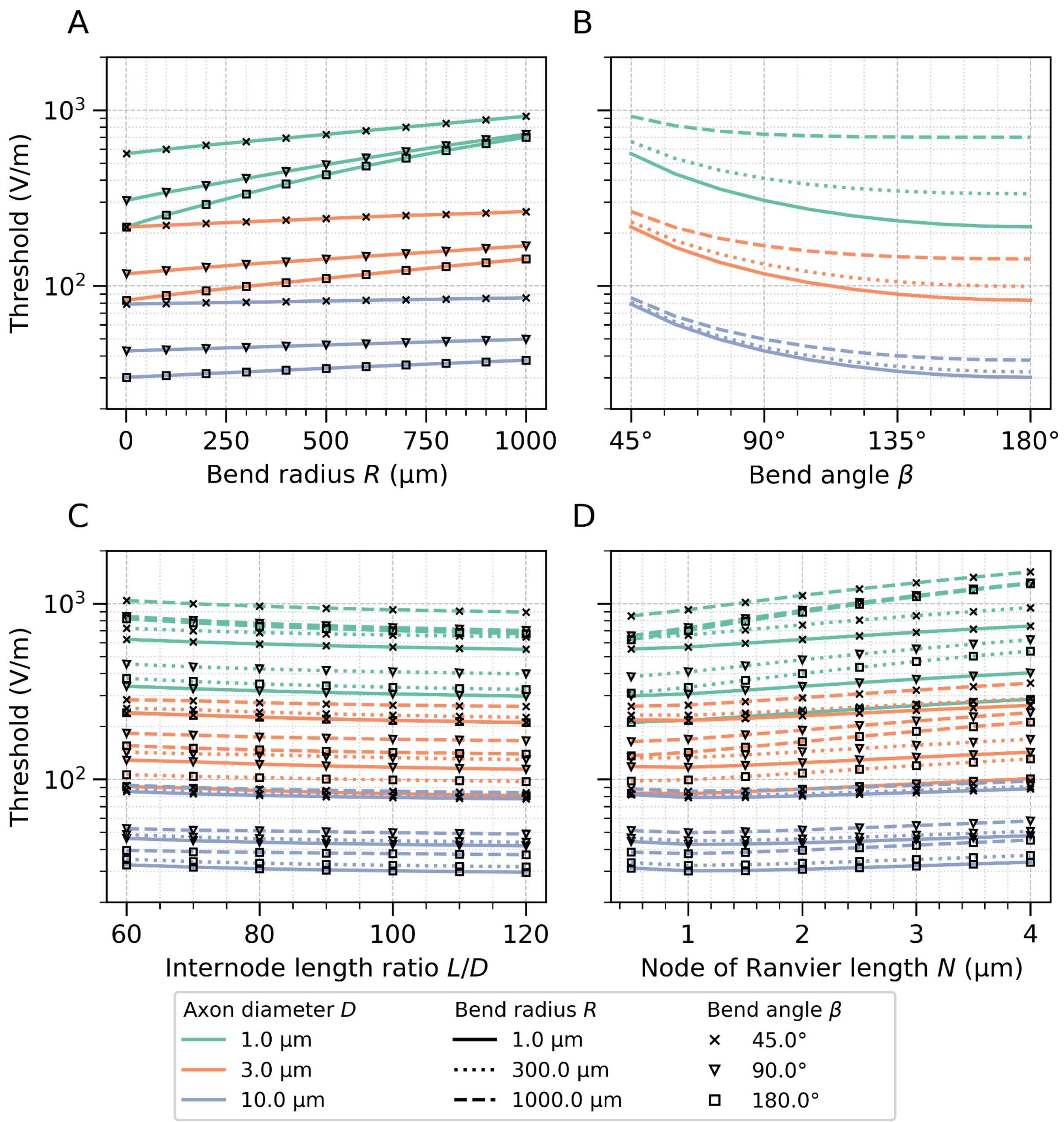


**Figure S11:** Relationship between bifurcation model parameters and the activation threshold for (A) the distance after the bifurcation between the first internodes and the bifurcation, (B) the bifurcation angle, (C) the internode length ratio (*L*/*D*) and (D) the node of Ranvier length. For panels A–C, the nodes of Ranvier had a length of 1 µm and for panels A, B, and D, the internode length ratio (*L*/*D*) was 100.

### Sensitivity of the Bend Model

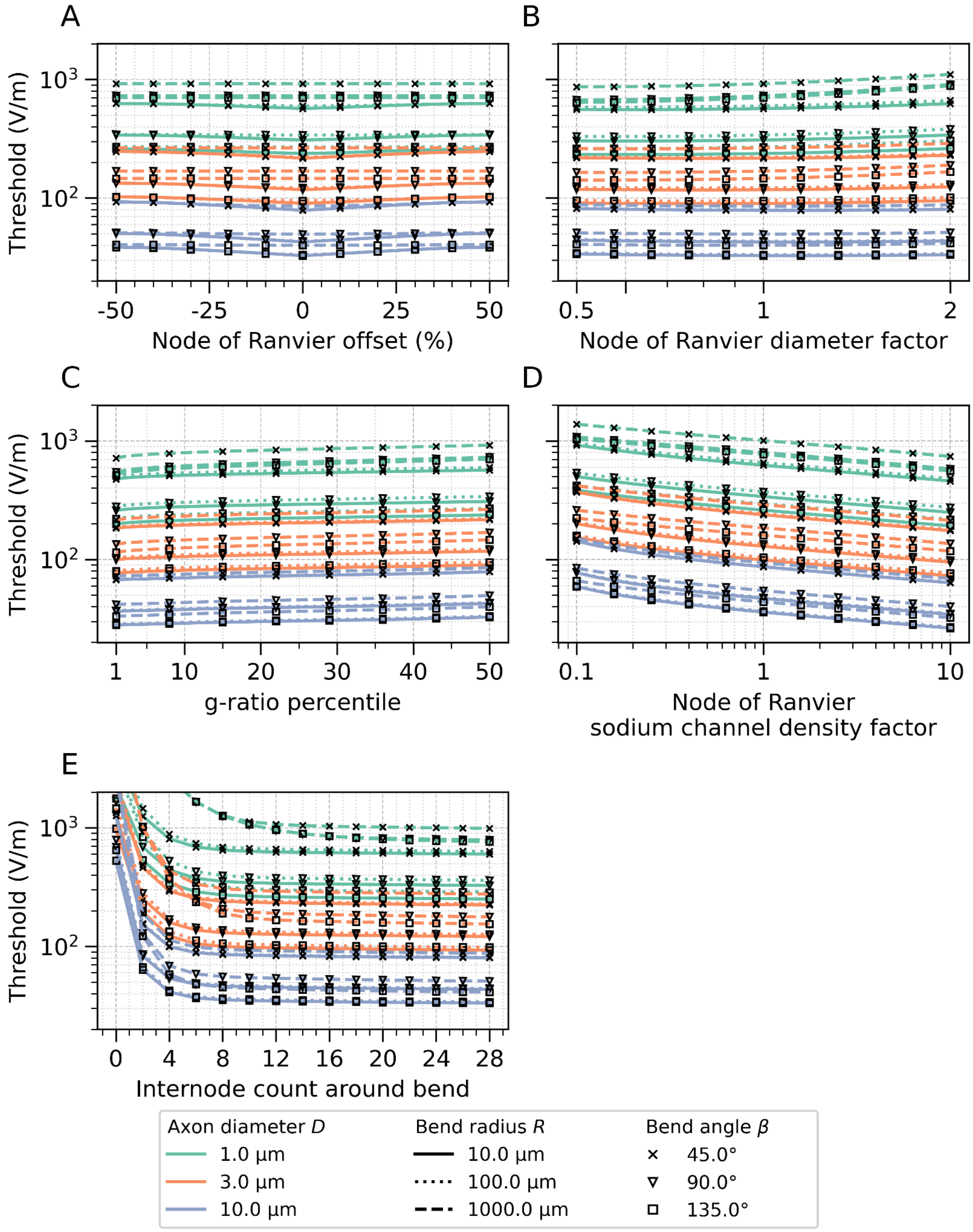

**Figure S12:** Relationship between bend model parameters and the activation threshold. (A) Offset of the node of Ranvier from the center of the bend, where 0% indicates a node at the bend center and 50% indicates the closest node positioned half an internode length from the center. (B) Variation in node of Ranvier diameters from half to twice the axon diameter. (C) Variation in the fitted g-ratio logarithmic function by changing the included measured g-ratios from the lowest 50% to the lowest 1% (see Appendix A). (D) Variation in maximal sodium conductance density (proportional to sodium channel density) from one-tenth to ten times the original value. (E) Variation in the number of internodes centered around the bend, where 0 corresponds to an unmyelinated axon, 2 corresponds to one internode before and one after the bend, and 28 corresponds to 14 before and 14 after the bend. For all panels A–E, nodes of Ranvier were 1 μm long, and the internode length ratio ($L/D$) was 100.

#### Simulation Results of Sharp Bends in Layer I

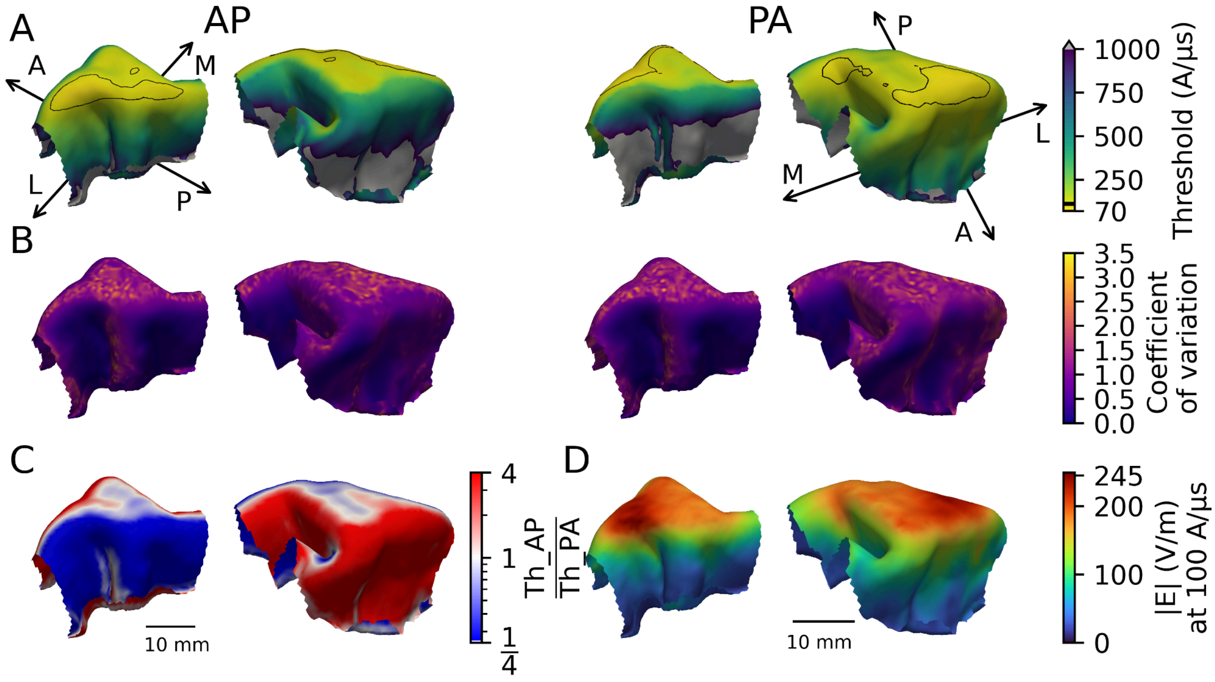


**Figure S13:** Excitability towards a monophasic pulse of bend axons centered in layer I with a bend radius of 100 µm, a diameter of 2 µm and an internode length factor (*L*/*D*) of 100. For each surface position, 18 axon directions are simulated. (A) The minimum activation thresholds (in A/µs) for each position for monophasic anterior–posterior (AP, left) and posterior–anterior (PA, right) stimulation. The black border surrounds the regions with a minimum threshold at or below the 5th percentile (AP: 115 A/µs, PA: 110 A/µs). Medial–lateral and anterior–posterior anatomical axes are annotated on the surfaces. (B) The coefficient of variation $\left(\frac{SD}{Mean}\right)$ of activation thresholds across the 18 axon orientations at each surface position. (C) Log-scaled ratio between AP and PA stimulation thresholds at each surface position. (D) E-field magnitude interpolated in the center of layer I for a stimulus amplitude of 1 A/µs.

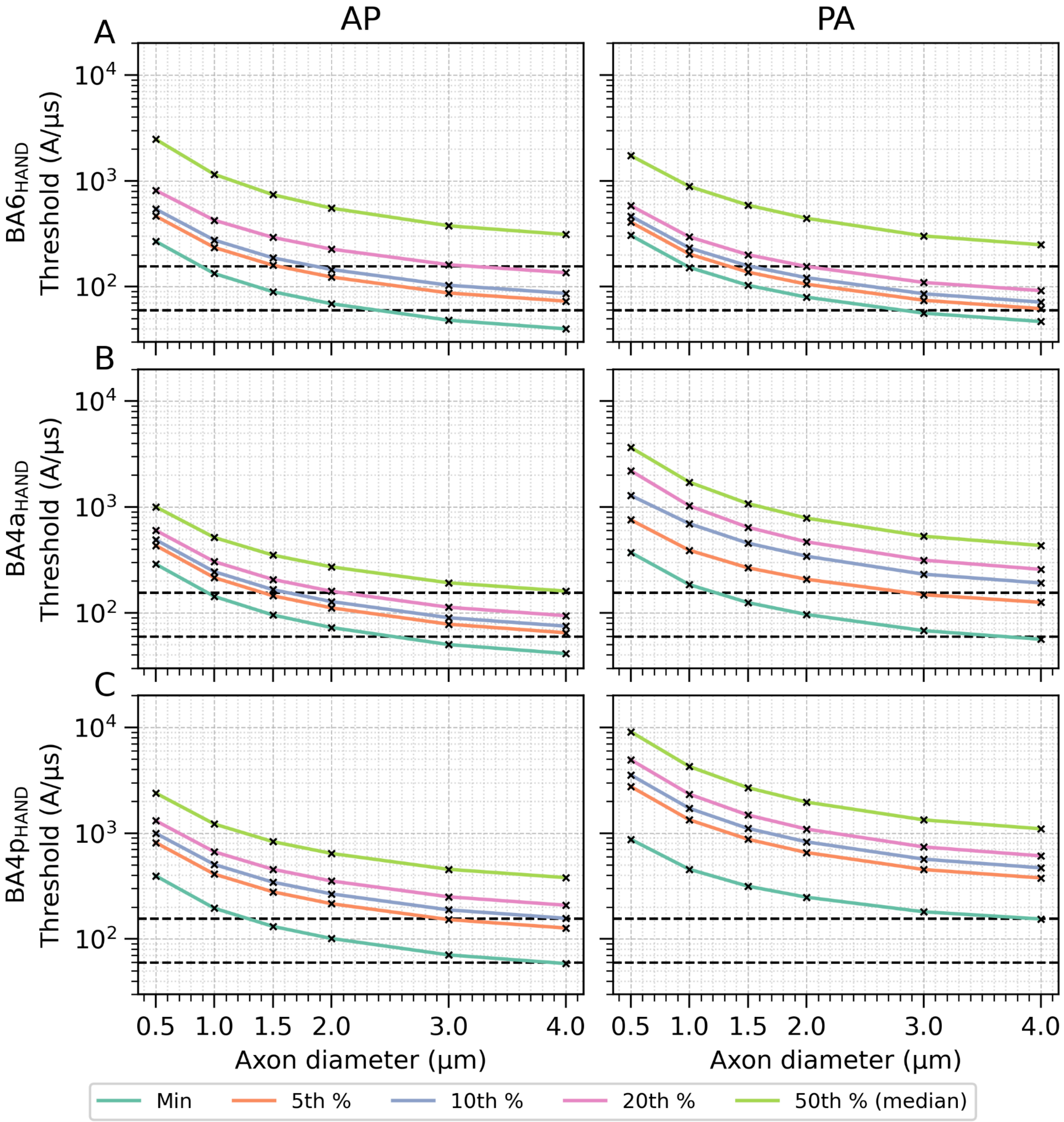


**Figure S14:** Excitability towards a monophasic pulse of bend axon populations located at the center of layer I, with a bend radius of 100 µm and an internode length factor (*L*/*D*) of 100. Simulations were run separately for six axon diameters (0.5, 1.0, 1.5, 2.0, 3.0 and 4.0 µm). For each condition, the minimum activation threshold and four percentiles (5th, 10th, 20th, and 50th) of the threshold distribution for 18 rotations per location are shown for three cortical regions: (A) $BA6_{HAND}$ with 555 locations, (B) $BA4a_{HAND}$ with 558 locations and (C) $BA4p_{HAND}$ with 1537 locations. The lower dashed line marks the experimentally estimated motor threshold of 59.7 A/µs, while the upper dashed line marks the maximum stimulator output at 155.3 A/µs.

#### Simulation Results of Sharp Bends in Layer IV

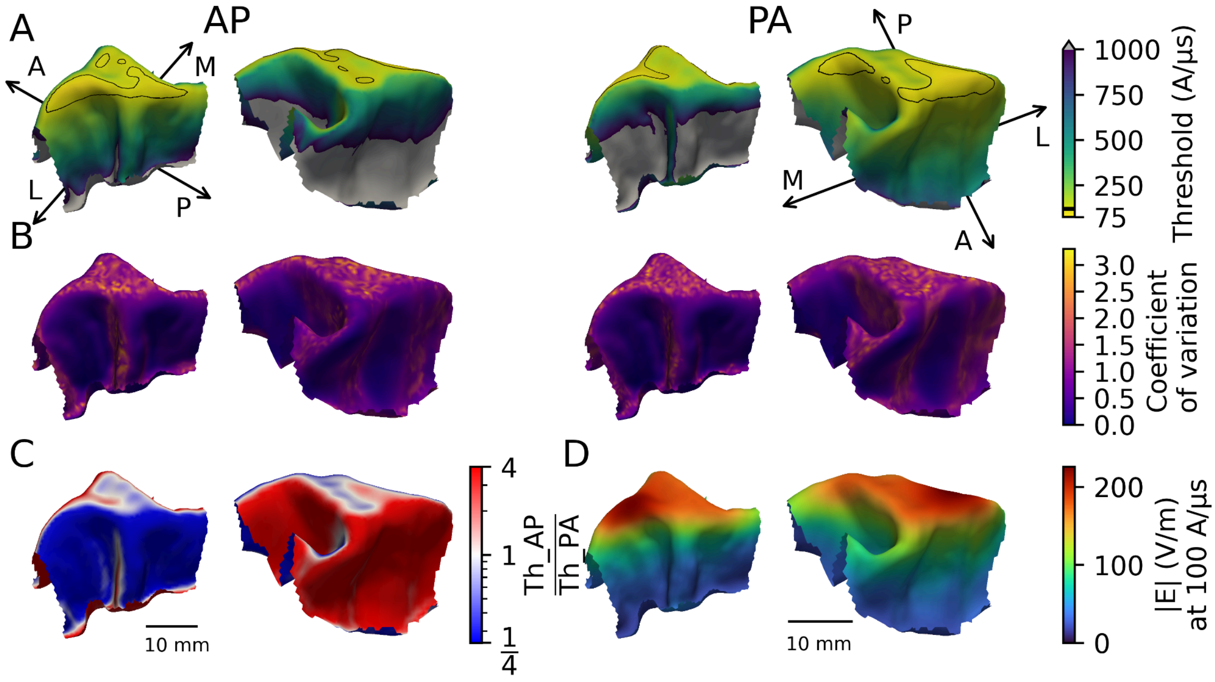


**Figure S15:** Excitability towards a monophasic pulse of bend axons centered in layer IV with a bend radius of 100 µm, a diameter of 2 µm and an internode length factor (*L*/*D*) of 100. For each surface position, 18 axon directions are simulated. (A) The minimum activation thresholds (in A/µs) for each position for monophasic anterior–posterior (AP, left) and posterior–anterior (PA, right) stimulation. The black border surrounds the regions with a minimum threshold at or below the 5th percentile (AP: 124 A/µs, PA: 119 A/µs). Medial–lateral and anterior–posterior anatomical axes are annotated on the surfaces. (B) The coefficient of variation $\left(\frac{SD}{Mean}\right)$ of activation thresholds across the 18 axon orientations at each surface position. (C) Log-scaled ratio between AP and PA stimulation thresholds at each surface position. (D) E-field magnitude interpolated in the center of layer IV for a stimulus amplitude of 1 A/µs.

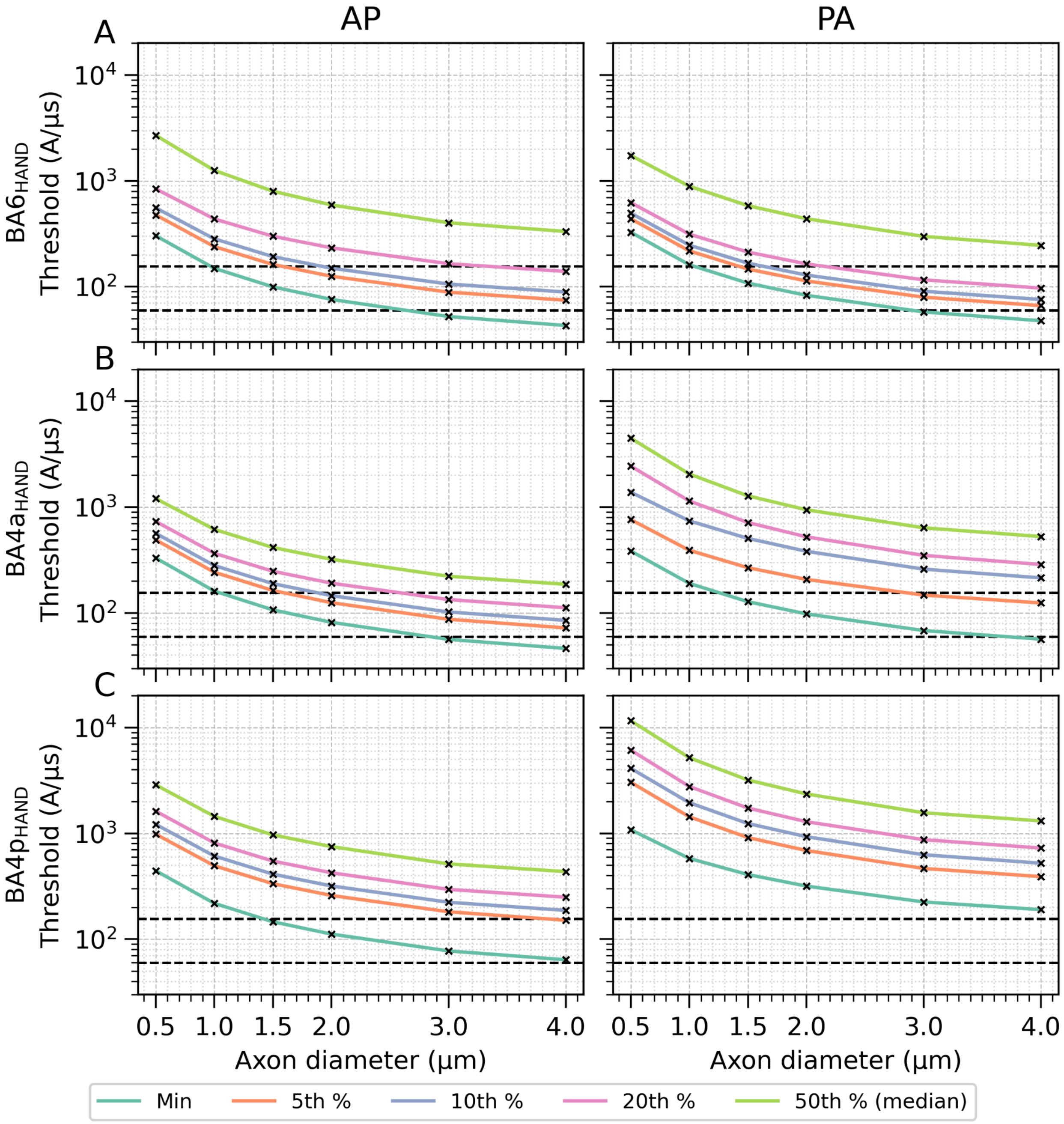


**Figure S16:** Excitability towards a monophasic pulse of bend axon populations located at the center of layer IV, with a bend radius of 100 µm and an internode length factor (*L*/*D*) of 100. Simulations were run separately for six axon diameters (0.5, 1.0, 1.5, 2.0, 3.0, and 4.0 µm). For each condition, the minimum activation threshold and four percentiles (5th, 10th, 20th, and 50th) of the threshold distribution for 18 rotations per location are shown for three cortical regions: (A) $BA6_{HAND}$ with 555 locations, (B) $BA4a_{HAND}$ with 558 locations and (C) $BA4p_{HAND}$ with 1537 locations. The lower dashed line marks the experimentally estimated motor threshold of 59.7 A/µs, while the upper dashed line marks the maximum stimulator output at 155.3 A/µs.

### Biphasic Stimulation of Bends

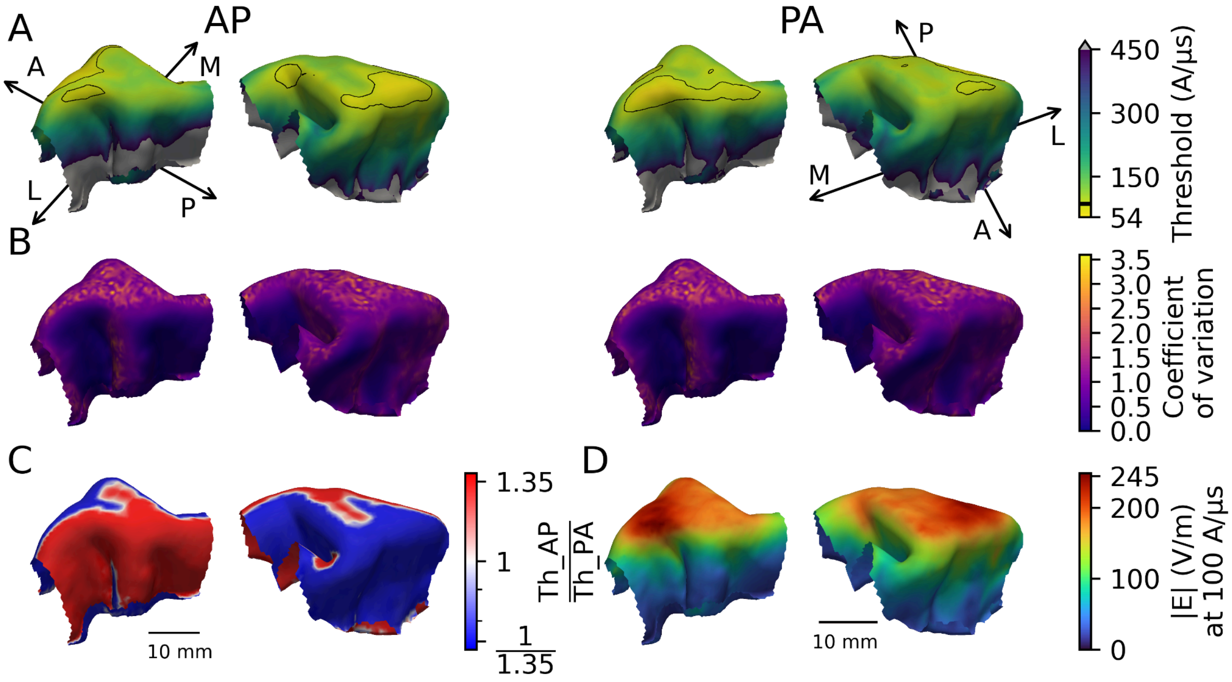


**Figure S17:** Excitability towards a biphasic pulse of bend axons centered in layer I with a bend radius of 100 µm, a diameter of 2 µm and an internode length factor (*L*/*D*) of 100. For each surface position, 18 axon directions are simulated. (A) The minimum activation thresholds (in A/µs) for each position for biphasic AP (left) and PA (right) stimulation. The black border surrounds the regions with a minimum threshold at or below the 5th percentile (AP: 84 A/µs, PA: 88 A/µs). Medial–lateral and anterior–posterior anatomical axes are annotated on the surfaces. (B) The coefficient of variation $\left(\frac{SD}{Mean}\right)$ of activation thresholds across the 18 axon orientations at each surface position. (C) Log-scaled ratio between AP and PA stimulation thresholds at each surface position. (D) E-field magnitude interpolated in the center of layer I for a stimulus amplitude of 100 A/µs.

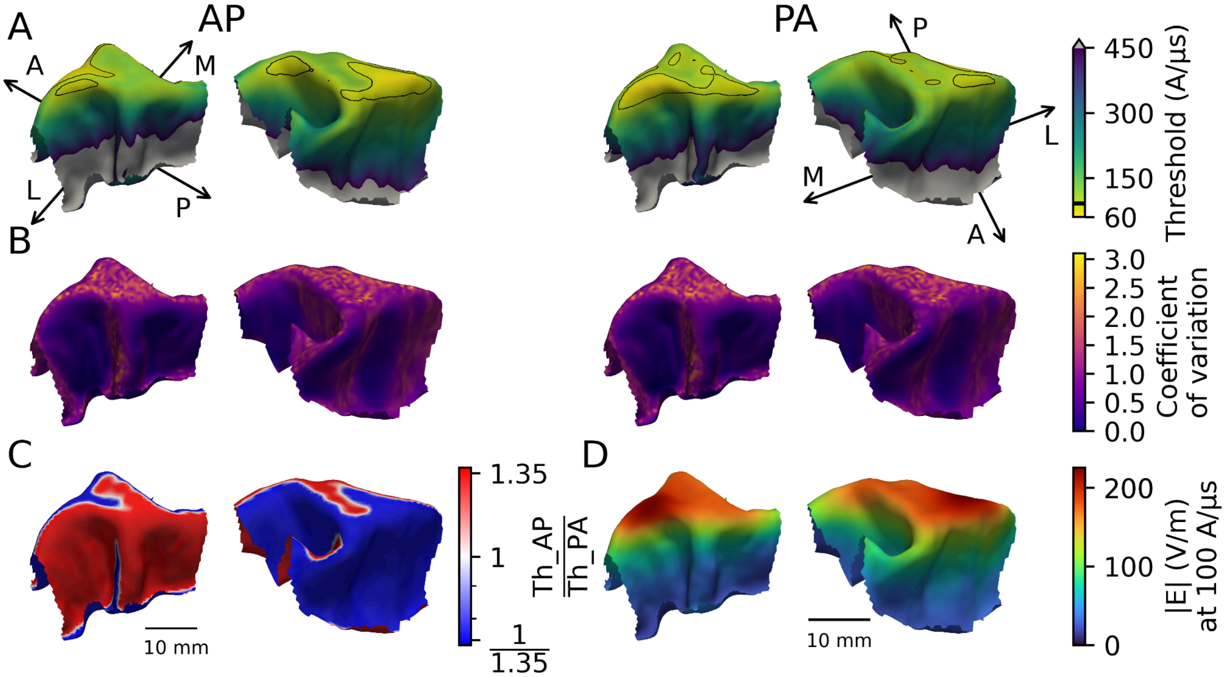


**Figure S18:** Excitability towards a biphasic pulse of bend axons centered in layer IV with a bend radius of 100 µm, a diameter of 2 µm and an internode length factor (*L*/*D*) of 100. For each surface position, 18 axon directions are simulated. (A) The minimum activation thresholds (in A/µs) for each position for biphasic AP (left) and PA (right) stimulation. The black border surrounds the regions with a minimum threshold at or below the 5th percentile (AP: 91 A/µs, PA: 95 A/µs). Medial–lateral and anterior–posterior anatomical axes are annotated on the surfaces. (B) The coefficient of variation $\left(\frac{SD}{Mean}\right)$ of activation thresholds across the 18 axon orientations at each surface position. (C) Log-scaled ratio between AP and PA stimulation thresholds at each surface position. (D) E-field magnitude interpolated in the center of layer IV for a stimulus amplitude of 100 A/µs.

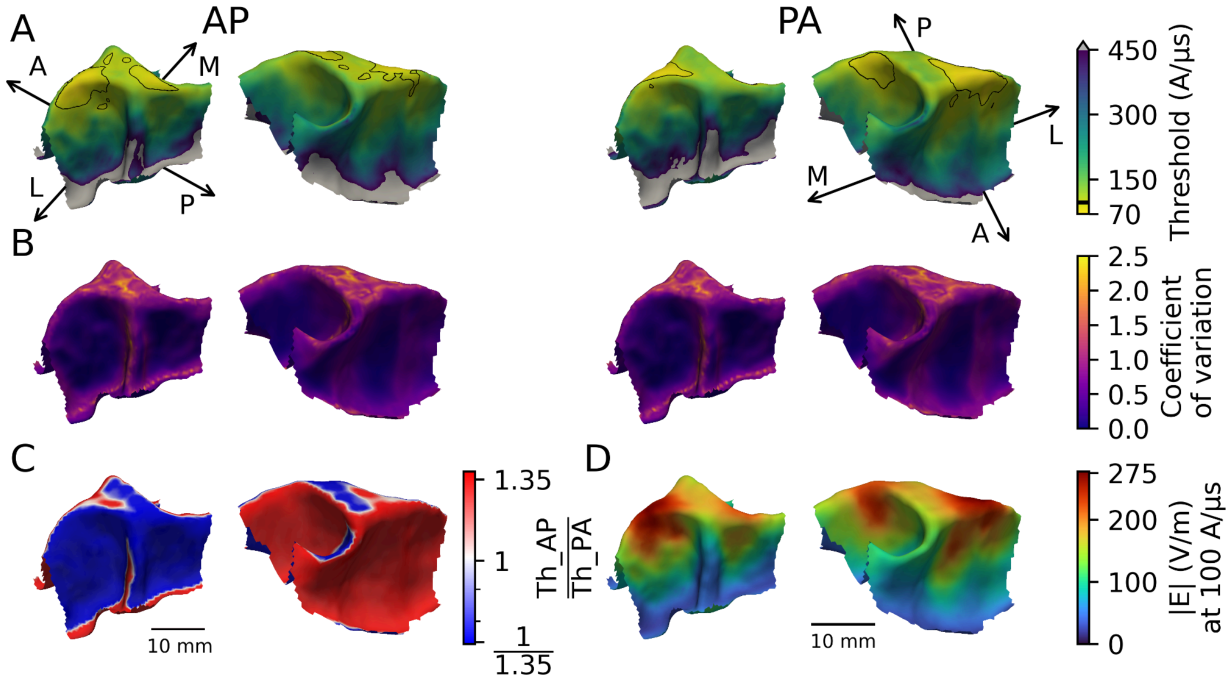


**Figure S19:** Excitability towards a biphasic pulse of bend axons 0.5 mm below the WM surface with a bend radius of 100 μm, a diameter of 2 μm and an internode length factor (*L*/*D*) of 100. For each surface position, 18 axon directions are simulated. (A) The minimum activation thresholds (in A/μs) for each position for biphasic AP (left) and PA (right) stimulation. The black border surrounds the regions with a minimum threshold at or below the 5th percentile (AP: 100 A/μs, PA: 94 A/μs). Medial–lateral and anterior–posterior anatomical axes are annotated on the surfaces. (B) The coefficient of variation $\left(\frac{SD}{Mean}\right)$ of activation thresholds across the 18 axon orientations at each surface position. (C) Log-scaled ratio between AP and PA stimulation thresholds at each surface position. (D) E-field magnitude interpolated at 0.5 mm below the WM surface for a stimulus amplitude of 100 A/μs.

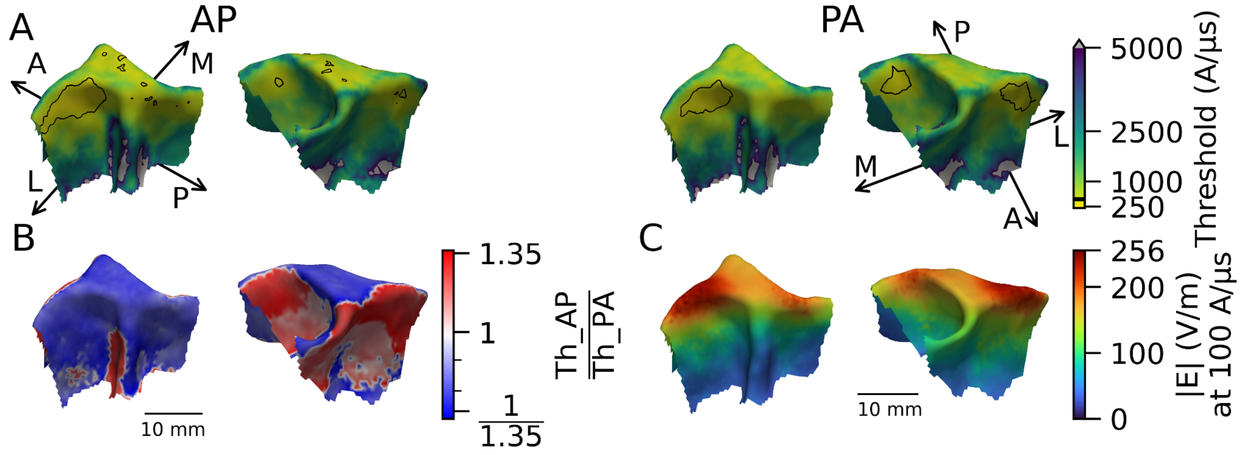


**Figure S20:** Excitability towards a biphasic pulse of smoothly bending axon projections into deep WM. The axons start 0.5 mm below the pial surface and terminate in deep WM. They have a diameter of 2 µm and an internode length factor (*L*/*D*) of 100. (A) The minimum activation thresholds (in A/µs) for each position for E-fields oriented in the AP (left) and PA (right) directions. The black border surrounds the regions with a minimum threshold at or below the 5th percentile (AP: 456 A/µs, PA: 472 A/µs). Medial–lateral and anterior–posterior anatomical axes are annotated on the surfaces. (B) Log-scaled ratio between AP and PA stimulation thresholds at each surface position. (D) E-field magnitude interpolated at the point of minimum bend radius of each simulated fiber.

### Supplementary Material I

**Table S-I1**: A non-exhaustive overview of the methods used in the studies presented in this section.

| Source | Post Mortem Interval | Fixative | Processing | Embedding | Sectioning | Staining | Mounting | Imaging Method |
|---|---|---|---|---|---|---|---|---|
| Benavides-Piccione et al. [33] | 1 - 4h | Immersion: 4% paraformaldehyde in phosphate buffer, 24h in 4% paraformaldehyde solution | - | - | Vibratome (300 µm) | Immunofluorescence: DAPI, Lucifer yellow, anti-LY antibody, Alexa Fluor 488 | ProLong GoldAntifade Reagent or Glycerol 50% in PB | Confocal Laser Scanning Microscopy |
| Szocsics et al. [35] | 2h53´ - 5h5´ | Immersion: Zamboni fixative, overnight in 4% PFA + 0.2% picric acid in phosphate buffer | Freezing: Sucrose cryoprotection, freeze–thaw treatment | - | Vibratome (60 µm) | Immunofluorescence: NeuN | Vectashield | Wide-Field Fluorescence Microscopy |
| Tomasi et al. [19] | Immediate | Perfusion: 4% PFA in phosphate buffer Immersion: overnight in the same | Freezing: Sucrose cryoprotection in PBS | - | Microtome (34 µm) | Injected: BDA Stained: Cresyl violet or Gallyas method for myelin | Not Reported | Brightfield Light Microscopy |

| | | | | | | | | |
|---|---|---|---|---|---|---|---|---|
| | | solution | | | | | | |
| Firmin et al. [13] | Immediate | Perfusion: 1% PFA and 1% glutaraldehyde in PBS Immersion: 24-48h in the same solution | Dehydration | Resin | Vibratome (200 μm), Not reported (1 μm), Not reported (100 nm) | 1% osmium tetroxide, 1% toluidene blue, 1% uranyl acetate and 1% lead citrate | Copper Grids | Electron Microscopy |
| Ruthig et al. [12] | 20h - 40h | Immersion: At least 4 weeks in 3% formaldehyde and 1% glutaraldehyde in PBS buffer | Stored in 0.2M cacodylate buffer, Dehydration | Durcupan™ araldite resin | Vibratome (70 μm), ultramicrotome (50 nm) | 1% osmium tetroxide in cacodylate buffer, 1% uranyl acetate in 70% acetone | slot grids coated with a 10–20 nm Formvar film | Transmission Electron Microscopy |
| Andersson et al. [68] | Immediate (Stored) | Perfusion: 4% PFA in PBS buffer immersion: 2.5% glutaraldehyde | Dehydration | EPON resin | Not reported (2-4 mm), biopsy punch (diameter 1mm) | 0.5% osmium tetroxide | - | Synchrotron XNH Imaging |
| Liewald et al. [23] | ≤ 48 h (Stored) | Immersion: 4% formaldehyde | Dehydration | Epon / Araldite | Not reported | 2% osmium tetroxide, uranyl acetate and lead citrate | mesh grids | Electron Microscopy |
| Skoven et | Immediate | Perfusion: | 2 weeks in a | Epon | Not reported | uranyl acetate | Not | Transmission Electron |

| | | | | | | | | |
|---|---|---|---|---|---|---|---|---|
| al. [14] | | 4% formaldehyde <u>Immersion</u>: 3 weeks in the same solution | 2.5% solution of glutaraldehyde in cacodylate buffer, <u>Dehydration</u> | | (3-4 mm), biopsy punch (diameter 1mm), ultramicrotome (~40-70 nm) | and lead citrate | Reported | Microscopy |
| Skoven et al. [14] | Immediate | <u>Perfusion</u>: 4% formaldehyde | 2h in KPBS, <u>Freezing</u>: high-pressure freezing, freeze substitution using 1% $OsO_4$ in acetone | Epon | Not reported (3-4 mm), vibratome (200 µm), ultramicrotome (50-70 nm) | uranyl acetate and lead citrate | Not Reported | Transmission Electron Microscopy |
| Rivara et al. [34] | ≤12h | <u>Immersion</u>: 10% buffered formalin for up to 9 days | <u>Freezing</u>: Sucrose (up to 25%) cryoprotection | - | Not reported (50 µm) | cresyl violet | gelatin-coated slides and coverslipped with Entellan | Brightfield Light Microscopy |
| Fish et al. [69] | Immediate | <u>Perfusion</u>: 1% PFA in phosphate buffer, 4% PFA in phosphate buffer | <u>Freezing</u>: Sucrose cryoprotection, <u>Permeabilization</u>: 0.3% Triton X-100 in PBS for 30 min | - | Not reported (5 mm), Not reported (40 µm) | <u>Immunofluorescence</u>: PV, vGAT, γ2, AnkG, NeuN | Not Reported | Spinning Disk Confocal Microscopy |
| Senol et al | 5h - 33h | <u>Immersion</u>: | dried, | - | Not reported | <u>Immunofluores</u> | Dapi-Fluoro | Fluorescence |

| | | | | | | | | |
|---|---|---|---|---|---|---|---|---|
| [38] | | −50°C isopentane, stored at −85°C, 2% PFA for 5 min | Blocking: 10% NGS and 0.4% Triton | | (1cm), Not reported (20 μm) | cence: AnkG, SMI32, AnkG, Calbindin | mount-G | Microscopy |
| Li et al. [37] | Not reported (Stored) | Chemical fixation | Freezing: Snap freezing | - | Not reported | Immunofluorescence: See source | Not reported | Not reported |
| Sloper and Powell [39] | Immediate | Perfusion: 4% PFA and 1% glutaraldehyde Immersion: Same solution, 2 % osmic acid in phosphate buffer | Dehydration | Epon-Araldite | Not reported (1 mm), Not reported (1 μm) | uranylacetate, methylene blue and azure II, alkaline lead citrate and uranyl acetate | Formvar-coated copper grid | Electron Microscopy |
| Höfflin et al. [40] | organotypic tissue culture | Immersion: 4% PFA for 2h | Blocking: 3% bovine serum albumin, 3% normal goat serum, and 0.5% Triton X-100 in TBS | - | McIlwain tissue chopper (350 μm) | Immunofluorescence: mCherry and ßIV-spectrin | glass coverslips using Roti®–Mount | Fluorescence Microscopy |
| Pascual-García et al. [45] | Not reported | Immersion: transported in | Blocking: PBS containing | - | Not reported (300 μm), Not reported (40 | Filling: with 5 mg/mL biocytin | cover-slipped and sealed | Fluorescence Microscopy |

| | | | | | | | | |
|---|---|---|---|---|---|---|---|---|
| | | carboxygenated, 4% PFA overnight | 0.4% Triton X-100 and 5% BSA | | µm) | Immunofluorescence: Alexa488 and Cy3-secondary | | |
| Stedehouder et al. [41] | <10 min | Immersion: transported in carboxygenated, 4% PFA overnight | Blocking: 0.5% Triton X-100% and 5% bovine serum albumin | - | Not reported (300 µm), freezing microtome (40 µm) | Filling: biocytin Immunofluorescence: Streptavidin–Cy3, Mouse anti-MBP, Alexa-conjugated secondary antibodies | slides, cover slipped with Vectashield H1000 fluorescent mounting medium | Fluorescence Microscopy |
| Peters et al. [20] | Immediate | Perfusion: 1% PFA and 1.25% glutaraldehyde in either a phosphate buffer or cacodylate buffer Immersion: several days in 2% paraformaldehyde and 2.5% glutaraldehyde | Dehydration | Araldite | Not reported (2 mm), Not reported (“Thin sections”) | osmic acid, uranyl acetate and lead citrate | Not reported | Electron Microscopy |
| Zikopoulos | 14h - 20h | Immersion: | microwave | Araldite at | Vibratome | 1% uranyl | single-slot | Transmission Electron |

| | | | | | | | | |
|---|---|---|---|---|---|---|---|---|
| and Barbas [24] | (Stored) | 2% PFA and 2.5% glutaraldehyde, in phosphate buffer for 2d, stored again at −20°C in antifreeze solution, 1% osmium tetroxide with 1.5% potassium ferrocyanide in PB | with 6% glutaraldehyde, Dehydration Clearing: propylene oxide | 60°C | (50 µm), ultramicrotome (50 nm) | | grids | Microscopy |
| Innocenti et al. [21] | Immediate | Perfusion: sotonic saline followed by 4% PFA in phosphate buffered saline Immersion: overnight in the same solution | Freezing: Sucrose (30%) cryoprotection | - | microtome (34 µm) | Filling: biotinylated dextran amine Staining: Cresyl Violet or the Gallyas method | Not reported | Light Microscopy |
| Graf von Keyserlingk and Schramm [22] | 5h - 10h | Post Mortem Perfusion: 1% PFA and 1% glutaraldehyde | Dehydration | Araldite | razorblade (“thick slices”), Not reported (“further divided”), | 0.5% uranyl acetate, uranyl acetate and lead citrate | square mesh grids | Electron Microscopy |

| | | | | | | | | |
|---|---|---|---|---|---|---|---|---|
| | | Immersion: 1 % osmium tetroxide in sodium cacodylate | | | vibratom (20 µm) | | | |
| Yamashita et al. [42] | Immediate | Perfusion: 0.02% heparin sodium in NaCl or Ringer's solution, 2% PFA and 0.5% glutaraldehyde in phosphate buffer Immersion: overnight in 4% PFA in PB | Freezing: Sucrose (30%) cryoprotection Dehydration Clearing: xylene | - | freezing microtome (50 µm) | Filling: BDA, BDA was visualised with avidin-biotin-complex peroxidase method using the ABC elite kit | gelatin-coated slides covered with coverslips | Light Microscopy |
| DeFelipe et al. [43] | Immediate | Perfusion: 1% PFA and 2.5% in glutaraldehyde phosphate buffer | Freezing: Sucrose cryoprotection | - | frozen (60 µm) | Filling: HRP, cobalt-enhanced diaminobenzidine method | | Light Microscopy |
| Rockland and Virga [49] | Immediate | Perfusion: 0.9% NaCl and 0.5% NaNO3 in | Freezing: Sucrose (30%) cryoprotecti | - | freezing microtome (40-50 µm) | Filling: PHA-L, avidin-biotin immunoperoxi | coverslipped from xylene | Light Microscopy |

| | | | | | | | | |
|---|---|---|---|---|---|---|---|---|
| | | distilled $H_2O$, 4% PFA in PB, 2.5-10% sucrose in PB | on, Dehydration | | | dase method, incubating in 0.05% DAB and 0.1% nickel ammonium sulfate with 0.009% $H_2O_2$, 0.01% osmium | | |
| Mortazavi et al. [51] | Immediate | Perfusion: Krebs–Heinsleit buffer, buffered 4% PFA with up to 0.01% gadolinium Immersion: 4% PFA with up to 0.01% gadolinium overnight | Freezing: glycerol cryoprotection, flash frozen in −75°C 2-methyl-butane, Dehydration Clearing | | sliding microtome (30 μm) | Filling: BDA, incubated with Vector ABC Elite solution and nickel (0.05%) 3,3′-diaminobenzidene tetrachloride solution and 0.03% $H_2O_2$, counterstained with neutral red | gelatin-coated slides, cover-slipped with Permount | Confocal Laser Scanning Microscopy |
| Hess and Young [70] | Immediate | Immersion: ammonium molybdate | Teasing: glycerine | - | Not reported | perfusion methylene blue | Not reported | Light Microscopy |
| Ford et al. [27] | Immediate | Perfusion: Ringer's solution containing | Blocking: 1% bovine serum albumin, 2% | - | Vibratome (120 μm) | Filling: tetramethylrhodamine dextran, | coverslipped with Vectashield mounting | Confocal Laser Scanning Microscopy |

| | | | | | | | | |
|---|---|---|---|---|---|---|---|---|
| | | heparin, Immersion: 4% PFA overnight | Triton X-100 and 0.1% saponin in PBS | | | Immunofluorescence: ankyrin G, $K_v$1.2, NFH, Pan $Na_v$, MAP2 | medium | |
| Ibrahim et al. [26] | Immediate | Perfusion: 2.5% glutaraldehyde in PBS Immersion: Overnight in the same solution | Dehydration Clearing: xylene | - | Not reported | 2% $OsO_4$ in PBS, blackened in 70% alcohol | whole-mounted in DPX | Light Microscopy |
| Butt et al. [25] | Immediate | Perfusion: 4% PFA in PBS Immersion: 24h in the same solution, 15% triton in PBS for 24h | Blocking: 0.3% $H_2O_2$, 1.5% goat serum | - | Not reported | Immunofluorescence: Rip, Rabbit anti-mouse IgG, peroxidase antiperoxidase (PAP) technique | whole-mounted, dried onto untreated glass slides, mounted in citifluor | Light Microscopy |
| Stedehouder et al. [71] | <10 min | Immersion transported in oxygenated ice-cold cutting solution, 4% PFA overnight | Freezing: Sucrose (30%) cryoprotection, Blocking: 0.5% Triton X-100 and 5% bovine serum | - | freezing microtome (40 µm) | Immunofluorescence: mouse anti-PV, goat anti-MBP, Alexa-conjugated secondary antibodies, cyanine dyes, streptavidin-A4 | cover slipped with Vectashield H1000 fluorescent mounting medium | Confocal Microscopy |

| | | | | | | | | |
|---|---|---|---|---|---|---|---|---|
| | | | albumin | | | 88<br>Nuclear staining using DAPI | | |
| Micheva et al. [44] | Immediate | Perfusion:<br>2% glutaraldehyde, 2% formaldehyde in PB<br>Immersion:<br>Overnight in the same solution | Freezing:<br>glycerol cryoprotection, quick-frozen in a dry ice/ethanol bath, Freeze-substitution | infiltrated with Lowicryl HM-20 | Vibratome (200 µm), ultramicrotome (70 nm) | Immunofluorescence:<br>See source | gelatin-coated coverslips | Epi-Fluorescence Microscopy |
| Micheva et al. [44] | Immediate | Perfusion:<br>2% glutaraldehyde, 2% formaldehyde in PB<br>Immersion:<br>Overnight in the same solution | Freezing:<br>glycerol cryoprotection, quick-frozen in a dry ice/ethanol bath, Freeze-substitution | infiltrated with Lowicryl HM-20 | Vibratome (200 µm), ultramicrotome (70 nm) | 5% aqueous uranyl acetate, 1% Reynolds' lead citrate | carbon-coated coverslips | Field Emission Scanning Electron Microscopy |
| Call and Bergles [46] | Immediate | Perfusion:<br>4% PFA<br>Immersion:<br>4% PFA for 6-12 h | Freezing:<br>Sucrose (30%) cryoprotection, frozen in TissueTek,<br>Blocking: | - | cryostat microtome(35–50 µm) | Immunofluorescence:<br>See source | mounted on slides with Aqua Polymount | Confocal Laser-Scanning Microscopy |

| | | | | | | | | |
|---|---|---|---|---|---|---|---|---|
| | | | 5% normal donkey serum, 0.3% Triton X-100 in PBS | | | | | |
| Miguel-Hidalgo et al. [32] | Immediate | - | Freezing: frozen in dry ice | - | cryostat microtome (20 µm) | Immunofluorescence: See source | mounted on slides | Super-Resolution Laser Confocal Microscope |
| Miguel-Hidalgo et al. [32] | 7.25h - 32h | - | Freezing: rapidly frozen | - | cryostat microtome (20 µm) | Immunofluorescence: See source | mounted on slides | Super-Resolution Laser Confocal Microscope |
| Korrell et al. [31] | Immediate | Perfusion: 4% formaldehyde in PBS Immersion: 24-48h in the same solution | Blocking: 2% donkey or goat serum and 0.2% Triton-X100 in PBS | - | vibrating microtome (50 µm) | Immunofluorescence: 6-diamidine-2-phenylindole dihydrochloride | Fluorsave | Confocal Microscopy |
| Arancibia-Cárcamo et al. [30] | Immediate | Perfusion or Immersion: 4% paraformaldehyde in PBS | Blocking and Permeabilisation: 10% horse serum and 0.5% Triton X-100 in PBS | - | vibratome (50 µm) | Immunofluorescence: rabbit anti-$Na_V$1.6, mouse anti-Caspr clone K65/35, anti-rabbit AlexaFluor488, anti-mouse Dy-Light 647 | Dako Fluorescent Mounting Medium | Confocal Microscopy |

| | | | | | | | | |
|---|---|---|---|---|---|---|---|---|
| H01 Shapson-Coe et al. [18] | | Immersion: 2.5% PFA and 2.5% glutaraldehyde in Sodium Cacodylate Buffer, Over night in the same solution | Dehydration<br><br>Plasma-treatmend | 50:50 mixture of PO and 812 Epon resin, 30:70 PO Epon mixture immersion, 100% Epon immersion | Vibratome (300 μm), Ultramicrotome (trimming), automated tape collection Ultramicrotome (30-80nm) | reduced osmium tetroxide-thiocarbohydrazide (TCH)-osmium , en-bloc 2% uranyl acetate, 4% uranyl acetate, 3% lead citrate | carbon-coated Kapton tape, ound or square silicon wafers | Multibeam Scanning Electron Microscopy |

**Table S-I2**: Soma volumes and diameters reported across species and brain regions.

| Source | Species | Brain Area | Layer | Measure | N | Value | Est. Diameter (μm) (based on sphere) |
|---|---|---|---|---|---|---|---|
| Benavides-Piccione et al. [33] | Human | BA17 | 3a | Largest Slice Area ($\mu m^2$) | 55 | mean 154.1<br>range 93.4 - 284.6 | mean 14.0<br>range 10.9 - 19.0 |
| Benavides-Piccione et al. [33] | Human | BA20 | 3a | Largest Slice Area ($\mu m^2$) | 90 | mean 243.1<br>range 163.3 - 406.7 | mean 17.6<br>range 14.4 - 22.8 |
| Benavides-Piccione et al. [33] | Human | BA21 | 3a | Largest Slice Area ($\mu m^2$) | 50 | mean 252.4<br>range 122.1 - 376.8 | mean 17.9<br>range 12.5 - 21.9 |
| Tomasi et al. [19] | Macaque Monkeys | BA46 | 3, 6 | Diameter (μm) of a fitted circle | 339 | range 9 – 15<br>mean 10.6 | range 9 – 15<br>mean 10.6 |
| Tomasi et al. [19] | Macaque Monkeys | BA4 | 3, 6 | Diameter (μm) of a fitted circle | 448 | range 9 – 25<br>mean 13.7 | range 9 – 25<br>mean 13.7 |
| Szocsics et al. [72] | Human | BA4<br>Betz cells | 5 | Largest Slice Area ($\mu m^2$) | 645 | mean range 1148 - 1951 | mean range 38.23 - 49.84 |
| Rivara et al. [34] | Human | BA4<br>Pyramidal cells | 5b | Volume ($\mu m^3$) | 1344 | range 2000 - 18501<br>mean 4274 | range 15.63 - 32.81<br>mean 20.14 |
| Rivara et al. [34] | Human | BA4<br>Betz cells | 5b | Volume ($\mu m^3$) | 2229 | range 10905 - 495000<br>mean 86685 | range 27.51 - 98.15<br>mean 54.9 |
| Brodman [36] | Human | BA4<br>Betz cells | | Width x height ($\mu m^2$) | - | range ? - 53 × 106<br>mean 27-50 x 66-100 | range ? - 74.95<br>mean 42.213 - 70.71 |
| Betz [36] | Human | BA4<br>Betz cells | | Width x height ($\mu m^2$) | - | range ? - 60 × 120 | range ? - 84.85 |

**TableS-I3**: Axon initial segment soma distance and length across species and brain regions.

| Source | Species | Brain Area | Layer | N | Remarks | Distance from Soma (μm) | Length (μm) |
|---|---|---|---|---|---|---|---|
| Fish et al. [69] | Macaque Monkeys | BA 46 | 2-4 | ≥ 500 | pyramidal neurons 3 brains | - | mean 19.7±1.4 |
| Senol et al. [38] | Human | Neocortex | 5-6 | 176 / 45 | pyramidal neurons 5 cases / 5 cases | range 0 - 7.4 mean 1.48 ± 0.37 μm | range 28.6 - 48.5 mean 36.92 ± 0.74 |
| Li et al. [37] | Human | Cerebral cortex | - | > 70 | | - | range 7.8 - 85.8 mean 33.1 |
| Sloper and Powell [39] | Rhesus Monkey | BA 4 BA 3b | 2, 3 | 8 | pyramidal neurons 21 brains | - | range 31 - 54 mean 41.125 |
| Sloper and Powell [39] | Rhesus Monkey | BA 4 BA 3b | 5, 6 | 3 | pyramidal neurons 21 brains | - | range 36 - 49 mean 42.33 |
| Sloper and Powell [39] | Rhesus Monkey | BA 4 BA 3b | 5 | 3 | Betz cells 21 brains | - | range 45 - 50 mean 47.33 |
| Sloper and Powell [39] | Rhesus Monkey | BA 4 BA 3b | 2, 3, 5, 6 | 2 | large stellate cells 21 brains | - | range 22.9 - 23.8 mean 23.35 |
| Höfflin et al. [40] | Mouse organotypic tissue culture | visual cortex | 1-6 | 92 / 178 | pyramidal neurons | range 1.43 - 66.5 mean 13.5 μm ± 12.6 | mean 28.5 μm ± 10.7 range 11.2 - 73.13 |
| Höfflin et al. [40] | Mouse organotypic tissue culture | visual cortex | 1-6 | 48 / 85 | interneurons | range 1.59 - 53.174 mean 14.3 μm ± 10.9 | mean 25.7 μm ± 11.1 range 10.7 - 61.1 |

**TableS-I4**: Cortical axon diameters across species.

| Source | Species | Brain Area | Layer | N | Remarks | Axon Diameter (µm) |
|---|---|---|---|---|---|---|
| Benavides-Piccione et al. [33] | Human | BA17 | 3a | 6 | Neurolucida360<br>average segment diameter | range 0.52 - 0.83 |
| Benavides-Piccione et al. [33] | Human | BA20 | 3a | 25 | Neurolucida360<br>average segment diameter | range 0.51 - 0.81 |
| Benavides-Piccione et al. [33] | Human | BA21 | 3a | 15 | Neurolucida360<br>average segment diameter | range 0.57 - 0.78 |
| Tomasi et al. [19] | Macaque Monkeys | BA46 | 3, 6 | 18 | Neurolucida 7 (coronally cut sections) | range 0.275 - 0.645 |
| Tomasi et al. [19] | Macaque Monkeys | BA4 | 3, 6 | 15 | Neurolucida 7 (coronally cut sections) | range 0.458 - 1.306 |
| Pascual-García et al. [45] | Human | Right temporal lobe, right fronto-temporal lobe, right frontal lobe, right temporo-parietal lobe | 2/3 | 144<br>10 cells | Neurolucida 360 / Explorer<br>Average segment diameter<br>Pyramidal cells close to soma | range 0.134 - 0.607<br>mean 0.29±0.008 s.e.m. |
| Stedehouder et al. [41] | Human | Right parieto-occipital lobe, right temporal lobe, right frontoparietal lobe | 3, 2-5 | 96<br>4 cells | Neurolucida 360 / Explorer<br>Average segment diameter<br>fast-spiking interneurons close to soma | range 0.181 - 0.737<br>mean 0.357±0.1234 |
| Tomasi et al. [19] | Macaque Monkeys | Origin BA9<br>Measured BA9 | GM | 149 | Neurolucida 7 (coronally cut sections) | range 0.28 - 1.1<br>mean 0.48±0.11 |

| | | | | | | |
|---|---|---|---|---|---|---|
| | | towards sulcus principalis | | | | |
| Peters et al. [20] | rhesus monkey | BA 17 | 4Cβ | > 200 | cross-sectional area to equivalent circle diameter<br>Myelinated horizontal fibers | range 0.1 - 2.2<br>mean 0.79±0.275 |

**Table S-I5**: Superficial white matter axon diameters across species.

| Source | Species | Brain Area | Layer | N | Remarks | Axon Diameter (μm) |
|---|---|---|---|---|---|---|
| Zikopoulos and Barbas [24] | Human | Measured SWM below BA32 | SWM<br>(<2mm from the white/gray matter border)<br>Parallel to grey/white boundary | 8000 | Manual measures using ImageJ<br>(Only control case) | mean 0.55±0.04<br>0.0-0.35: 31%±2%<br>0.35-0.7: 49%±4%<br>0.7-1.4: 16%±4%<br>>1.4: 4%±4% |
| Zikopoulos | Human | Measured SWM | SWM | 8000 | Manual measures using | mean 0.44±0.01 |

| | | | | | | |
|---|---|---|---|---|---|---|
| and Barbas [24] | | below BA11 | (<2mm from the white/gray matter border) Parallel to grey/white boundary | | ImageJ (Only control case) | 0.0-0.35: 46%±3%<br>0.35-0.7: 43%±2%<br>0.7-1.4: 10%±1%<br>>1.4: 1%±1% |
| Zikopoulos and Barbas [24] | Human | Measured SWM below BA46 | SWM (<2mm from the white/gray matter border) Parallel to grey/white boundary | 8000 | Manual measures using ImageJ (Only control case) | mean 0.54±0.01<br>0.0-0.35: 36%±6%<br>0.35-0.7: 42%±2%<br>0.7-1.4: 19%±2%<br>>1.4: 4%±2% |
| Tomasi et al. [19] | Macaque Monkeys | Origin BA9 Measured SWM below BA9 | SWM Orthogonal to grey/white boundary | 182 | Neurolucida 7 (coronally cut sections) | range 0.37 - 1.1<br>mean 0.49±0.14 |
| Innocenti et al. [21] | Macaque Monkeys | Origin PEc (BA 5) Measured SWM below BA 4 | SWM Orthogonal to grey/white boundary | 207 | Circular approximation | range 0.29 - 3.79<br>mean 0.95±0.57 |
| Innocenti et al. [21] | Macaque Monkeys | Origin PEc (BA 5) Measured SWM below BA 24 | SWM Orthogonal to grey/white boundary | 203 | Circular approximation | range 0.44 - 1.6<br>mean 0.77±0.21 |
| Innocenti et al. [21] | Macaque Monkeys | Origin PEc (BA 5) Measured SWM below BA 6 | SWM Orthogonal to grey/white boundary | 164 | Circular approximation | range 0.44 - 2.41<br>mean 0.96±0.36 |
| Innocenti et al. [21] | Macaque Monkeys | Origin BA4 Measured SWM below BA 4 | SWM Orthogonal to grey/white boundary | 105 | Neurolucida / Neuroexplorer Axon diameter at surface of section | range 0.2 - 3.9<br>mean 0.79±0.67 |

| | | | | | | |
|---|---|---|---|---|---|---|
| Ruthig et al. [12] | Human | S1 (BA 3), M1 (BA 4) and V2 (BA 18) | SWM (<3mm from the white/gray matter border) Orthogonal and parallel to grey/white boundary | 220431 | Slices, Minor axis of a fitted ellipsoid | range 0.046 - 7.073<br>mean 0.613<br>0.0 - 0.5: 51.1843% (112826)<br>0.5 - 1.5: 44.928% (99035)<br>1.5 - 2.5: 3.3797% (7450)<br>2.5 - 3.5: 0.426% (940)<br>3.5 - 4.5: 0.06714% (148)<br>4.5-5.5: 0.0118% (26)<br>5.5-6.5: 0.0023% (5)<br>6.5-7.5: 0.0005% (1) |

**TableS-I6**: White matter axon diameters across species.

| Source | Species | Brain Area | N | Remarks | Axon Diameter (μm) |
|---|---|---|---|---|---|
| Zikopoulos and Barbas [24] | Human | Measured in deep WM below BA32 | 8000 | Manual measures using ImageJ (Only control case) | range 0.1 - <7.0<br>mean 0.58±0.03<br>0.0-0.35: 29%±5%<br>0.35-0.7: 49%±3%<br>0.7-1.4: 16%±4%<br>>1.4: 7%±2%<br><br>(Likely to contain axons between 2-4μm) |
| Zikopoulos and Barbas [24] | Human | Measured in deep WM below BA11 | 8000 | Manual measures using ImageJ (Only control case) | range 0.1 - <7.0<br>mean 0.49±0.04<br>0.0-0.35: 38%±6%<br>0.35-0.7: 46%±3%<br>0.7-1.4: 13%±4%<br>>1.4: 2%±2%<br><br>(Likely to contain axons between 2-4μm) |
| Zikopoulos and Barbas [24] | Human | Measured in deep WM below BA46 | 8000 | Manual measures using ImageJ (Only control case) | range 0.1 - <7.0<br>mean 0.53±0.03<br>0.0-0.35: 36%±6%<br>0.35-0.7: 42%±4%<br>0.7-1.4: 18%±2%<br>>1.4: 3%±1%<br><br>(Likely to contain axons between 2-4μm) |
| Tomasi et al. [19] | Macaque Monkeys | Origin BA4 Measured CC | 1069 | Circular approximation | range 0.25 - 4.375<br>mode 0.6 |
| Tomasi et al. [19] | Macaque Monkeys | Origin BA9 Measured CC | 634 | Circular approximation | range 0.4 - 2.0<br>mode 0.7 |

| | | | | | |
|---|---|---|---|---|---|
| Tomasi et al. [19] | Macaque Monkeys | Origin MT/V4 Measured CC | 592 | Circular approximation | range 0.25 - 2.875 mode 0.7 |
| Tomasi et al. [19] | Macaque Monkeys | Origin BA9 Measured WM towards CC | 226 | Neurolucida 7 (coronally cut sections) | range 0.27 - 1.25 mean 0.43±0.16 |
| Tomasi et al. [19] | Macaque Monkeys | Origin BA9 Measured WM towards proximal internal capsule | 117 | Neurolucida 7 (coronally cut sections) | range 0.28 - 1.01 mean 0.45±0.14 |
| Tomasi et al. [19] | Macaque Monkeys | Origin BA9 Measured WM towards distal internal capsule | 364 | Neurolucida 7 (coronally cut sections) | range 0.18 - 1.38 mean 0.45±0.12 |
| Innocenti et al. [21] | Macaque Monkeys | Origin PEc (BA 5) Measured SLF | 273 | Circular approximation | range 0.29 - 4.45 mean 1.03±0.75 |
| Innocenti et al. [21] | Macaque Monkeys | Origin BA4 Measured WM towards caudatus | 199 | Neurolucida / Neuroexplorer Axon diameter at surface of section | range 0.2 - 1.7 mean 0.454±0.232 |
| Innocenti et al. [21] | Macaque Monkeys | Origin BA4 Measured WM towards proximal internal capsul | 195 | Neurolucida / Neuroexplorer Axon diameter at surface of section | range 0.2 - 5.2 mean 0.834±0.614 |
| Innocenti et al. [21] | Macaque Monkeys | Origin BA4 Measured WM towards thalamus | 184 | Neurolucida / Neuroexplorer Axon diameter at surface of section | range 0.2 - 2 mean 0.664±0.325 |
| Innocenti et al. [21] | Macaque Monkeys | Origin BA4 Measured WM | 185 | Neurolucida / Neuroexplorer Axon diameter at surface of section | range 0.2 - 6.8 mean 1.384±0.992 |

| | | | | | |
|---|---|---|---|---|---|
| | | towards distal internal capsul | | | |
| Innocenti et al. [21] | Macaque Monkeys | Origin BA9 Measured WM towards caudatus | 80 | Neurolucida / Neuroexplorer Axon diameter at surface of section | range 0.1 - 0.7 mean 0.36±0.11 |
| Innocenti et al. [21] | Macaque Monkeys | Origin BA9 Measured WM towards proximal internal capsul | 103 | Neurolucida / Neuroexplorer Axon diameter at surface of section | range 0.3 - 1.5 mean 0.59±0.25 |
| Innocenti et al. [21] | Macaque Monkeys | Origin BA9 Measured WM towards thalamus | 100 | Neurolucida / Neuroexplorer Axon diameter at surface of section | range 0.2 - 1.0 mean 0.33±0.13 |
| Innocenti et al. [21] | Macaque Monkeys | Origin BA9 Measured WM towards distal internal capsul | 100 | Neurolucida / Neuroexplorer Axon diameter at surface of section | range 0.2 - 1.6 mean 0.54±0.27 |
| Firmin et al. [13] | Rhesus Monkey | Medullary Pyramid | 4172 | Slcies, Minimal inner diameter | range 0.04 - 9.48 mean 0.91 **Outer** diameter distribution 0.0-0.5: 14% 0.5-1.0: 38% >1.0: 48% |
| Graf von Keyserlingk and Schramm [22] | Human | Medullary Pyramid | 2400 | MOP AM 03 (Kontron) | range <0.3 - >10 mode ~0.5 **Outer** diameter distribution 0 - 4: 87.9% 4 - 10: 10.77% >10: 1.4% |

| | | | | | |
|---|---|---|---|---|---|
| Ruthig et al. [12] | Human | CC (rostrum, genu, anterior midbody, midbody, isthmus, splenium) | 16313<br>2 | Slices,<br>Minor axis of a fitted ellipsoid | range 0.066 - 7.48<br>mean 0.764<br>0.0 - 0.5: 29.925% (48817)<br>0.5 - 1.5: 62.928% (102656)<br>1.5 - 2.5: 5.9173% (9653)<br>2.5 - 3.5: 0.9655% (1575)<br>3.5 - 4.5: 0.20781% (339)<br>4.5-5.5: 0.0448% (73)<br>5.5-6.5: 0.0104% (17)<br>6.5-7.5: 0.0018% (3) |
| Andersson et al. [68] | Vervet monkey | CC (splenium) | 54 | Mean axon diameter (length: 124-170 μm) | range 2.05 - 3.77 |
| Liewald et al. [23] | Human | Sup. longitudinal fascicle | 1479 | Slices, longest diameter perpendicular to longest axis | range 0.19 - 4.79<br>mean 0.74<br>largest axon found: 9 |
| Liewald et al. [23] | Human | Uncinate/inferior occipitofr. fasc. | 2930 | Slices, longest diameter perpendicular to longest axis | range 0.16 - 4.42<br>mean 0.52 |
| Liewald et al. [23] | Human | CC (genu, truncus, splenium) | 2362 | Slices, longest diameter perpendicular to longest axis | range 0.17 - 5.13<br>mean 0.69 |

**Table S-I7**: Axon bend radii across species and brain regions.

| Source | Species | Brain Area | Type | Bend Radius (µm) |
|---|---|---|---|---|
| Cerebral Cortex Volume 10, Chapter 6, Fig 12 [50] | Primate | V1,V2 | Myelinated elbow-shaped turns in layer 1 | C, D: 40.6<br>E: 25.8<br>F: 23.4 |
| Yamashita et al. [42] | Macaque Monkey | BA4a-3a | Likely myelinated axon collateral from L4 PC bending in gray matter | Fig 3: 255.3 |
| DeFelipe et al. [43] | cynomolgus, pigtailed | BA1-2 | Likely myelinated axon collateral from L2/3 PC bending in gray matter | Fig 9: 461.2 |
| Yamashita et al. [42] | Macaque Monkey | BA4b-3a | Myelinated corticocortical axon from L3 PC bending in WM | Fig 4: 95.5<br>Fig 5: 71.7 |
| Rockland and Virga [49] | Fascicularis Monkey | V1, V2 | Likely myelinated corticocortical axon bending in WM | Fig 7: 75.3<br>Fig 9-20: 38.4<br>Fig 9-19: 71.3<br>Fig 10-18: 99.0, 30.6<br>Fig 10-17: 76.8, 3.1, 61.9<br>Fig 13-33: 82.7 |
| DeFelipe et al. [43] | cynomolgus, pigtailed | BA2<br>BA3a<br>BA4<br>BA2 | Myelinated corticocortical axon bending in WM | Fig 4: 9.5<br>Fig 6: 9.7<br>Fig 8: 27.2<br>Fig 10: 7.7, 10.5 |
| Yamashita et al. [42] | Macaque Monkey | BA4a-3a-3b | Myelinated U-fiber from L3 PC | Fig 5: 1596.9 |
| DeFelipe et al. [43] | cynomolgus, pigtailed | BA3b-3a | Likely myelinated U-fiber | Fig 12: 592.1 |
| Li et al. [55] | Macaque Monkey | M1 | L3 and L5 PCs smoothly project into deep WM<br>(2, 4, 6, 7, 8, 9, 10,11, 12, 13, 14, 16, 17, 19, 21, 22, 25, | range 788.7 - 1560.8<br>mean 1217.3±210.4 |

| | | | | |
|---|---|---|---|---|
| | | | 26) | |
| Mortazavi et al. [51] | Rhesus Monkeys | BA4 | Myelinated axons bending in deep WM | range: 13 - 17<br>median: 14 |

**Table S-I8**: Axon internode lengths across species and brain regions.

| **Source** | **Species** | **Area** | **N** | **$R^2$** | **Remark** | **Axon Diameter (D) (µm) to Internode Length (L) (µm)** |
|---|---|---|---|---|---|---|
| Hess and Young [70] | Rabbit | Spinal Cord | - | - | outer diameter | L = 109.5 D + 64.0<br>range D: 3 - 15<br>range L: 380 - 1720 |
| Ford et al. [27] | Rodent | Auditory Brainstem SBC | - | - | outer diameter<br>> 500 µm from the medial somatodendritic region of the MSO | L = 122 D<br>mean L: 164.8±12.4<br>mean D: 1.35±0.03 |
| Ford et al. [27] | Rodent | Auditory Brainstem GBClat | - | - | outer diameter<br>> 700 mm from the terminals | L = 64.9±3.1 D<br>mean L: 195.7±8.5<br>mean D: 3.21±0.05 |
| Ford et al. [27] | Rodent | Auditory Brainstem GBCmed | - | - | outer diameter<br>> 700 mm from the terminals | L = 99.6±5.7 D<br>mean L: 230.4±11.1<br>mean D: 2.40±0.05 |
| Ibrahim et al. [26] | Rodent | anterior medullary velum IVth tract | 145 | 0.2813 | outer diameter | L = 42.023 D + 85.317 |
| Ibrahim et al. [26] | Rodent | anterior medullary velum rostra regio | 129 | 0.192 | outer diameter | L = 24.707 D + 124.46 |
| Ibrahim et al. [26] | Rodent | anterior medullary velum caudal region | 93 | 0.0086 | outer diameter | L = 6.005 D + 244.45 |
| Ibrahim et al. [26] | Rodent | anterior medullary velum | > 326 | 0.2852 | outer diameter | L = 30.475 D + 117.52 |
| Butt et al. [25] | Rodent | anterior medullary velum | - | 0.317 | outer diameter | L = 111.17 Ln(D) + 97.316 |
| Andersson et al. | Vervet | CC (splenium) | 6 | - | | range L: 250.0 - 423.1 |

| | | | | | | |
|---|---|---|---|---|---|---|
| [68] | monkey | | | | | range D: ~2 - 5 |
| Pascual-García et al. [45] | Human | Right temporal lobe, right fronto-temporal lobe, right frontal lobe, right temporo-parietal lobe Layer 2/3 | 20 segments | - | Pyramidal cells close to soma | mean L: 49.37 ± 12.69 |
| Stedehouder et al. [41] | Human | Right parieto-occipital lobe, right temporal lobe, right frontoparietal lobe | 36 segments | - | fast-spiking interneurons close to the soma | mean L: 44.7 ± 4.9 |
| Stedehouder et al. [71] | Human | right temporo-occipital lobe, right temporal lobe | 194<br>10 cells | | fast-spiking interneurons (basket cells, double-bouquet cell) | range L: ~10 - >80<br>mean L: 33.0 ± 1.4 |
| Micheva et al. [44] | Mouse | S1 Layer 2/3 and 4 | 23 | - | GABA axons | mean L: 22.54 ± 2.1 |
| Micheva et al. [44] | Mouse | S1 Layer 2/3 and 4 | 32 | - | non-GABA axons | mean L: 29.29 ± 2.02 |
| Call and Bergles [46] | Mouse | Neurons with axons in L1 of S1 | 119 | - | GABAergic neurons (parvalbumin, somatostatin), layer VI corticothalamic, layer VIb subplate; layer Va/Vb pyramidal | range L 16.9 - 137.8<br>mean L 57.2 |

**Table S-I9**: Node of Ranvier lengths across species and brain regions.

| Source | Species | Area | N | Remark | Node of Ranvier Length (μm) |
|---|---|---|---|---|---|
| Miguel-Hidalgo et al. [32] | Human | PFC WM | - | 200 μm from the WM / gray matter boundary<br>11 brains | mean 2.5 |
| Miguel-Hidalgo et al. [32] | Rat | PFC WM | - | 6 brains | mean 1.26 |
| Korrell et al. [31] | Mouse | CC | 116 | IUE control<br>6 brains | range 0.1 - 4.35<br>mean 1.61 ± 0.07 |
| Korrell et al. [31] | Mouse | CC | 46 | LVI SNAP25 control<br>3 brains | range 0.69 - 3.5<br>mean 1.47 ± 0.06 |
| Korrell et al. [31] | Mouse | CC | 33 | LV SNAP25 control<br>3 brains | range 0.81 - 2.85<br>mean 1.56 ± 0.09 |
| Arancibia-Cárcamo et al. [30] | Rat | Motor Cortex L5 | 158 | | range 0.43 - 3.72<br>mean 1.50 ± 0.05 |
